# NeuroPAL: A Neuronal Polychromatic Atlas of Landmarks for Whole-Brain Imaging in *C. elegans*

**DOI:** 10.1101/676312

**Authors:** Eviatar Yemini, Albert Lin, Amin Nejatbakhsh, Erdem Varol, Ruoxi Sun, Gonzalo E. Mena, Aravinthan D.T. Samuel, Liam Paninski, Vivek Venkatachalam, Oliver Hobert

**Affiliations:** Department of Biological Sciences, Columbia University, New York; Howard Hughes Medical Institute; Department of Physics, Center for Brain Science, Harvard University, Cambridge, MA; Departments of Statistics and Neuroscience, Grossman Center for the Statistics of Mind, Center for Theoretical Neuroscience, Zuckerman Institute, Columbia University, New York; Department of Statistics and Data Science Initiative, Harvard University, Cambridge, MA; Department of Physics, Northeastern University, Boston, MA

## Abstract

Comprehensively resolving single neurons and their cellular identities from whole-brain fluorescent images is a major challenge. We achieve this in *C. elegans* through the engineering and use of a multicolor transgene called NeuroPAL (a **Neuro**nal **P**olychromatic **A**tlas of **L**andmarks). NeuroPAL worms share a stereotypical multicolor fluorescence map for the entire hermaphrodite nervous system that allows comprehensive determination of neuronal identities. Neurons labeled with NeuroPAL do not exhibit fluorescence in the green, cyan, or yellow emission channels, allowing the transgene to be used with numerous reporters of gene expression or neuronal dynamics. Here we showcase three studies that leverage NeuroPAL for nervous-system-wide neuronal identification. First, we determine the brainwide expression patterns of all metabotropic receptors for acetylcholine, GABA, and glutamate, completing a map of this communication network. Second, we uncover novel changes in cell fate caused by transcription factor mutations. Third, we record brainwide activity in response to attractive and repulsive chemosensory cues, characterizing multimodal coding and novel neuronal asymmetries for these stimuli. We present a software package that enables semi-automated determination of all neuronal identities based on color and positional information. The NeuroPAL framework and software provide a means to design landmark atlases for other tissues and organisms. In conclusion, we expect NeuroPAL to serve as an invaluable tool for gene expression analysis, neuronal fate studies, and for mapping whole-brain activity patterns.

## INTRODUCTION

Whole-brain imaging and molecular profiling are widely used to study nervous system development and brain function (Ahrens and Engert, 2015; Jones et al., 2009; Lichtman and Denk, 2011; Zeng and Sanes, 2017). One limitation in interpreting whole-brain images is the difficulty of assigning unique identities to every neuron in a volume of densely packed and similarly labeled cells. Identifying neurons is a challenge even in small nervous systems like that of the nematode *C. elegans*. While it is possible to perform multineuronal functional imaging with single-cell resolution in *C. elegans* (Kato et al., 2015; Kotera et al., 2016; Nguyen et al., 2016; Venkatachalam et al., 2016), identifying neurons remains laborious and uncertain, requiring substantial expertise, even in light of recent advances (Bubnis et al., 2019; Toyoshima et al., 2020). The approach of using separate and sparsely-labeled landmark strains is often helpful, but not scalable. Also, many neurons lack well-established reporters, and it is not always possible to cross-validate every neuron of interest in a densely-labeled volume, even with a suitable landmark strain. Moreover, while the *C. elegans* nervous system is widely regarded as stereotyped, this stereotypy does not extend to the relative positions of cell bodies within ganglia (White et al., 1986). An invariant color map of all neurons is thus needed to achieve comprehensive cell identification. Here, we leveraged the small size of the worm nervous system and its powerful genetics to develop a new method to identify all neurons in a whole-brain image with a single reagent. We describe the development of a transgene that we call NeuroPAL (**Neuro**nal **P**olychromatic **A**tlas of **L**andmarks).

The NeuroPAL transgene contains a combination of 41 selectively overlapping neuron-specific reporters, each of which expresses a subset of four distinguishably-colored fluorophores. The NeuroPAL combination of reporters and colors generates a comprehensive color-coded atlas for the entire hermaphrodite nervous system. Our approach is fundamentally different from previously described “Brainbow” approaches (Livet et al., 2007; Richier and Salecker, 2015; Weissman and Pan, 2015). In “Brainbow”, multicolor labeling of the nervous system occurs when each neuron randomly expresses a subset of fluorophores. In NeuroPAL, each neuron expresses a stereotyped combination of fluorophores. NeuroPAL yields an invariant color map across individuals, where every neuron is uniquely identified by its color and position. We engineered NeuroPAL to be compatible with widely-used reporters for gene expression and neuronal activity. None of the NeuroPAL fluorophores emit in the spectral bands of green, cyan, or yellow fluorescent proteins. Thus, NeuroPAL can be co-expressed with numerous markers – GFP, CFP, YFP, mNeonGreen, or reporters of neuronal dynamics like GCaMP – without affecting its color map.

We demonstrate the versatility of NeuroPAL in studies of gene expression patterns, cell fate, and whole-brain activity imaging. First, we mapped with single neuron resolution the complete gene expression patterns of all metabotropic receptors for common neurotransmitters (acetylcholine, GABA, and glutamate) encoded in the *C. elegans* genome. This map delineates a significant portion of the second-messenger communication system and, when combined with previous expression pattern analysis (Bamber et al., 1999; Beg and Jorgensen, 2003; Gendrel et al., 2016; Jobson et al., 2015), completes the entire GABA communication network in *C. elegans*. Our analysis of this network suggests considerable extrasynaptic GABA reception, primarily in sensory neurons. Second, we analyzed neuronal fate defects caused by mutations in the highly-conserved transcription factors (TFs) EOR-1/PLZF and PAG-3/Gfi. Third, we measured the complete circuit-level responses to a gustatory repellent (a high concentration of NaCl) and two olfactory attractants (2-butanone and 2,3-pentanedione). We observed stimulus-specific brainwide response patterns and uncovered novel left/right asymmetric responses in sensory and interneurons.

To facilitate the use of NeuroPAL, we provide an open-source software package that enables semi-automated identification of all neurons in whole-brain images. To extend the NeuroPAL technique for general use, we also provide software that chooses reporter-fluorophore assignments for other tissue and organisms. In conclusion, NeuroPAL now allows the *C. elegans* community to easily identify all neurons in whole-brain images for diverse applications.

## RESULTS

### Constructing the color palette for comprehensive landmarks

The *C. elegans* nervous system contains 302 neurons (organized into 118 different classes) distributed among 11 ganglia throughout the body (Sulston, 1983; White et al., 1986). The set of neurons in each ganglion are the same from animal to animal, but the relative location of cell bodies within each ganglion are variable. The largest ganglia contain around 30 neurons. We reasoned that roughly 30 unique colors would be needed to reliably identify all neurons in each ganglion, and thus all neurons in the nervous system. Three spectrally-distinct fluorophores, distinguishable at four or more different levels (high, medium, low, and undetectable), yield at least 64 different colors. Thus, three carefully chosen fluorophores should be enough to landmark the entire *C. elegans* nervous system (**Figure 1**).

**Figure 1.**
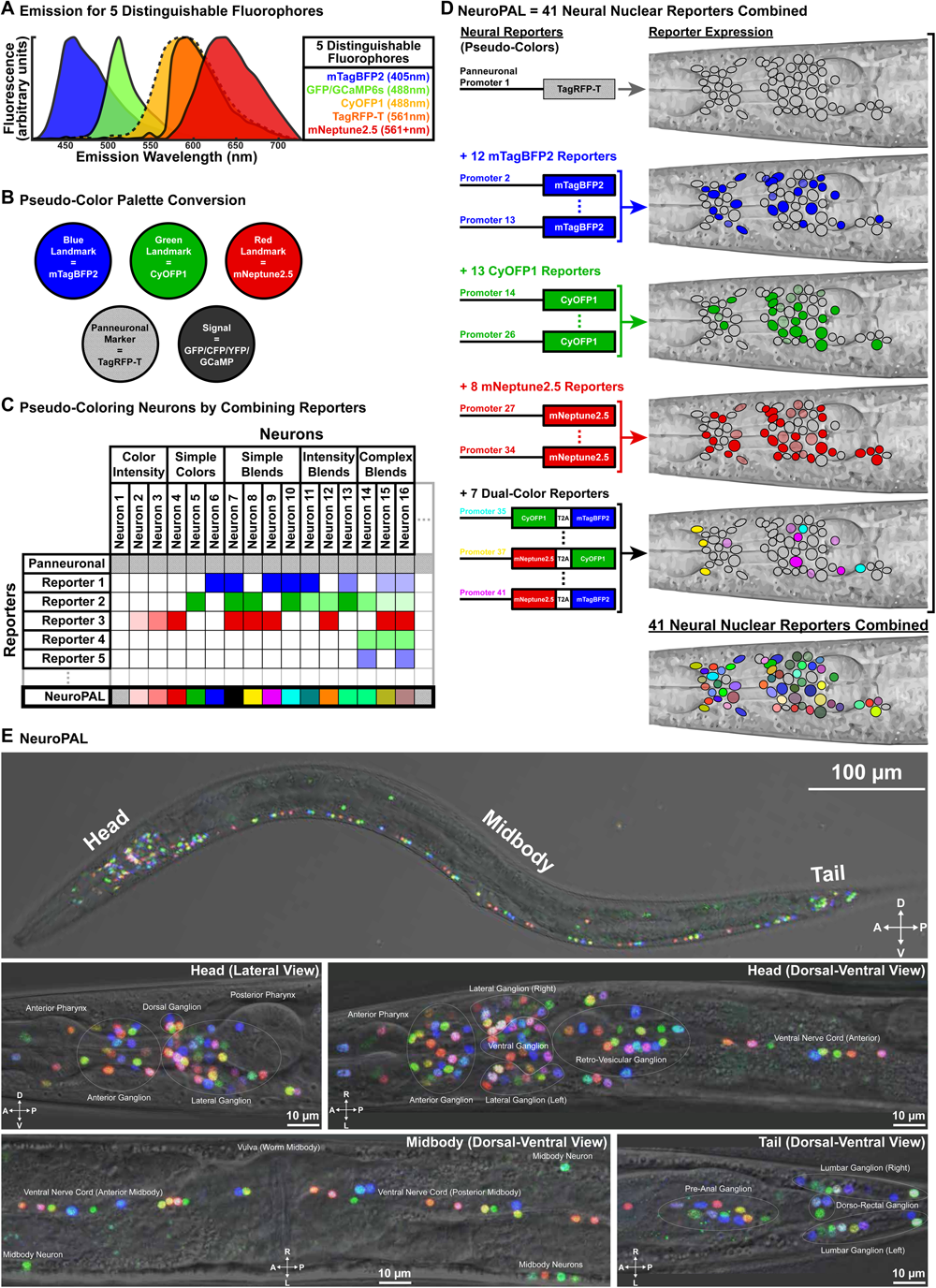
NeuroPAL method and images. (A) The emission for five distinguishable fluorophores. Each fluorophore’s excitation wavelength is listed in parentheses. (B) Fluorophores are converted into pseudo colors to construct a primary color palette. Three fluorophores are designated as landmarks and pseudo colored to construct an RGB color palette: mNeptune2.5 is pseudo-colored red, CyOFP1 is pseudo-colored green, and mTagBFP2 is pseudo-colored blue. The fluorophore TagRFP-T is used as a panneuronal marker. The fluorophores GFP/CFP/YFP/GCaMP6s are reserved for reporters of gene expression or neural activity. TagRFP-T and GFP/CFP/YFP/GCaMP6s are visualized separately from the RGB landmarks to avoid confusion. They can be assigned any pseudo color. (C) An example of how to stably pseudo color neurons, across animals. A set of reporters (rows), with stable neuronal expression (columns), are used to drive the fluorophores (table elements). NeuroPAL colors (last row) result from the combined patterns of reporter-fluorophore expression. The panneuronal reporter is expressed in all neurons. The remaining reporters have differential neuronal expression patterns and are used to drive the pseudo-colored landmark fluorophores. Combinations of these differentially expressed reporters assign stable and distinguishable pseudo colors to neurons. For example, Reporter 3 drives the red landmark fluorophore (mNeptune2.5), to color Neurons 2-4 with distinguishable red intensities and contributes to the blended coloring of several other neurons. In contrast, Neuron 1 does not express any of the landmark fluorophores but is still marked by the panneuronal reporter. (D) NeuroPAL scales this concept to 41 reporters that, in combination, disambiguate every neuron in *C. elegans* and, thus, generate, a single stereotyped color map across all animals (see **Table S1** for data). For convenience, seven of the NeuroPAL reporters use a self-cleaving peptide sequence (T2A) to simultaneously drive expression of two different colors. (E) Young adult NeuroPAL worms (*otIs669*) have a deterministic color map that remains identical across all animals (see **Video S1-S2** for 3D images). Each neuron is distinguishable from its neighbors via color. All worm ganglia are shown. *\* All images may employ histogram adjustments to improve visibility. Images without a transmitted light channel (e*.*g*., *Nomarski) may further be adjusted with a gamma of ∼0*.*5 to improve visibility on the dark background*.

We wanted our landmark reagent to be usable in animals that co-express transgenic reporters for gene expression or neuronal dynamics. The most popular fluorescent reporters include CFP, GFP/GCaMP, and YFP. We did not want our landmark fluorophores to contaminate emission signals from any of these reporters and vice-versa. Therefore, we sought fluorophores with unique excitation/emission profiles that also left free the cyan, green, and yellow emission bands. We tested a wide variety of fluorophores and found mTagBFP2, CyOFP1, mNeptune2.5, and TagRFP-T to be the best available candidates (**Figure 1A and S1**) (Chu et al., 2014; Chu et al., 2016; Shaner et al., 2008; Subach et al., 2011). By pseudo-coloring these fluorophores blue, green, red, and white respectively, their combinations generate RGB pseudo-colors (**Figure 1B-1C**).

Given the resolution limitations of light microscopy, we did not want fluorescence signals from neighboring cells to contaminate one another. To minimize spatial overlap in fluorescence emission, we localized fluorophore expression to cell nuclei via nuclear-localization sequences or histone tagging. We assigned one fluorophore, TagRFP-T, to act as a panneuronal label. To minimize differential and variable expression levels associated with many known panneuronal drivers (Stefanakis et al., 2015), we constructed a synthetic ultra-panneuronal (UPN) driver by fusing the *cis*-regulatory elements of four different panneuronally expressed genes (**Table S1**). The UPN driver delivered bright, nearly uniform expression of TagRFP-T throughout the nervous system.

### Empirical assembly of NeuroPAL, a transgene combining neuronal landmarks

Next, we sought to differentially express the remaining three fluorophores (mTagBFP2, CyOFP1, and mNeptune2.5) to enable unique cellular identification (**Figure 1C-1D**). We used two criteria to build a stereotyped cellular color map for unambiguous and comprehensive assignment of identity: a) each neuron should express a stable amount of each fluorophore, across animals; and b) nearby cells in each ganglion should express visually distinguishable amounts of the fluorophores.

We began with a candidate list of 133 published neuronal reporters known to have differential gene expression patterns. This list included both broad and narrowly expressed reporters (**Table S1**). We validated and completed identification of the expression of these candidates by co-expressing them with well-characterized cell-identity reporters. Candidates with variable or weak expression were dropped from further consideration. We then proceeded empirically and iteratively to build a single transgene for comprehensive neuronal identification. We started with a small set of broadly-expressed reporters that spanned most of the nervous system. We gradually expanded this initial set, adding and changing reporters so as to target progressively smaller subsets of neurons that needed distinguishable colors. In each iteration, we assessed which neurons could and could not be identified based on color and position. We repeated these steps by trial and error until we found a suitable transgene that colored all neurons distinguishably from every neighboring cell. The final transgene, composed of 41 different reporter-fluorophore fusions, allowed us to unambiguously assign identities to every neuron in *C. elegans* based on a stereotyped color map (**Figure 1E; Video S1-S2; Table S1; NeuroPAL Manuals: https://www.hobertlab.org/neuropal/**). We called this transgene NeuroPAL.

### Neuronal color verification and phenotypic assessment of NeuroPAL strains

We integrated the extrachromosomal NeuroPAL transgene into the genome, outcrossed the brightest integrants (*otls669, otls670*, and *otls696*) eight times, and confirmed that these transgenes exhibited stable expression for more than 100 generations. The color scheme of the NeuroPAL strains matched our expectations based on the combination of reporter-fluorophore fusions used in their construction. We verified the identity of each neuron by crossing the NeuroPAL integrants to 25 different GFP reporter lines with well-defined expression patterns (**Table S1**). The position, color, and identity of all neurons were verified using predominantly two or more GFP reporter lines and multiple NeuroPAL integrants. We found that the NeuroPAL expression pattern was stable, robust, and stereotyped throughout the nervous system over hundreds of scored animals (**see NeuroPAL Manuals: https://www.hobertlab.org/neuropal/)**. A minor exception were four neuron classes that exhibited variable brightness (AVL, RIM, RIS, and PVW). This minor variability did not affect our ability to comprehensively identify all neurons.

We assessed the general health of our NeuroPAL integrants (**Figure S2; Table S2**). All NeuroPAL integrants were able to be revived from frozen stock and generate progeny from either hermaphrodite or male parents. Thus, every integrant can be combined with other transgenic reporter lines using genetic crosses. We tested all NeuroPAL integrants with standard assays including brood size, growth, morphology, locomotion, and chemotaxis. The brightest integrant was *otls669*. The integrant with locomotion and chemotactic behavior closest to wild type, *otls670*, is less bright but perhaps more suitable for behavioral analysis and calcium imaging. All NeuroPAL integrants are available at the *Caenorhabditis* Genetics Center (CGC).

### Variability in neuronal cell body positions

The *C. elegans* hermaphrodite nervous system is widely regarded as stereotyped. However, the original electron-micrograph reconstructions (White et al., 1986) and subsequent analysis (Toyoshima et al., 2020) have reported variability in the positions of individual cell bodies within each ganglion of the nervous system. Variability in the position of individual cell bodies makes it impossible to assign cell identities based on relative position alone (**Figure 2A-2B**), underscoring the need for a reagent like NeuroPAL that disambiguates these identities based on genetic-expression factors. Nevertheless, a probabilistic map of neuronal positions would be useful in many studies. To construct this probabilistic map, we globally aligned neurons in the head and tail of 10 young-adult NeuroPAL hermaphrodites (*otls669*) of identical age, and measured the spatial coordinates of every neuron (**Table S3; Text S1**). This map revealed different positional variability across neuron types (**Figure 2C-2D**).

**Figure 2.**
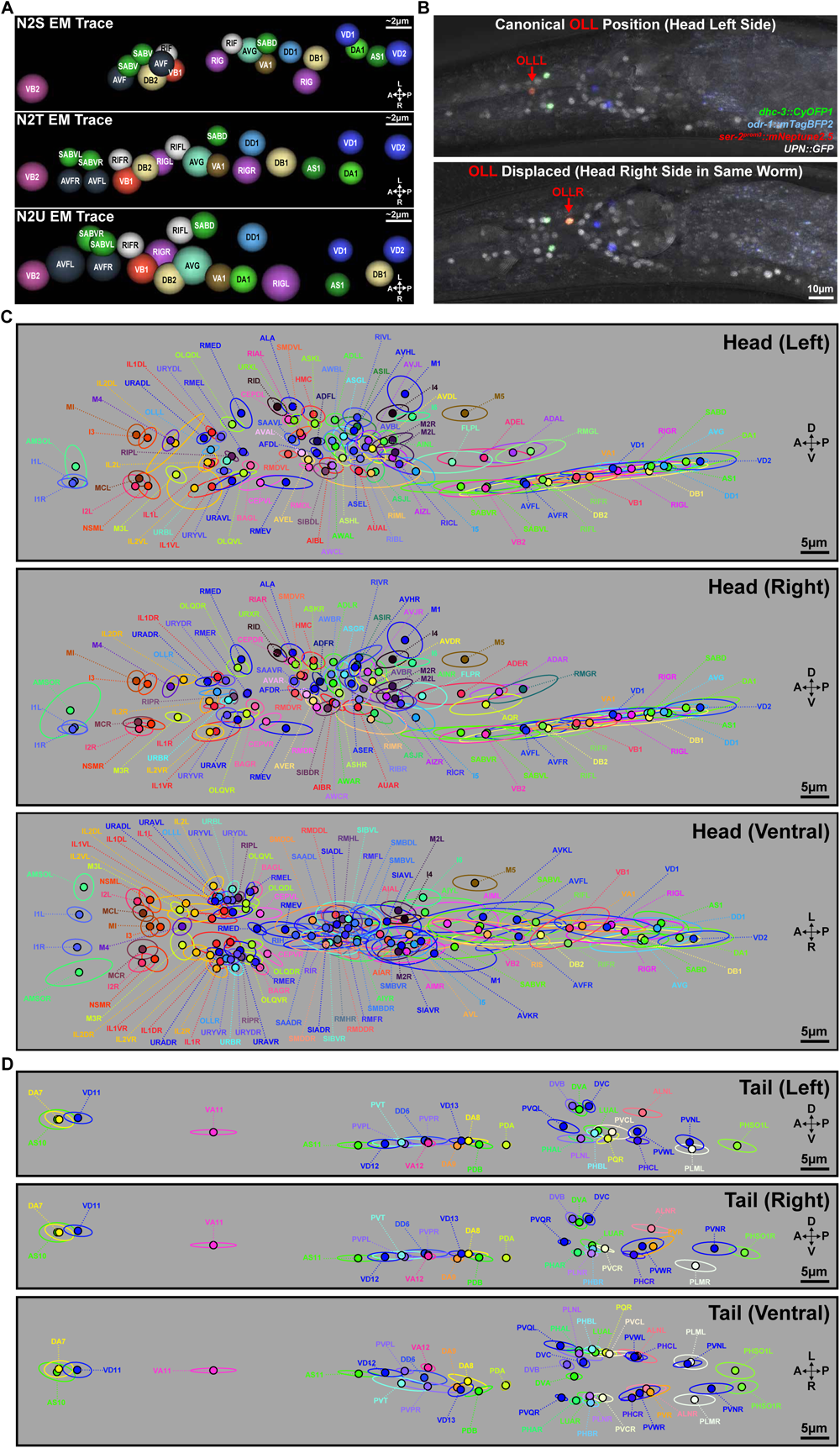
Neuron locations and their positional variability. (A) Neuron locations and variability, in the retrovesicular ganglion, taken from electron micrographs of three adult hermaphrodites N2S, N2T, and N2U (Hall and Altun, 2007; White et al., 1986). (B) An example of substantial positional variability. The OLL left (OLLL) and right (OLLR) neurons, within a single animal, should share equivalent positions. Instead they show substantial anterior-posterior displacement relative to each other. The transgenic reporters and their pseudo colors are noted on the figure. (C,D) Canonical neuron locations (filled circles displaying the NeuroPAL colors) and their positional variability (encircling ellipses with matching colors) for all ganglia, as determined by NeuroPAL (*otIs669*) (see **Table S3** for data). Positional variability is shown as the 50% contour for neuronal location (measured as a Gaussian density distribution), sliced within a 2D plane (**Text S1**). We show both the left-right and dorsal-ventral planes to provide a 3D estimation of positional variability. (C) Left, right, and ventral views of the head neuron positions. OLLR exhibits over twice the positional variability of OLLL in its anterior-posterior axis, echoing the displacement seen with the non-NeuroPAL transgene in panel B. (D) Left, right, and ventral views of the tail neuron positions.

### Expression maps for all metabotropic neurotransmitter receptors

To showcase NeuroPAL as a tool for expression-pattern analysis, we focused on neurotransmitter-signaling maps. Maps of neurotransmitter expression have been determined for the entire *C. elegans* nervous system (Gendrel et al., 2016; Pereira et al., 2015; Serrano-Saiz et al., 2013). Postsynaptic neurons receive neurotransmitter signals by either ionotropic or metabotropic receptors. However, the map of neurotransmitter receptor expression remains largely unknown, leaving their communication networks incomplete. We used NeuroPAL to map the complete expression of all metabotropic neurotransmitter receptors using primarily fosmid-based reporters. The worm genome predicts three cholinergic metabotropic receptors (*gar-1, gar-2*, and *gar-3*), three glutamatergic metabotropic receptors (*mgl-1, mgl-2*, and *mgl-3*), and two GABAergic metabotropic receptors (*gbb-1, gbb-2*) (Hobert, 2013)(**Figure 3 and 4A**). Strikingly, these receptors are expressed in 97% of all neurons: 70% express GARs, 54% express MGLs, and 89% express GBBs (**Figure 4C; Table S4**). Previous work identified all presynaptic GABAergic neurons as well as all postsynaptic ionotropic GABA_A_ receptor-expressing neurons (Bamber et al., 1999; Beg and Jorgensen, 2003; Gendrel et al., 2016; Jobson et al., 2015). Thus, our GBB map (the identity of every GABA_B_ receptor-expressing neuron) has completed the GABA communication network in *C. elegans* (**Figure 4B**).

**Figure 3.**
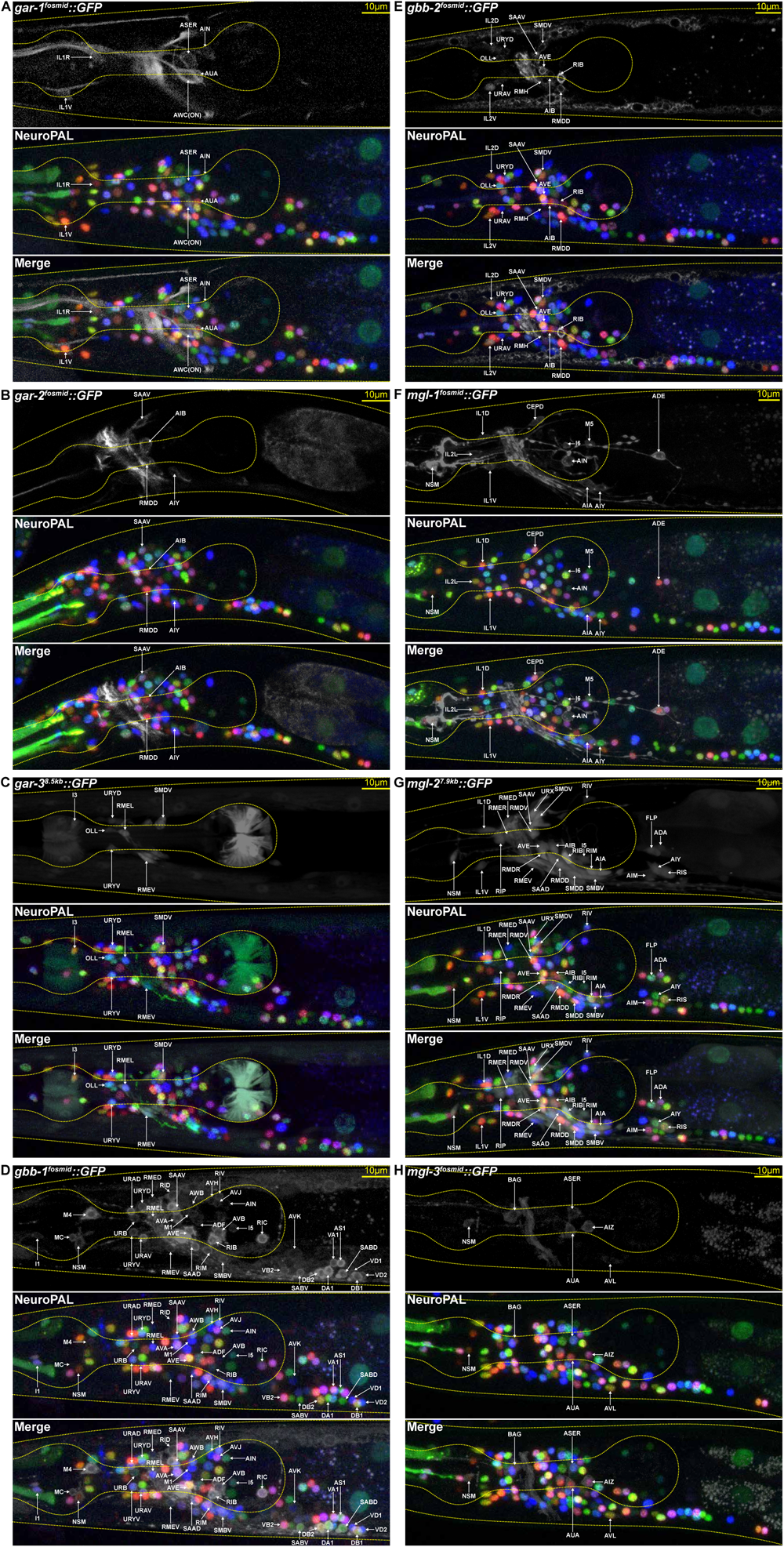
Expression for all metabotropic neurotransmitter receptors. (A-H) NeuroPAL is used to identify the GFP expression patterns for all metabotropic neurotransmitter receptors (**Table S4**): the three acetylcholine receptors (A) GAR-1, (B) GAR-2, and (C) GAR-3; the two GABA receptors (D) GBB-1 and (E) GBB-2; and the three glutamate receptors (F) MGL-1, (G) MGL-2, and (H) MGL-3.

**Figure 4.**
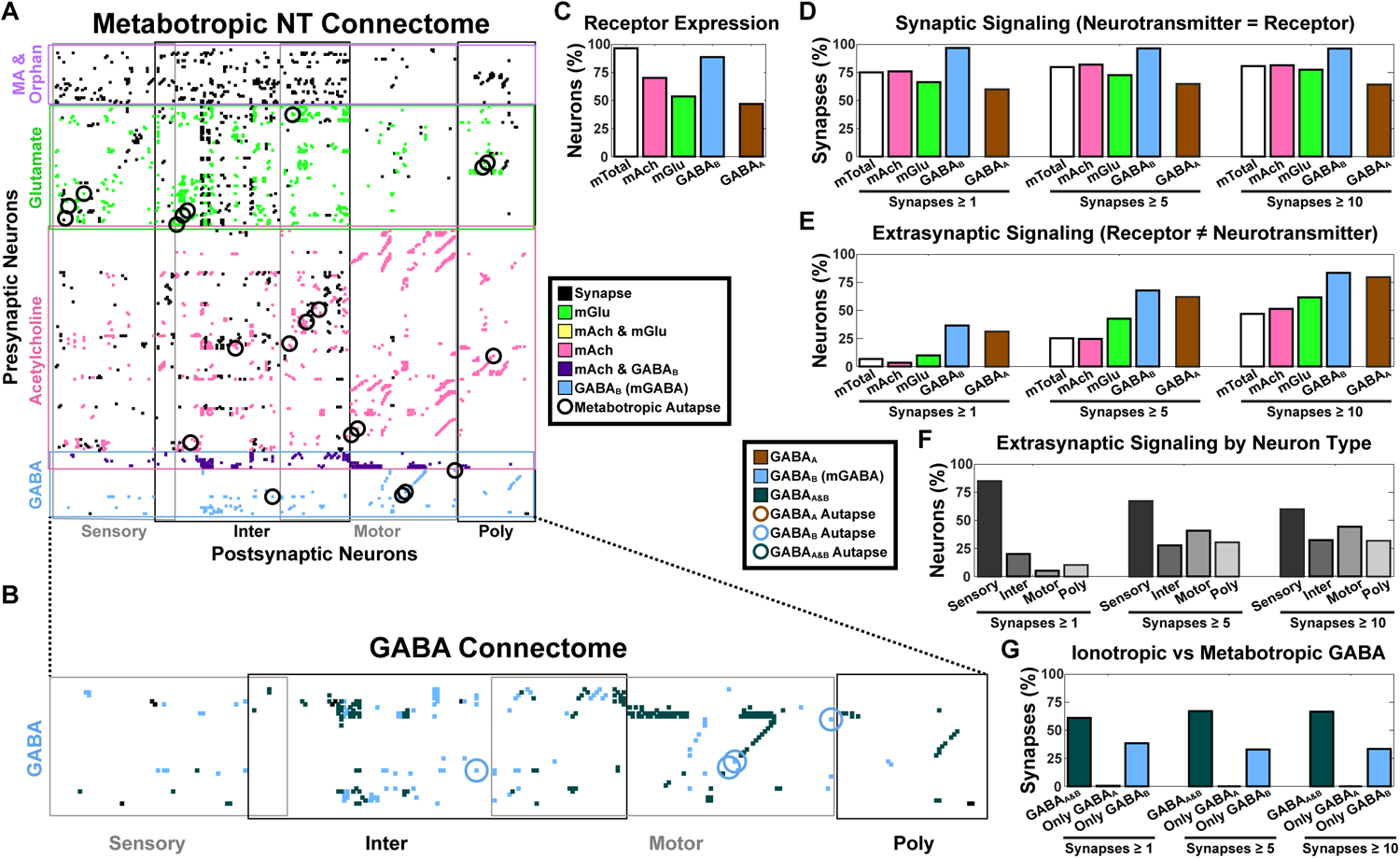
Metabotropic neurotransmitter communication network. Metabotropic receptor abbreviations: acetylcholine=mAch, glutamate=mGlu, GABA=GABA_B_, all=mTotal. The ionotropic GABA receptor is abbreviated GABA_A_. See **Table S4** for data. (A) Expression for the metabotropic neurotransmitter receptors incorporated into the existing, anatomically-defined connectome. Rows are presynaptic neurons (organized by neurotransmitter). Columns are postsynaptic neurons (organized by neuron type). Cognate synaptic connections (where the presynaptic neurotransmitter matches the postsynaptic metabotropic receptor) are marked by a colored dot (see legend for neurotransmitter coloring). All other synaptic connections are marked by a black dot. Metabotropic autapses with cognate self-connectivity are circled. (B) The complete GABA communication network: all presynaptic GABA neurons and their corresponding postsynaptic ionotropic GABA_A_ and metabotropic GABA_B_ expressing neurons. Synaptic connections are marked by a colored dot to indicate postsynaptic GABA_A_ and/or GABA_B_ receptors (see legend for receptor-type coloring). GABA autapses are circled. All GABAergic autapses express GABA_B_ receptors and none express GABA_A_ receptors. (C) Metabotropic (and ionotropic GABA_A_) receptor expression as a percentage of all neurons. Metabotropic receptors are expressed in almost all neurons. (D) Synaptic connections with cognate neurotransmitter receptors as a percentage of the total synapses associated with each neurotransmitter. Metabotropic communication is extensive and these percentages are robust when removing weak synaptic connections by thresholding for at least 1, 5, or 10 synapses. *\* A polyadic synapse that connects AVF (a neuron with no means of releasing its GABA) onto AIM and several cognate GABA*_*B*_ *expressing neurons, reduces our count of GABA synaptic signaling to just under 100%*. (E) Neurons expressing a neurotransmitter receptor that have no presynaptic partners expressing their cognate neurotransmitter. The absence of presynaptic GABAergic partners for over 31% of the GABA_A_ and 37% of the GABA_B_ receptor-expressing neurons suggests substantial extrasynaptic GABA signaling. For each neurotransmitter, we show the percentage of neurons lacking a cognate presynaptic partner relative to all neurons expressing the neurotransmitter receptor. Removing weak synaptic connections increases the potential for extrasynaptic signaling. (F) The types of neurons expressing a neurotransmitter receptor that have no presynaptic partners expressing their cognate neurotransmitter (suggesting extrasynaptic signaling). We show the neuron type percentages. Sensory neurons represent the majority and are robust against removing weak synaptic connections. *\* Neurons are often categorized as multiple types and thus the percentages exceed 100%*. (G) Ionotropic versus metabotropic GABA communication. Over 60% of GABA connections share both GABA_A_ and GABA_B_ receptors at their postsynaptic sites. The remaining nearly 40% of connections are accounted for solely by metabotropic GABA_B_ at the postsynaptic sites. These percentages remain robust when removing weak synaptic connections.

We compared the GABA communication network to the synaptic connectivity between all cells as predicted by the recently updated *C. elegans* wiring diagram (**Figure 4B and 4G**)(Cook et al., 2019). We found that every postsynaptic partner of every GABA-releasing neuron expresses one of the two GABA_B_ receptors. In contrast, only 60% of postsynaptic partners of GABAergic neurons express any of the seven GABA_A_ receptors (Gendrel et al., 2016). Metabotropic communication similarly extends broadly over the cholinergic and glutamatergic signaling networks: 76% of the postsynaptic partners of cholinergic neurons express GAR receptors; 66% of the postsynaptic partners of glutamatergic neurons express MGL receptors (**Figure 4D**).

Surprisingly, we found that a considerable portion of the neurons expressing GABA receptors do not receive any connections from a presynaptic GABAergic neuron (**Figure 4E**); 31% of GABA_A_ and 37% of GABA_B_ receptor-expressing neurons have no presynaptic GABAergic partner. These results suggest widespread extrasynaptic GABA communication. Such extrasynaptic communication may also be present for the other metabotropic neurotransmitter receptors: 3% of neurons that express GAR receptors and 10% of neurons that express MGL receptors do not receive synaptic inputs from any glutamatergic and cholinergic neurons, respectively (**Figure 4E**). Notably, sensory neurons appear to be the most prominent recipients of extrasynaptic communication (**Figure 4F**).

### Uncovering determinants of cell fate

The 41 reporters used to assemble NeuroPAL each serve as indicators of neuronal differentiation and identity. Thus, mutations that affect cell fate can cause informative changes in the NeuroPAL color map as different neuron types acquire different genetic identities. Therefore, NeuroPAL provides a fast and unbiased method to screen for cell-fate alterations. As an example, the conserved TF PAG-3/Gfi has been shown to orchestrate the fates of the VA and VB ventral motor neuron classes (Cameron et al., 2002). We crossed NeuroPAL (*otls669*) with strains carrying null alleles of *pag-3* (*n3098* and *ok488*). As predicted, the color codes that identify the VA and VB cell types in NeuroPAL are absent in *pag-3* mutants (**Figure 5A-5B; Table S5**). Unexpectedly, we also found color changes in the AVE and PVR interneurons caused by *pag-3* mutation. By checking a fosmid-based reporter, we discovered *pag-3* expression in neurons that had gone previously unidentified, including AVE and PVR (**Figure 5C; Table S5**).

**Figure 5.**
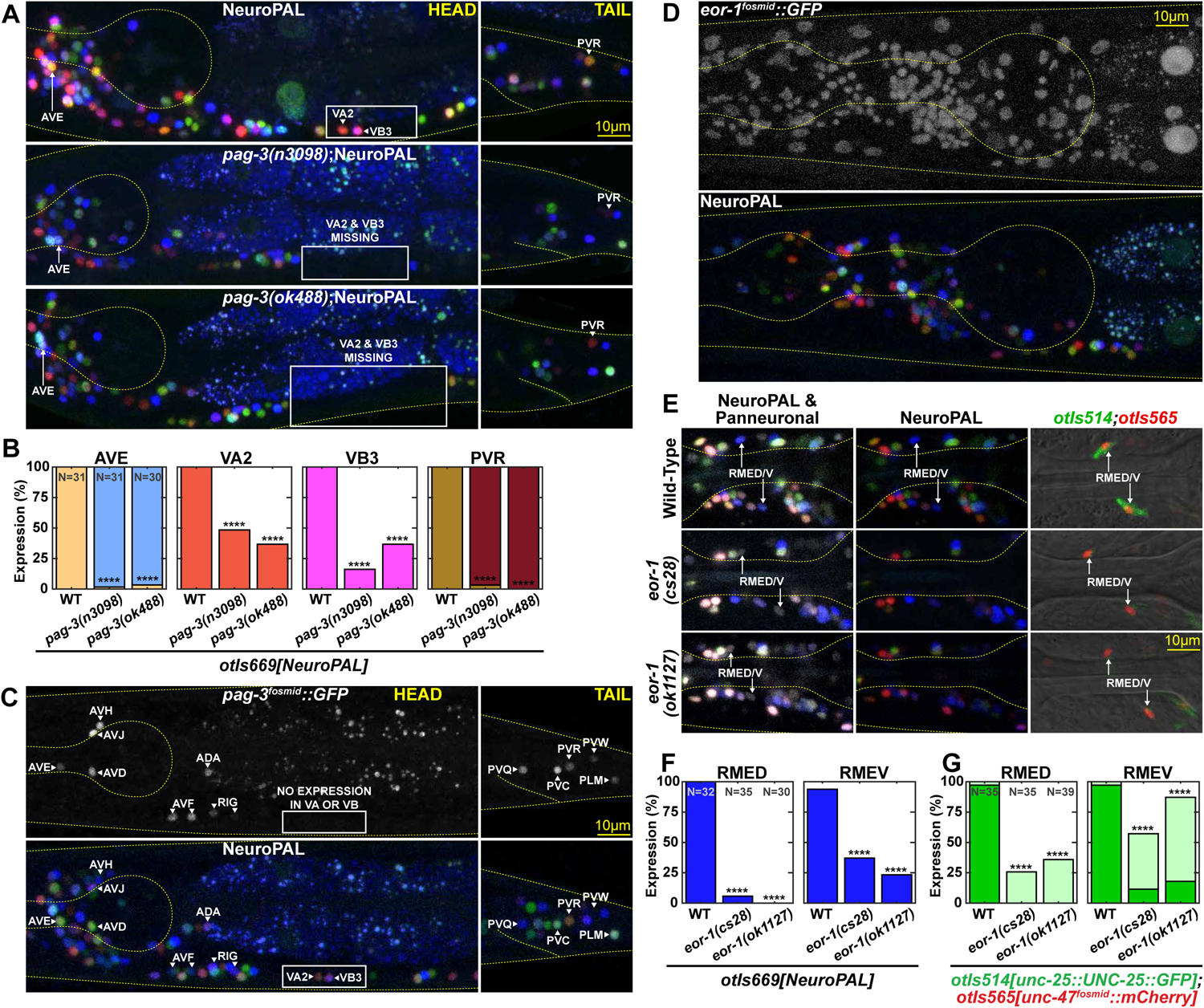
Mutant analysis of the conserved transcription factors PAG-3/Gli and EOR-1/PLZF See Table S5 for data. (A-B) Alterations in NeuroPAL coloring reveal neurons with altered fate in *pag-3(-)* mutant backgrounds: the AVE pre-motor interneuron gains blue-color expression, the PVR interneuron loses all green-color expression, and the VA and VB motor neurons are missing entirely. *\* Statistical bar graph colors in panel B are sampled from the images in panel A*. (C) PAG-3/Gli expression as assessed with a fosmid reporter (*wgIs154)*: **ADA**, ALM, **AQR, AVD, AVE**, AVF, **AVH, AVJ**, AVM, BDU, **DVC, I1, I2, I6**, PLM, **PQR, PVC**, PVM, **PVQ, PVR, PVW, RID**, RIG, **RMG**, VA11-12, and **URY** (previously-unpublished expression in bold). (D) EOR-1/PLZF has broad, likely ubiquitous, expression, as assessed with a fosmid reporter (*wgIs81*). (E-G) NeuroPAL’s blue reporters in RMED/V are *ggr-3, pdfr-1*, and *unc-25*. (E-F) NeuroPAL exhibits total loss of its RMED/V blue-color expression in *eor-1(-)* mutant backgrounds but retains panneuronal marker expression (shown in white), indicating the preservation of RMED/V neuronal fate. (G) RMED/V fate analysis in *eor-1(-)* mutants with *unc-25 (otIs514)* and *unc-47 (otIs565)* reporter transgenes.

NeuroPAL is particularly useful when the expression pattern of presumptive cell fate regulators are either unknown or too broad to easily formulate hypotheses about their effects on cell identity. As an example, we examined EOR-1/PLZF, a ubiquitously-expressed and highly-conserved TF (Howard and Sundaram, 2002). Given its ubiquitous expression (**Figure 5D**), EOR-1 may be involved in controlling the differentiation program of anywhere between none to all of the worm’s neurons. By crossing *eor-1* null mutants (*cs28* and *ok1127*) to NeuroPAL, we discovered neuron-subtype-specific differentiation defects. The dorsal and ventral RME neurons (RMED/V) lost all their blue coloring but retained their panneuronal label (**Figure 5E-5F; Table S5**). In contrast, the left and right RME neurons (RMEL/R) exhibited no changes in their color codes. Expression of the blue landmark fluorophore in the RME neurons is driven by three reporters: *ggr-3, pdfr-1*, and *unc-25/GAD* (**Table S1**). We validated the NeuroPAL color alterations with an endogenously tagged allele of UNC-25/GAD (**Figure 5G; Table S5**), but at the same time found no defects in UNC-47/VGAT expression. In conclusion we have identified a selective function of the ubiquitously expressed EOR-1 protein in the RME neuron class, a function that would have been impossible to predict based on the ubiquitous expression of EOR-1.

### Whole-brain activity imaging of gustatory and olfactory responses

A major challenge in analyzing panneuronal calcium imaging data in *C. elegans* has been determining neuronal identities (Kato et al., 2015; Nguyen et al., 2016; Venkatachalam et al., 2016). To solve this problem, we combined NeuroPAL with the panneuronally-expressed calcium reporter GCaMP6s (strain OH16230). We then used multicolor imaging to comprehensively identify all neurons. We recorded 21 worm heads (189 neurons) and, separately, 21 worm tails (42 neurons), with a median representation of 18 animals per neuron. We studied brainwide responses, in young-adult hermaphrodites, to a repulsive taste (160 mM NaCl) and two attractive odors (10^−4^ 2-butanone and 10^−4^ 2,3-pentanedione), each previously described to activate different sensory neurons. These stimuli were delivered in chemotaxis buffer to the nose of the animal using a multichannel microfluidic device (**Figure 6A-6B; Video S3-S4; Table S6; Methods)**(Si et al., 2019). We subjected each animal to the three chemical stimuli delivered in a randomized order (10 s pulses spaced by 50 s intervals). Using activity traces from all identified neurons, we assembled the mean brainwide response to each stimulus. Brainwide imaging revealed both known and novel neuronal responses to each stimulus, encompassing multiple sensory and interneurons, many of which were not previously implicated in behavioral responses (**Figure 6C**).

**Figure 6.**
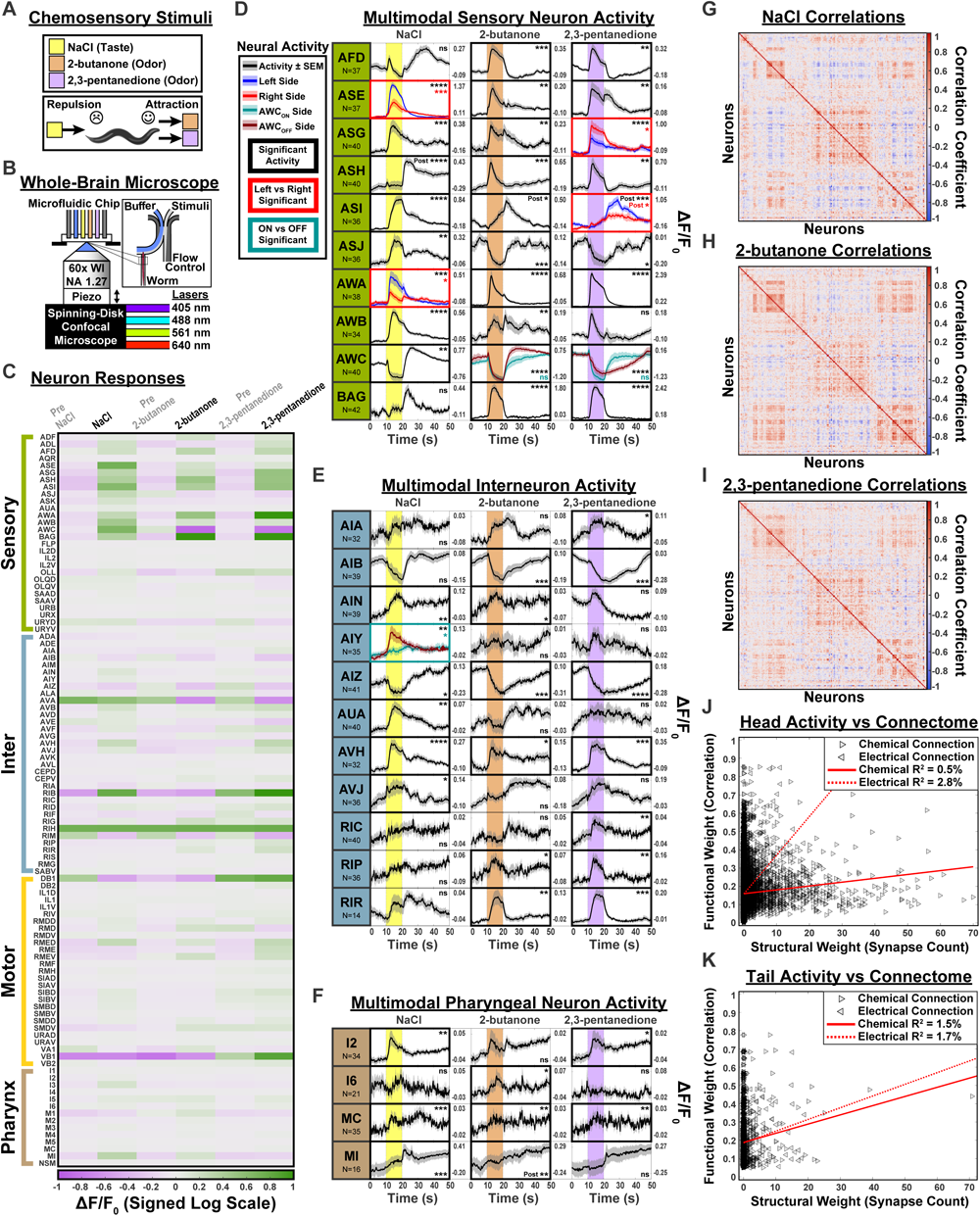
Whole-brain neuronal activity imaging of taste and odor responses See Table S6 for primary data. (A) *C. elegans* were subjected to three chemosensory stimuli: a repulsive taste (160 mM NaCl) and two attractive odors (10^−4^ 2-butanone and 10^−4^ 2,3-pentanedione). (B) Animals were immobilized inside a microfluidic chip. Stimuli were delivered in chemotaxis buffer. Each animal was imaged using a spinning disk confocal microscope with four excitation lasers. The NeuroPAL color map was imaged to identify all neurons. Thereafter, brainwide activity was recorded via the panneuronal calcium sensor GCaMP6s (**Video S3-S4**). (C) Peak neuronal activity, before and during stimulus presentation, for 109 head neuron classes. (D-F) Neuronal activity traces for selected (D) sensory neurons, (E) interneurons, and (F) pharyngeal neurons that responded to stimuli. The 10-second stimulus delivery period is indicated by the vertical colored bar. Black activity traces represent all neurons combined into one representative group. Colored activity traces divide neurons into groups exhibiting asymmetric responses. Two types of asymmetric groups are shown: stereotyped asymmetries between the left (L) and right (R) neurons and stochastic asymmetries between neurons corresponding to the L and R sides of the AWC^ON^ and AWC^OFF^ neuron pair. Significant responses (p or q ≤ 0.05) are highlighted by bold borders. Asymmetric L/R and AWC^ON/OFF^-sided responses are indicated by colored borders. “Post” = significant post-stimulus response. “ns” = no significant response. Note that the AWC neurons are stochastically asymmetric and, of the L/R pair, one neuron will differentiate into AWC^ON^ and the other into AWC^OFF^ (Alqadah et al., 2016). These AWC^ON/OFF^ identities correspond to the odor-specific preferences for our chosen stimuli (2-butanone and 2,3-pentanedione, respectively). NeuroPAL distinguishes these stochastic identities by using *srsx-3* to distinguishably color AWC^OFF^ (F-I) Average pairwise correlations between 189 neurons in the 30 seconds following onset of (G) NaCl, (H) 2-butanone, and (I) 2,3-pentanedione. All three correlation maps are presented on the same axes, determined by clustering the full-time-course correlations. The set of correlated and anti-correlated neurons differs for each stimulus presentation. (J-K) Comparison of functional activity to the connectome. We observe minimal correspondence between synapse counts and pairwise-functional-activity correlations for the (J) head and (K) tail.

As expected from previous work (Ortiz et al., 2009; Suzuki et al., 2008), NaCl evoked stereotyped and left/right (L/R) asymmetric responses in the two ASE neurons. However, ASER exhibited an increase in [Ca++] upon NaCl presentation, a result that likely reflects our choice of a much higher repulsive NaCl concentration, compared to previously-published experiments (Ortiz et al., 2009; Suzuki et al., 2008). We observed many novel responses, in a surprisingly large number of sensory and interneurons, corresponding to the NaCl stimulus pulse (**Figure 6C-6F; Table S6**). In particular, we detected significant NaCl responses in six interneurons with no previously known function: AIN, AVF, AVH, AVJ, I2, and MI. Strikingly, we also observed novel asymmetries in several neuron responses to NaCl. Among these, AWA exhibited stereotyped L/R activity: the AWAL response was significantly larger than AWAR (**Figure 6D; Table S6**). AIY interneurons also displayed left/right asymmetric responses to NaCl; this asymmetry was stochastic: either the left or the right AIY neuron displayed a stronger response to NaCl (**Figure 6E; Table S6**).

We found that odors also evoked responses in a surprisingly large set of both sensory and interneurons, throughout the brain, but exhibited more inhibitory activity (decreasing [Ca++]) than seen for NaCl (**Figure 6C**). As expected from previous work (Wes and Bargmann, 2001), 2-butanone and 2,3-pentanedione evoked stochastically asymmetric responses in the AWC neuron pair (**Figure 6D**). With 2,3-pentanedione, we also observed stereotyped L/R asymmetries in the ASG and ASI sensory neurons. The ASGL response was significantly smaller than ASGR. Upon stimulus removal, the ASIL response was significantly larger than ASIR. The set of neurons activated by all three stimuli was partly overlapping but distinct for each stimulus (**Figure 6C-6F**). For example, the ASJ sensory neuron was excited by NaCl but inhibited by both odors, whereas the RIC interneuron was excited solely by 2,3-pentanedione. Downstream of sensory neurons, we observed significant responses in many interneurons, many with no previously known function (e.g. AIN, AVH, RIF, RIG, RIR) (**Figure 6E-F; Table S6**). In the tail, the PVQ interneurons were significantly excited upon presentation of 2,3-pentanedione, while a mix of sensory and interneurons exhibited significant post-stimulus responses (upon removal of the stimulus from the animal’s nose) to all three stimuli (**Table S6**). Many of these post-stimulus responses were observed in interneurons with no previously known behavioral function (LUA, PVN, PVQ, PVR, and PVW).

Surprisingly, both salt and odors elicited responses across the pharyngeal nervous system (**Figure 6F; Table S6**), a heavily interconnected network of 20 neurons that synapses with the main nervous system through a single interneuron, RIP (Cook et al., 2019). Recent anatomical re-analysis of the pharyngeal connectome revealed that most neurons in the pharynx have potential sensory endings (Cook et al., 2020). Thus our activity recordings suggest that, despite its small size, the worm’s pharyngeal network may encode its own representation of behavioral responses to chemosensory cues.

We noted that in many animals, the pair of AVF interneurons displayed robust cyclical activity for the entire 4-minute recording, irrespective of delivery of gustatory or olfactory stimuli. The frequency of this activity was approximately 0.3 Hz, similar to the crawling frequency of freely-moving worms of this strain (**Figure S3; Table S2**). Previous experiments indicated that neurons in the region of the retrovesicular ganglion (the location of the AVF neurons) likely contribute to the central pattern generator for forward locomotion (Fouad et al., 2018). AVF ablation has also been shown to abolish forward locomotion (Hardaker et al., 2001). These results suggest that AVF may be part of the central pattern generator for forward locomotion in *C. elegans*. Further experiments will be required to confirm this hypothesis.

We compared the robustness of our imaging results to those collected from another strain (*otIs669;otIs672*, self-crossed 23x to drive isogenicity, OH15500). For OH15500 animals, we used a higher concentration and duration of NaCl (200mM for 20s) and delivered stimuli in water instead of buffer. We found similar results to those in OH16230 (detailed above) albeit with lower statistical significance due to lower sampling, N=7 heads (**Table S6**).

### Whole-brain neuronal dynamics and connectivity

Exploring network-level dynamics, we computed pairwise correlations between the activity of all identified neurons, in each animal, from our whole-brain activity recordings. We found similar stimulus-specific neuronal correlations across individuals, but each stimulus generated its own brainwide correlation pattern (**Figure 6G-6I**). Even the two attractive odors produced distinct sets of neuronal correlations among and between sensory and interneurons. These results can be seen in the brainwide neuronal trajectories through low-dimensional PCA-space (**Figure S4**). For instance, the AWB sensory neuron and its synaptically-connected interneuron partners AUA and AVH exhibit distinct pairwise correlations depending on the stimulus (**Figure 6D-6E**). Thus we find that brain dynamics are stimulus-specific.

We asked whether a simple relationship exists between these functional correlations and the synaptic counts previously measured from the anatomical connectome (Cook et al., 2019; White et al., 1986). To do so, we compared our correlation matrices of pairwise functional activity to the connectome matrix of pairwise synaptic connectivity. These matrices represent functional and structural measures of neuronal communication, respectively. We used the absolute value of the correlation matrices since the connectome lacks excitatory/inhibitory synaptic information. We found low Pearson correlation between these functional and structural matrices. For electrical connectivity, R^2^=2.8% in the head and R^2^=1.7% in the tail, and for chemical connectivity, R^2^=0.5% in the head and R^2^=1.5% in the tail (**Figure 6J-K**). We tried multiple variations in our calculations (e.g., using ranked correlation metrics, log-scaling synaptic counts, and limiting functional activity to only stimulus or non-stimulus delivery periods) but these did not noticeably improve correlation between functional activity and the structural connectome (**Table S6**). The low correlation values we measured suggest considerable contributions from non-synaptic signaling networks (which are not reflected in the anatomical connectome). These non-synaptic signaling networks likely include: a) pervasive neuropeptidergic signaling (Bargmann and Marder, 2013); b) extensive aminergic signaling (Bentley et al., 2016); and c) potential extrasynaptic signaling as hinted by our expression maps of the metabotropic neurotransmitter receptors (see **Expression maps for all metabotropic neurotransmitter receptors**). However, we note that spatiotemporal limitations of our calcium imaging and synaptic variability in the anatomical connectome may have further contributed to the low correlation values we measured between functional activity and synaptic connectivity.

### Semi-automated neural identification

We developed an instruction guide to help researchers use NeuroPAL (**NeuroPAL Manuals: https://www.hobertlab.org/neuropal/**). This guide covers a variety of NeuroPAL-compatible microscope configurations and provides instructions on how to identify all neurons using the NeuroPAL color map. However, manual annotation of neurons is laborious and time-consuming. To speed annotation, we developed a software pipeline that partially automates this task (**Text S1; NeuroPAL ID Software: https://www.hobertlab.org/neuropal/**). This software pipeline uses three unsupervised algorithmic steps to automatically annotate neuronal identities in NeuroPAL images (**Figure 7A**). First, we filter out non-neuronal fluorescence. Second, we detect the color and position of each neuron. Third, we compute a probabilistic estimate of each neuron’s identity using a statistical atlas of NeuroPAL colors and positions (see **Variability in neuronal cell body positions**)(Varol et al., 2020). Lastly, a graphical user interface (GUI) permits manual review and error correction of all steps in our unsupervised pipeline.

**Figure 7.**
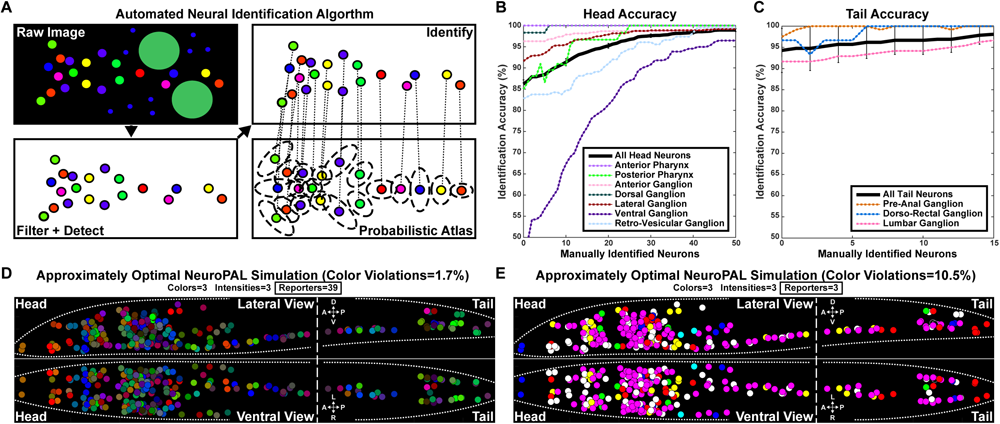
NeuroPAL software: an algorithm for semi-automated neuronal identification and an algorithm to generate optimal-coloring solutions for cell identification See Text S1-S2 for algorithmic details and validation. (A-C) The algorithm used for semi-automated neuronal identification. (A) Raw images are filtered to remove non-neuronal fluorescence and neurons are detected in the filtered image. Detected neurons are identified by matching them to a statistical atlas of neuronal colors and positions (**Table S3**). (B,C) Semi-automated neuronal identification accuracy begins at 86% for the head and 94% for the tail. Manually identifying eight neurons raises the head accuracy above 90%. Overall accuracy is displayed as a black line. Accuracy for each ganglion is displayed as a dotted, colored line (see legend). Many of the neurons and ganglia have high identification accuracy and confidence. The ventral ganglion is a problematic area, likely due to the density and high positional variance therein. (D-E) The algorithm used to generate optimal-coloring solutions for cell identification (for any collection of cells in any organism). We show simulations of two approximately optimal alternatives to NeuroPAL (**Table S7**), one that permits as many reporters as NeuroPAL (D) and one that restricts the transgene to only 3 reporters (E). With the exception of the number of reporters, both alternatives were generated using parameters similar to NeuroPAL: three landmark fluorophores, where each fluorophore is distinguishable at three intensities (high, medium, and low). Reporters were chosen by the algorithm from those available in WormBase, a community-curated database of cell-specific reporter expression. Similar databases are available for other model organisms (e.g., fly, fish, and mouse). We evaluated the two NeuroPAL alternatives by computing the percentage of their color violations, defined as neighboring neuron pairs with indistinguishable colors.

We evaluated the semi-automated neuronal identification performance of our pipeline. To do so, we cross-validated its performance and found our accuracy varied across ganglia (**Figure 7B-7C**). Accuracy was 86% for the head and 94% for the tail. Accuracy largely depended on cell density; for example, our pipeline achieved high accuracy for all tail ganglia but lower accuracy for the much denser ventral ganglion in the head. Our software incorporates supervised annotation of low-probability neuronal identities to improve the estimated identities of the remaining unlabeled neurons. Adding eight manual annotations, on average, brings the head accuracy above 90% (**Figure 7B**).

Our algorithm also provided a means of assessing the importance of color information in assigning cell identities. When we restricted the model to assign identities only on the basis of location, automated accuracy dropped to 50% for the head and 68% for the tail (**Text S1**). These results confirm a substantial improvement in accuracy with the color information provided by NeuroPAL.

### An optimal-coloring algorithm for other tissues and model organisms

We built NeuroPAL empirically, laboriously testing a large variety of reporter-fluorophore combinations. Our method can benefit research into other tissues and model organisms that require cell-specific identification. For example, in **Figure S5**, we suggest a design for a general-purpose “FlyPAL” that, among other applications, might be well suited for studying the neuronal circuit of ecdysis (*Drosophila* molting behaviors). To facilitate the generation of further analogous multicolor landmarking solutions, for any collection of cells in any organism, we developed an algorithm that computes approximately optimal reporter-fluorophore combinations to test *in vivo* (**Text S2; Optimal-Coloring Software: https://www.hobertlab.org/neuropal/**). With our optimal-coloring software, the amount of empirical testing required to construct future multicolor landmarking reagents is significantly reduced.

As an overview, in order to be individually identifiable, neighboring cells of different types must be distinguishable from each other. Cells that cannot be distinguished by morphology must be distinguished by color and/or intensity differences larger than a discrimination margin. In our software, the user chooses this margin, decides which cells must be distinguishable from each other, specifies the number of landmark fluorophores to use in combination with a list of available reporters that have known expression, and restricts the total number of reporter-fluorophore combinations permitted by their transgenesis techniques. Given these inputs, the algorithm generates multiple approximately optimal solutions (reporter-fluorophore combinations) that minimize the percentage of color violations, defined as neighboring neuron pairs which fall below the discrimination margin.

We ran our optimal-coloring algorithm using Wormbase (a database of worm reporter expression) to generate several NeuroPAL alternatives **(Text S2**). In our simulations, we achieve color violations of 1.7% when using 39 reporters and 10.5% when restricting the solution to only 3 reporters (**Figures 7D-7E**). This low percentage of indistinguishable neurons, in our two simulated NeuroPAL alternatives, indicates that they could prove beneficial for neuronal identification.

Our algorithmic and FlyPAL examples illustrate how the NeuroPAL-technique can be extended to offer multicolor landmarking solutions for any collection of cells, in any model organism, using similar databases of reporter expression (e.g., using Flybase for fly, ZFIN for zebrafish, and MGI for mouse). While transgenesis techniques in other model organisms may restrict the total number of reporters that can be used, we show that despite this limitation our algorithm can still offer beneficial solutions to devise stereotypic color maps for cellular identification.

## DISCUSSION

Understanding the nervous system requires an integrated view of its constituent molecular, cellular, and functional signaling networks. A primary bottleneck to mapping these networks across an entire brain has been the difficulty of reliably identifying neuronal cell types. Here, we introduced NeuroPAL, a tool that allows researchers to use a single multicolor landmark strain to determine all neuron identities in the *C. elegans* nervous system. The fluorophores used to build NeuroPAL were chosen to have negligible emission overlap with CFP, YFP, and GFP/GCaMP, while maintaining compatibility with common illumination sources and filter sets. This allows NeuroPAL to be used in conjunction with many different types of fluorescent reporters for diverse studies of gene expression, nervous-system development, and brain activity, as we have demonstrated here. NeuroPAL offers comprehensive neuronal identification, and thus substantially improves on an elegant, recently-published system that identifies, with some uncertainty, less than 60% of all neurons in the *C. elegans* nervous system (Toyoshima et al., 2020).

One major challenge in understanding the molecular networks that encode the development and function of the nervous system lies in mapping gene-expression patterns that determine cell fates and characterize the effect of genetic perturbations on neuronal fate. We have shown here that NeuroPAL can be used to identify sites of gene expression and, independently, can also be used as a cell-fate marker. By detecting mutation-induced perturbations in the NeuroPAL color map of gene expression, we uncover a neuron-fate-specific role for EOR-1/PLZF, an ubiquitously expressed and highly-conserved TF. We also completed the expression pattern of the well-studied TF PAG-3/Gli, and discovered new neuronal fate alterations caused by mutations therein. These findings demonstrate the effectiveness of NeuroPAL in screening for fate-specific roles of TFs. As a next step, we envision NeuroPAL being used to screen larger collections of TF-specific mutations or, in a totally unbiased approach, being used in conjunction with the comprehensive Million Mutant project (Thompson et al., 2013) to identify genes involved in neuronal differentiation across the nervous system.

The synaptic connectivity of the *C. elegans* nervous system has been mapped by electron microscopy. However, the pathways of neuronal communication therein cannot be fully understood without corresponding maps of neurotransmitter and receptor expression. Recent work has mapped the identities of all neurons that release each neurotransmitter type (Gendrel et al., 2016; Pereira et al., 2015; Serrano-Saiz et al., 2013). But mapping the neurons that “listen” to each neurotransmitter has been difficult due to the large diversity of receptor types and their often broad expression patterns. We used NeuroPAL to map all neurons that express every type of metabotropic neurotransmitter receptor encoded in the worm genome. We find that the breadth of the metabotropic communication network is far more extensive than previously thought. Metabotropic neurotransmitter receptors cover 97% of all neurons and are able to participate in 75% of all synaptic connections from neurotransmitter-releasing neurons.

Our contribution of the map of metabotropic GABA receptor (GABA_B_) expression, combined with previous work mapping the ionotropic GABA receptors (GABA_A_) (Bamber et al., 1999; Beg and Jorgensen, 2003; Gendrel et al., 2016; Jobson et al., 2015), completes the GABA communication network with startling results. First, GABA_B_ receptors are expressed in every postsynaptic partner of all GABA-releasing neurons, whereas GABA_A_ receptors are expressed in only 60% of these “listeners”. Metabotropic reception may be the most common means of GABA signaling in the worm nervous system. Second, many neurons that express GABA receptors are not postsynaptic partners of any GABAergic neuron. Previous work has shown that extrasynaptic GABA signaling can occur between specific cell types in *C. elegans* (Jobson et al., 2015). Our results suggest that extrasynaptic GABA signaling may be far more prevalent than previously thought. Indeed, GABA signaling may be an important means of integrating circuit activity throughout the nervous system. Only 10% of all worm neurons release GABA but 92% of all neurons express GABA receptors and yet, just 66% of this GABA-signaling network is synaptically connected. These results underscore the need for unbiased and brainwide analysis of how functional activity is shaped by neurotransmitters and receptors. However, we caution that our extrasynaptic communication findings may be the result of incomplete anatomical synaptic annotations and/or that metabotropic receptors may potentially serve as promiscuous postsynaptic partners for other signaling ligands. We envision future work, using NeuroPAL, to map expression of the remaining aminergic and ionotropic neurotransmitter receptors (Fernandez et al., 2020).

To date, functional networks have been investigated by recording the activity of small subsets of labeled neurons. More recent work has inaugurated whole-brain activity imaging with cellular resolution (Ahrens et al., 2013; Lemon et al., 2015; Mann et al., 2017; Schrodel et al., 2013). However, the inability to reliably identify all neurons within whole-brain recordings has precluded a full picture with circuit-level details. Thus, principal component analysis (PCA) has been commonly employed to construct low-dimensional representations of brain dynamics in individual animals, but the lack of a common basis has hampered animal-to-animal comparisons (Linderman et al., 2019). Coupling NeuroPAL with whole-brain activity imaging methods permits a unified view of network dynamics, across animals, without sacrificing circuit-level details.

Here, we used NeuroPAL for whole-brain imaging of panneuronal calcium dynamics in response to chemical stimuli. The three well-studied chemical stimuli that we used have been known to evoke activity in a small number of sensory neurons and interneurons. Our results show that the set of stimulus-evoked responses engage the nervous system far more broadly than previously realized, extending across many sensory neurons and interneurons, even those in the synaptically-isolated pharyngeal nervous system. In our studies, we discovered interneurons with no previously-known functions exhibited significant responses to anywhere from one to all three chemosensory stimuli.

Surprisingly, we found that many neurons respond with distinct dynamics to the two olfactory attractants that we used (2-butanone and 2,3-pentanedione) and the one gustatory repellent (a high concentration of NaCl). Our data revealed new neural asymmetries in the nervous system, both deterministic left/right asymmetries and stochastic ON/OFF asymmetries, that also escaped previous analyses. Thus asymmetric activity in the worm nervous system is more widespread than the well-studied examples of the stereotyped ASEL/ASER pair and the stochastic AWC^ON^/AWC^OFF^ pair. Comprehensive neuronal identification enabled us to examine the relationship between whole-brain activity and the connectome but we found no strong correlations between them. Unifying functional and anatomical views of the nervous system will require a deeper understanding of the properties of synaptic communication and the neuromodulation of activity patterns; we expect our work to aid in these endeavors (Bargmann and Marder, 2013; Brennan and Proekt, 2019; Kaplan et al., 2020; Kato et al., 2015). A richer set of network responses, even for simple chemosensory inputs, has broad relevance in understanding sensorimotor processing. Our work indicates that even simple behaviors employ large portions of the worm nervous system, engaging different brainwide neuronal correlations across behaviors.

We provide semi-automated software, with high initial neuronal-identification accuracy, wherein manual annotations progressively improve this accuracy. We also provide software which determines reporter and color assignments to distinguish any set of cells. Our ongoing efforts aim to further improve automated neuronal-identification accuracy, expand the software to cover all life stages of both sexes, and extend its pipeline to extract activity traces with neuronal identities.

Using a stereotyped multicolor landmark strain to easily identify neurons may be extended to other genetically-tractable animals including fruit fly and zebrafish. As their brains are considerably larger, generating multiple landmark strains that in aggregate cover their whole brain would require community-wide efforts. However, deterministic multicolor labeling of individual brain regions of interest should be tractable. Important and well-studied brain regions in these systems (e.g., the mushroom body or nerve cord of the fly and the olfactory bulb of the zebrafish) have substantial cellular diversity in a limited number of cells, making them good targets for a tool like NeuroPAL.

## STAR METHODS

### RESOURCE AVAILABILITY

#### Lead Contact

Further information and requests for resources and reagents should be directed to and will be fulfilled by the Lead Contact, Eviatar Yemini.

#### Materials Availability

All NeuroPAL integrant *C. elegans* strains are available at the *Caenorhabditis* Genetics Center (CGC).

#### Data and Code Availability

The imaging datasets generated in this study are freely available at:
https://zenodo.org/record/3906530

The code and software generated for this study are freely available at:

https://github.com/amin-nejat/CELL_ID

https://github.com/Eviatar/Optimal_Coloring

https://github.com/venkatachalamlab/NeuroPAL-traces/

### EXPERIMENTAL MODEL AND SUBJECT DETAILS

#### Worm Maintenance

All worms were raised at 20°C, on NGM plates, and fed OP50 *E. coli* as previously described (Brenner, 1974), unless otherwise noted.

#### Plasmids and injections

Fluorophores were ordered from IDT and/or cloned via standard techniques (Gibson, restriction-free, T4 ligase, or QuikChange™ mutagenesis) into the pPD95.62 Fire vector (a gift from Andrew Fire)d.

The 1.4 kb synthetic ultra-panneuronal driver (UPN) was generated by fusing *cis*-regulatory elements from four different panneuronally expressed genes: *unc-11*^*prom8*^, *rgef-1*^*prom2*^, *ehs-1*^*prom7*^, *ric-19*^*prom6*^ (Stefanakis et al., 2015). Fusion was done in a single quadruple PCR promoter fusion (Hobert, 2002). All cloned NeuroPAL reporters were made via PCR, gel purified, then inserted using standard techniques into the fluorophore vectors. To accommodate the large number of reporters and conserve space in extrachromosomal arrays, in place of plasmid backbones we used linear DNA amplified via PCR. Linear DNA has also been shown to improve expression levels (Etchberger and Hobert, 2008). Therefore, all injected NeuroPAL reporters were PCR amplified and gel purified to remove their vector backbone. Injection mixes consisted of complex arrays, with sheared bacterial DNA serving as spacers, to minimize potential crosstalk amongst reporters. Preliminary NeuroPAL strains were injected as complex arrays into *pha-1(e2123)* with *pBX[pha-1(*+)] to rescue at the selection temperature 25°C (Granato et al., 1994). Final NeuroPAL strains were injected into the wild-type, N2, without *pha-1(*+). We used the *rab-3* reporter to drive panneuronal GCaMP6s expression. We noted that sensory neurons exhibited weaker GCaMP6s expression and thus supplemented the *rab-3* reporter with the *arrd-4* pansensory reporter. The panneuronal GCaMP6s reporters were injected as complex arrays into N2. Integrations were performed using gamma irradiation. All integrant worms were outcrossed 8x. Non-integrant strains that were used to identify metabotropic and transcription factor expression were injected with *pAB1[inx-6*^*prom18*^::*TagRFP-T]* (an anterior pharyngeal marker) and *pBX[pha-1(*+)] into *pha-1(e2123)*, then raised at 25°C for selection. All reporters and their injected concentrations are in **Table S1**.

#### Transgenic and mutant strains

Transgenic and mutant strains used in this study are available in the supplement (**Table S1**).

### METHOD DETAILS

#### Worm phenotyping

Brood-size quantification, high-resolution behavioral phenotyping, dye-fill with DiO, chemotactic quadrant assays, and drop-test assays were performed using standard protocols (Bargmann et al., 1993; Chase and Koelle, 2004; Hedgecock et al., 1985; Hilliard et al., 2002; Yemini et al., 2013).

#### NeuroPAL imaging

NeuroPAL imaging can be performed using a wide variety of microscopes as further detailed in the manual “Configuring Your Microscope for NeuroPAL” (**NeuroPAL Manuals: https://www.hobertlab.org/neuropal/**). We imaged strains with a Zeiss LSM880, equipped with 7 laser lines: 405, 458, 488, 514, 561, 594, and 633 nm. Our standard configuration employed 405, 488, 561, and 633 nm to excite mTagBFP2, GFP/GCaMP + CyOFP1, TagRFP-T, and mNeptune2.5, respectively. The 8-color emission spectra (**Figure S1** was captured using strains that expressed each fluorophore individually. For these, we used the LSM880’s “lambda mode”, employing its 32-channel spectral detector to capture color spectra from 391-727nm, at ∼10 nm color resolution – several fluorophores were imaged by exciting them with wavelengths below peak excitation and significantly increasing both the laser power and gain. To that end, for the 8-color emission spectra, we used: 405 nm to excite mTagBFP, CFP, GFP, and CyOFP1; 488 nm to excite YFP and mNeonGreen; and, 561 nm to excite TagRFP-T and mNeptune2.5. All NeuroPAL reporter and mutant crosses were imaged with the same scope. When not performing a DIC overlay, gamma correction of ∼0.5 was applied to images so as to improve color visibility. Occasionally, histograms were adjusted to balance colors for visibility. These image adjustments are necessary and suggested for NeuroPAL identification in order to deal with a variable range of GFP/CFP/YFP reporters and color alterations in mutant backgrounds.

#### Whole-brain imaging setup with microfluidic stimulus delivery

To record whole-brain neuronal activity, while presenting an animal with chemosensory stimuli, we employed a modified version of a microfluidic system that delivers multiple odors to Drosophila larvae (Si et al., 2019). We adapted this system for use with *C. elegans* (Chronis et al., 2007). This microfluidic chip allowed us to record intact animals while presenting precisely timed stimuli. OH16230 animals were washed in M9 buffer containing 1 mM tetramisole hydrochloride to minimize motion, then placed in CTX buffer and loaded into the microfluidic chip (Larsch et al., 2013). For each animal, we first obtained a high-resolution 4-color landmark volume for neuronal identification. We then observed a 2-minute unilluminated waiting period to allow the animal to recover from its exposure to laser light. Lastly, we performed a 4-minute, single-channel recording to capture nuclear GCaMP6s activity, at a frequency of ∼4 Hz. During this recording, at the end of every minute, we presented 10 s pulses of 160mM NaCl 10^−4^ 2-butanone, and 10^−4^ 2,3-pentanedione. Stimuli were presented in CTX buffer maintaining a constant pH 6 and 350 mOsm/L across stimulus and non-stimulus presentation. The order of stimulus presentation was rotated periodically to avoid sampling order-dependent stimulus responses. After the experiments completed, we identified neurons within the 4-color identification volumes. We correlated these identified neurons in the landmark volume to their GCaMP6s recordings, and extracted their traces using deformable non-negative matrix factorization (dNMF)(Nejatbakhsh et al., 2020). To facilitate comparison, we downsampled recordings to the lowest frame rate observed within their group, 4 Hz for OH16230 heads and 3.87 Hz for OH16230 tails. Neurons within 5μm distance of each other, whose traces showed at least 95% correlation, were deemed too similar and removed. In the supplement we present similar experiments performed using OH15500 animals (**Table S6**). For OH15500 we used DI water instead of CTX buffer, the three stimuli remained the same but NaCl was presented for 20 s at 200 mM, neuronal traces were extracted using a previous method that predates dNMF (https://github.com/venkatachalamlab/NeuroPAL-traces/)(Venkatachalam et al., 2016), and visual inspection was used to identify and remove traces from neighboring neurons that exhibited mixed signals. OH15500 heads were downsampled to 4.1 Hz.

### QUANTIFICATION AND STATISTICAL ANALYSIS

#### Mutant analysis

Statistical analysis of NeuroPAL-color alterations, for four neuron classes, in two *pag-3(-)* mutant backgrounds was Bonferroni corrected for eight tests. We used a One-Sided Fischer’s Exact Test to analyze mutant-induced changes in PVR, VB2, and VB3. We used a One-Sided Rank Sum Test to analyze mutant-induced changes in AVE, scoring 0 if no changes were observed, 1 if either AVEL or AVER was altered, and 2 if both AVE neurons were altered. Statistical analysis of NeuroPAL color loss and UNC-25 reporter loss, for the RMED and RMEV neurons, in two *eor-1(-)* mutant backgrounds was Bonferroni corrected for eight tests. We used a One-Sided Fischer’s Exact Test to analyze mutant-induced loss of RMED/V blue coloring in NeuroPAL. We used a One-Sided Rank Sum Test to analyze mutant-induced changes in the RMED/V UNC-25 reporter expression, scoring 0 if expression was lost, 1 if expression was weak but visible, and 2 if no changes were observed.

#### Analysis of whole-brain imaging data

To analyze the whole-brain imaging stimulus responses, we reviewed ASE responses to salt and AWC responses to the odors – the primary sensory neurons for these stimuli. Worms were marked as stimulus responsive if either their left or right neuron showed the published response to their corresponding stimuli (ASE excitation for NaCl and AWC inhibition for odors) (Chalasani et al., 2007; Suzuki et al., 2008). 21 heads responded to all three stimuli, providing strong internal controls to compare their circuit activity across all stimuli. Additionally, 21 tails were included, without response verification as the heads of these animals were not simultaneously imaged. Premotor interneurons (AVA, AVB, AVD, AVE, PVC) and ventral-cord motor neurons (AS, DA, DB, DD, VA, VB, VC, VD) show spontaneous cyclical activity and thus were excluded from significance testing. We used t-tests (2-tailed, paired) to compare the mean signal during stimulus presentation with an identical period immediately prior, within the very same neuron (a strong internal control). These p-values were corrected for multiple testing using false discovery rate (FDR) adjusted q-values (Storey, 2002). Post-stimulus responses were identified by first ensuring the neuron’s mean stimulus response exceeded its pre-stimulus mean (to avoid mistaking repolarizing calcium activity for a post-stimulus response), then testing whether the post-stimulus mean exceeded the mean stimulus response (1-tailed, paired t-test). Post-stimulus p-values were corrected using FDR. We present blue light responses in the supplement (**Table S6**). Since GCaMP6s requires 488 nm excitation, we were unable to record a pre-stimulus period for blue-light responses. We reasoned that since worms habituate to light (as evidenced by our traces and previous reports (Liu et al., 2010), the blue-light response could instead be tested in reverse by comparing the 10s immediately after lights on, to the 10s period thereafter. Our protocol of two minutes in the dark, just after the identification volume, ensures that the 488 nm GCaMP6s excitation laser is a sudden, strong stimulus, evoking an immediate aversive response. Light comparisons similarly employed a 2-tailed, paired t-test and FDR corrections. To test asymmetric neuron responses we used 2-tailed, two-sample (unpaired) t-tests.

#### Statistical analysis software

All statistics and code were run in MATLAB, using standard toolboxes, with the exception of the OME Bio-Formats API (used to read in Zeiss CZI and Nikon ND2 file formats) (Linkert et al., 2010), dNMF (used to extract whole-brain calcium activity)(Nejatbakhsh et al., 2020), Matlab geometry toolbox for 2D/3D geometric computing (https://github.com/mattools/matGeom), and MathWorks FileExchange functions (**Text S1**).

## ADDITIONAL RESOURCES

NeuroPAL imaging can be performed using a wide variety of microscopes, as further detailed in the manual “Configuring Your Microscope for NeuroPAL.” Manuals for setup and cell identification are available online at: https://www.hobertlab.org/neuropal/.

## AUTHOR CONTRIBUTIONS

All authors contributed in writing this manuscript. OH initiated the project, EY designed and built the NeuroPAL and conducted all metabotropic receptor, and mutant experiments, generated the panneuronal GCaMP6s strain, and performed the non-automated identification of activity traces. EY designed and performed all non-stimulus behavioral phenotyping. AL designed and performed all chemotactic assays. AL, AS, and VV designed and built the whole-brain imaging scope and microfluidic device then, together with EY and OH, designed the whole-brain imaging experiments which were all conducted in the lab of AS. AL performed all whole-brain imaging experiments. AN, EV, LP, and VV designed and built the software for whole-brain activity imaging that connects neurons with their identities, then extracts, de-mixes, and normalizes neuron activity traces. AL, EY, and VV designed and built the software to analyze the whole-brain imaging data. AN, EV, GEM, LP, and RS computed the neuronal positional variability atlas, designed the semi-automated identification algorithms and software with accuracy validations, and together with EY built the GUI. EV designed and built software to compute coloring for NeuroPAL applications and EY generated the accompanying analysis. EY designed the FlyPAL concept. Correspondence about calcium imaging and physiology is to be addressed to AL, VV, and EY. Correspondence about algorithms for semi-automated cell identification is to be addressed to AN, EV, and EY.

## ACKNOWLEDGEMENTS

We thank Qi Chen for generating transgenic lines. We thank Molly Booth Reilly, Ibnul Rafi, and Emily Berghoff for alpha testing the NeuroPAL software. We thank Eduardo Leyva-Díaz for *ehs-1*^*prom7*^. We thank David Hall, Zeynep Altun, and Chris Crocker for use of the WormAtlas N2S/T/U EM reconstructions. We thank Michael Z. Lin, Vladislav Verkhusha, and Robert E. Campbell for their considerable help in choosing fluorescent proteins. We thank Benjamin White and Wesley Grueber for their considerable help conceptualizing NeuroPAL applications in Drosophila. We thank Michael Koelle for his generous gift of the gbb-2 reporter fosmid. We thank Scott Linderman and Andrew Leifer for many helpful discussions regarding automated neural identification. We thank members of the Hobert, Chalfie, Paninski, and Samuel labs for comments on the manuscript. Some strains were provided by the CGC, which is funded by NIH Office of Research Infrastructure Programs (P40 OD010440).

## SUPPLEMENTAL FIGURES

**Figure S1. Emission for NeuroPAL fluorophores and compatible signal fluorophores**. Fluorophore emission spectra were collected, at ∼10nm color resolution, using strains that expressed each fluorophore individually.

**Figure S2. NeuroPAL phenotypes**. (A) Brood size. (B) Dead eggs. (C) Crawling speed. (D-F) Chemotactic preferences for (D) 2-butanone, (E) 2,3-pentanedione, and (F) NaCl.

**Figure S3. Continuous cyclical activity in AVF neurons approximately matches the worm’s freely-moving crawling frequency**. (A) AVF left and right (blue and red) neuronal traces (median filtered at 0.5 s), from five OH16230 worms, display cyclical activity of ∼0.3 Hz (counting the number of peaks per 4-minute interval). (B) The freely-moving crawling frequency measured in NeuroPAL strains. OH16230 (enclosed within a black square) exhibits a crawling frequency of ∼0.3 Hz.

**Figure S4. Whole-brain neuronal dynamics in response to chemosensory stimuli**. (A) Average phase trajectories (N=21 animal heads) of whole-brain neuronal activity in response to stimuli (10 s of stimulus application and 20s thereafter). Trajectories are presented for 160 mM NaCl (yellow), 10^−4^ 2-butanone (orange), and 10^−4^ 2-3-pentanedione (purple), with the standard error of the mean shown as a mesh. The axes shown are the first three principal components of whole-brain activity. The average stimulus trajectories exhibit similarities but remain distinguishable. (B) The Pareto chart of variance explained per principal component.

**Figure S5. A *Drosophila* FlyPAL design concept**. (A) A “Universal-Reporter Line” for flies employing three orthogonal binary-expression systems to independently drive each of the three NeuroPAL landmark fluorophores (Brand and Perrimon, 1993; Kakidani and Ptashne, 1988; Lai and Lee, 2006; Potter et al., 2010). Optional T2A sequences enable co-expression of GCaMP6s and TagRFP-T to facilitate ratiometric calcium imaging. Genetic insulators are used to insulate the expression of each landmark fluorophore. ΦC31 integration is used to integrate this transgene into the fly genome. (B) An “Ecdysis-Specific Driver Line” employs five short enhancer fragments to drive expression of the NeuroPAL landmark fluorophores using their corresponding binary systems (Diao et al., 2017; Kim et al., 2006). (C) Crossing the Universal Reporter and Ecdysis-Specific Driver lines generates an “Ecdysis FlyPAL” wherein ecdysis-specific neurons can be identified by their color and position. Ecdysis-specific neurons can then be individually identified in calcium activity recordings.

## SUPPLEMENTAL VIDEOS

**Video S1. NeuroPAL head**. 3D image volume of a NeuroPAL worm head.

**Video S2. NeuroPAL tail**. 3D image volume of a NeuroPAL worm tail.

**Video S3. NeuroPAL head neuronal activity (panneuronal GCaMP6s)**. Maximum intensity projection of a NeuroPAL head, overlaid with the panneuronal activity-sensor GCaMP6s (strain OH16230). Neuronal activity and stimulus responses (white signal) are superimposed onto the multicolor neuronal-identification image. The video is sped up 5x. Stimuli were delivered in the following order: 10^−4^ 2,3-pentanedione, 10^−4^ 2-butanone, then 160mM NaCl. A selection of neurons, covering most head classes, are identified via labels above them. Premotor interneurons (AVA, AVB, AVD, and AVE) are labeled in red. Ventral-cord motoneurons (DB and VB) are labeled in pink. Neurons identified as exhibiting global motor dynamics (AIB, RIB, RIM, RIV, RME, and SMD)(Kato et al., 2015) are labeled in brown. The AVF neurons, labeled in turquoise, exhibit an oscillation period of 4.5±1.2 s and a phase differential between them of 1±0.7 s (mean ± standard deviation). Neurons exhibiting a strong response to specific stimuli remain labeled around the time the stimulus was applied. The exact stimulus application periods are shown via a label at the top right corner. AWC is inhibited and BAG is excited in response to both odors. ASEL, ASI, and AWB are excited in response to salt. AFD is excited upon salt removal.

**Video S4. NeuroPAL tail neuronal activity (panneuronal GCaMP6s)**. Maximum intensity projection of a NeuroPAL tail, overlaid with the panneuronal activity-sensor GCaMP6s (strain OH16230). Neuronal activity and stimulus responses (white signal) are superimposed onto the multicolor neuronal-identification image. The video is sped up 5x. Stimuli were delivered in the following order: 10^−4^ 2-butanone, 160mM NaCl, then 10^−4^ 2,3-pentanedione. A selection of neurons, covering most tail classes, are identified via labels above them. Premotor interneurons (PVC) are labeled in red. Ventral-cord motoneurons (AS, DA, DD, VA, and VD) are labeled in pink. The exact stimulus application periods are shown via a label at the top right corner.

## SUPPLEMENTAL TABLES

**Table S1**. NeuroPAL and Panneuronal GCaMP6s Transgene Data, Neuron-Specific Color Expression, Neuronal Identity Verification, and Strains, Related to Figures 1 and 6 and STAR Methods

**Table S2**. NeuroPAL Health, Phenotypes, and Chemotaxis, Related to Figures S2 and S4

**Table S3**. Neuron Locations, Positional Variability, and Color Variability, Related to Figures 2 and 7 and Text S1

**Table S4**. Metabotropic Neurotransmitter Receptor Expression, Statistics, and Strains, Related to Figures 3 and 4 and STAR Methods

**Table S5**. Transcription-Factor Mutant Expression, Cell-Fate Statistics, and Strains, Related to Figure 5 and STAR Methods

**Table S6**. Whole-Brain Imaging Statistics and Connectomic Correlations, Related to Figure 6

**Table S7**. Optimal-Coloring Software-Generated Reporter-Fluorophore Combinations, Related to Figure 7 and Text S2

## SUPPLEMENTAL TEXT

**Text S1**. Semi-Automated Neuronal Identification Software, Related to Figure 7

**Text S2**. Optimal-Coloring Software, Related to Figure 7

