## Supplementary figures and images for "NeuroPAL: A Neuronal Polychromatic Atlas of Landmarks for Whole-Brain Imaging in *C. elegans*"

### Figure S1

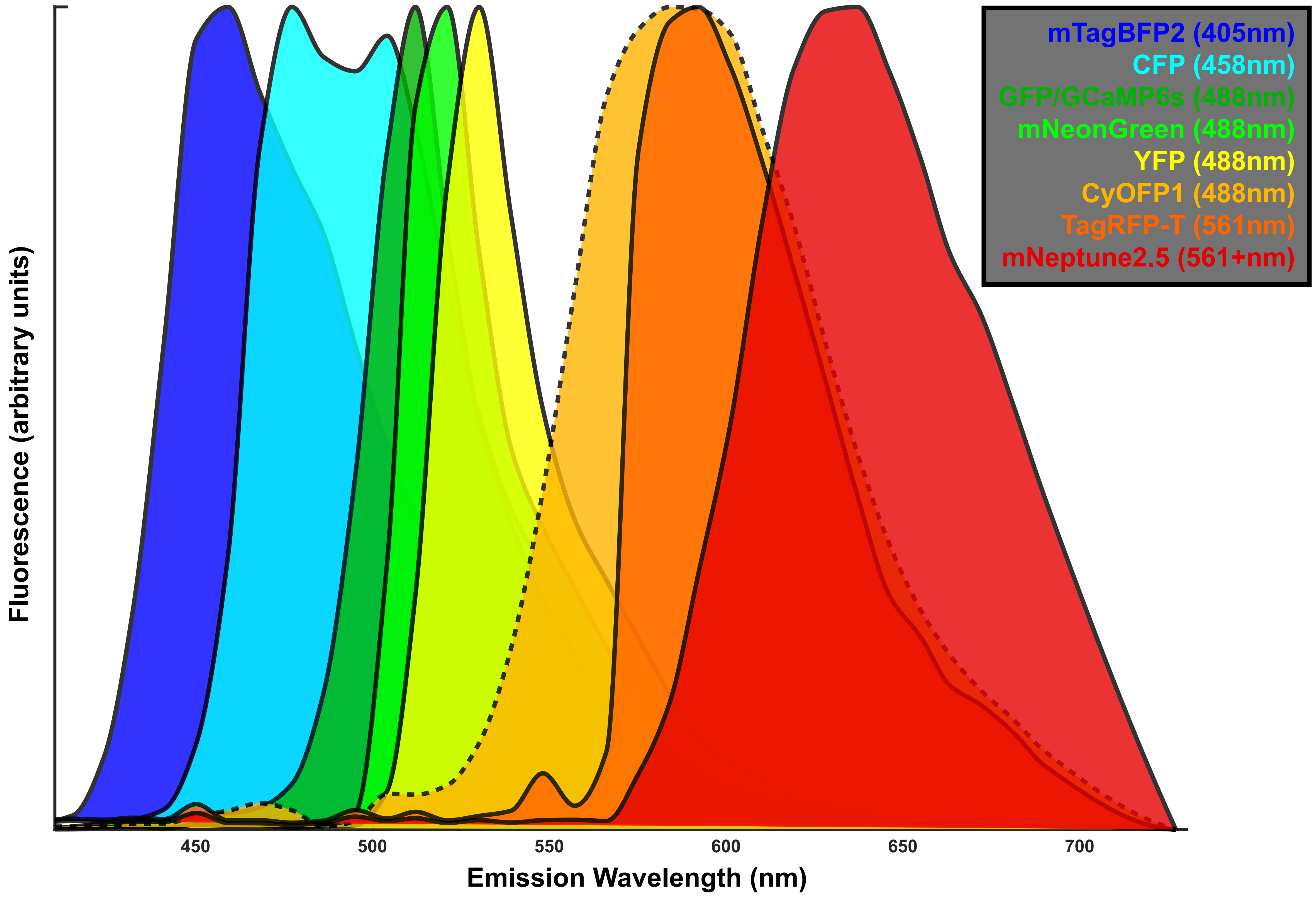

### Figure S2

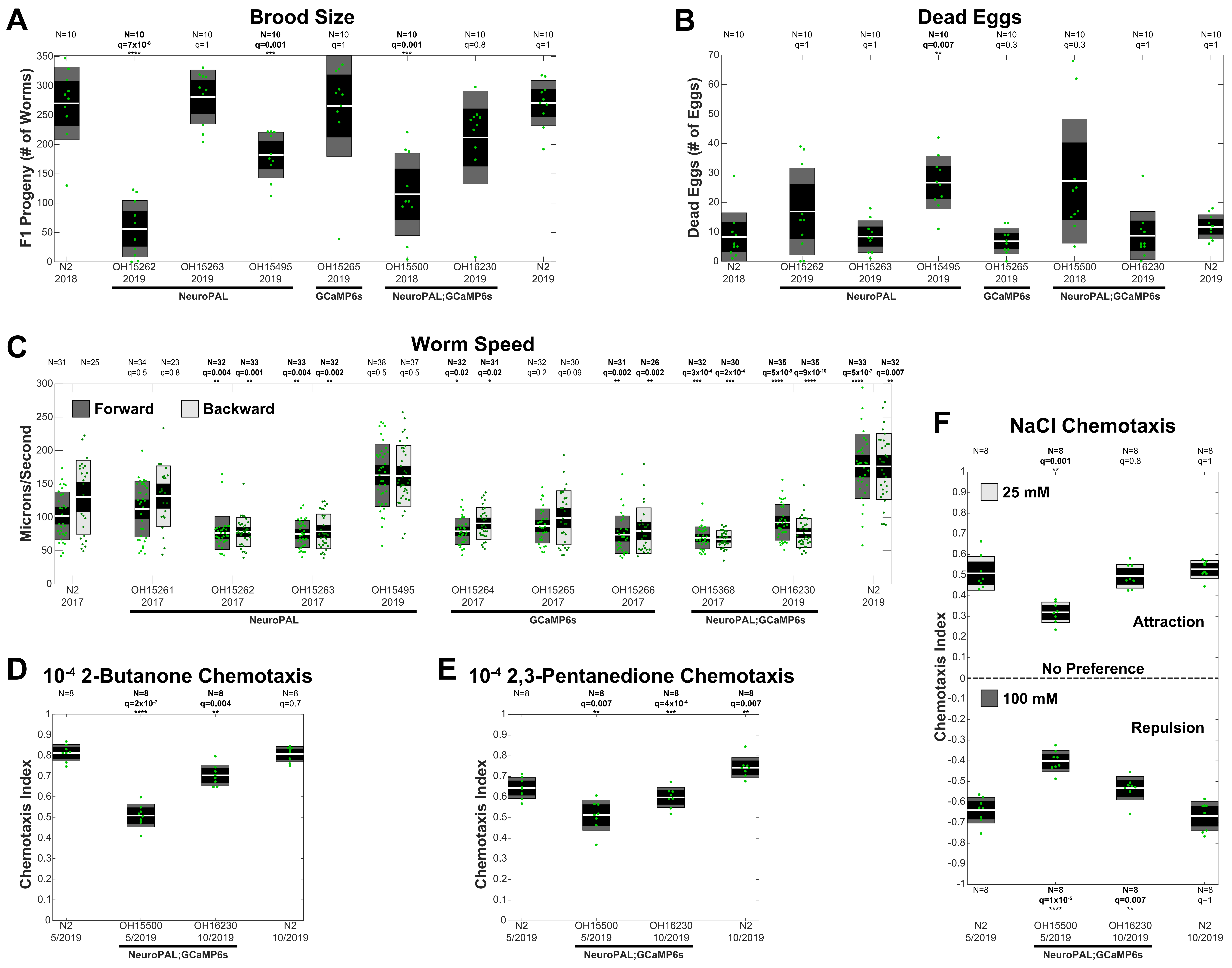

### Figure S3

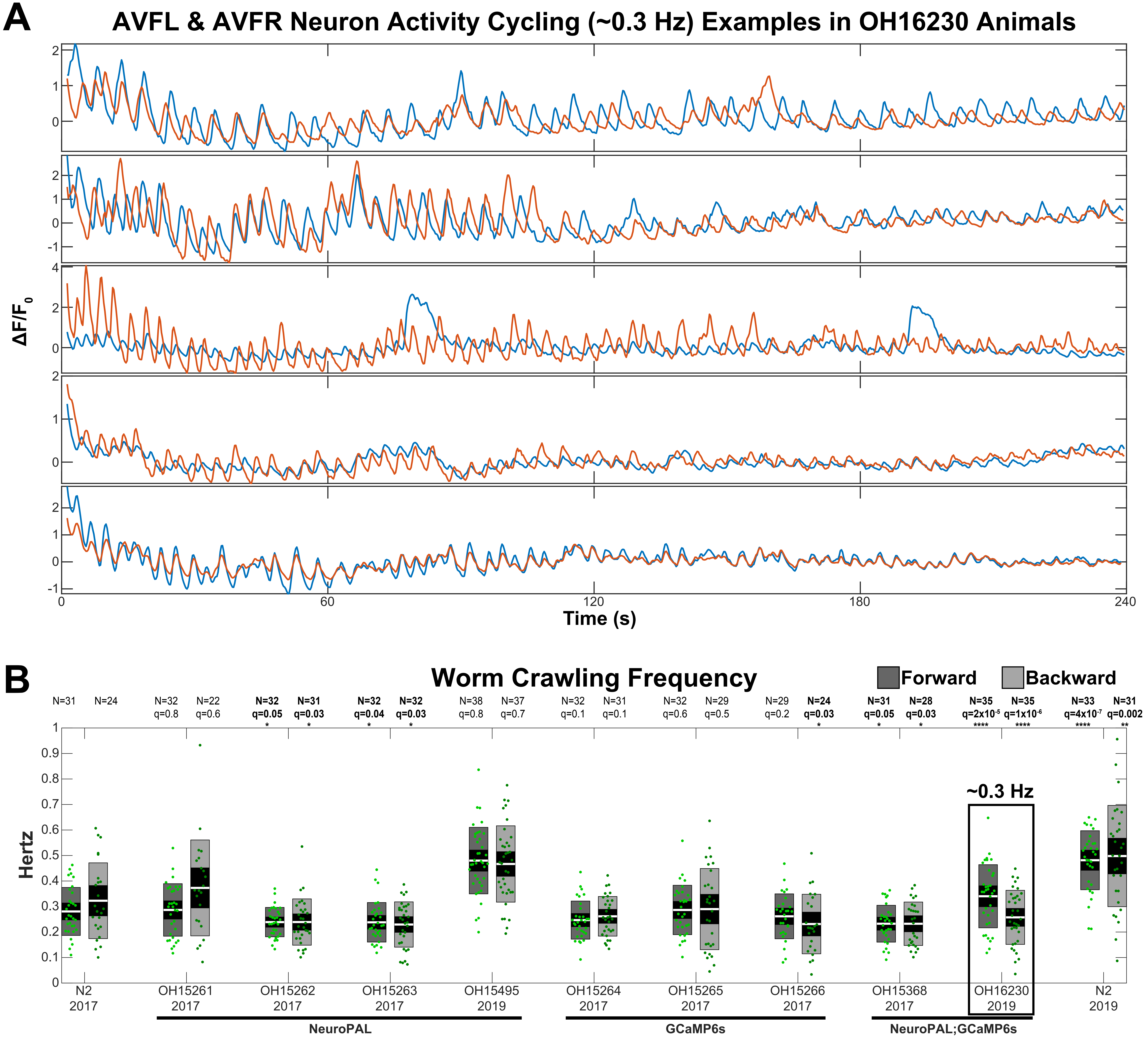

### Figure S4

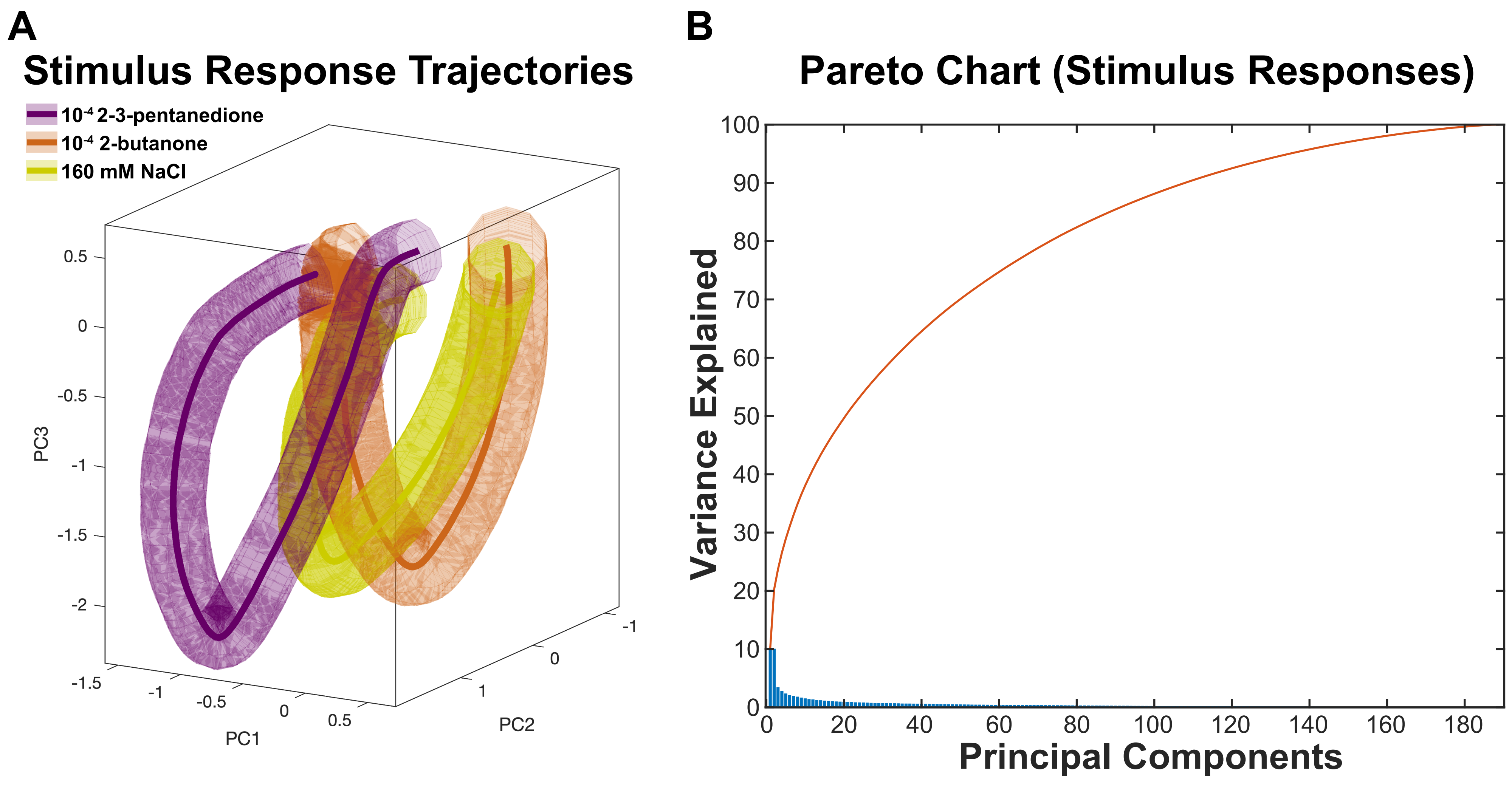

### Figure S5

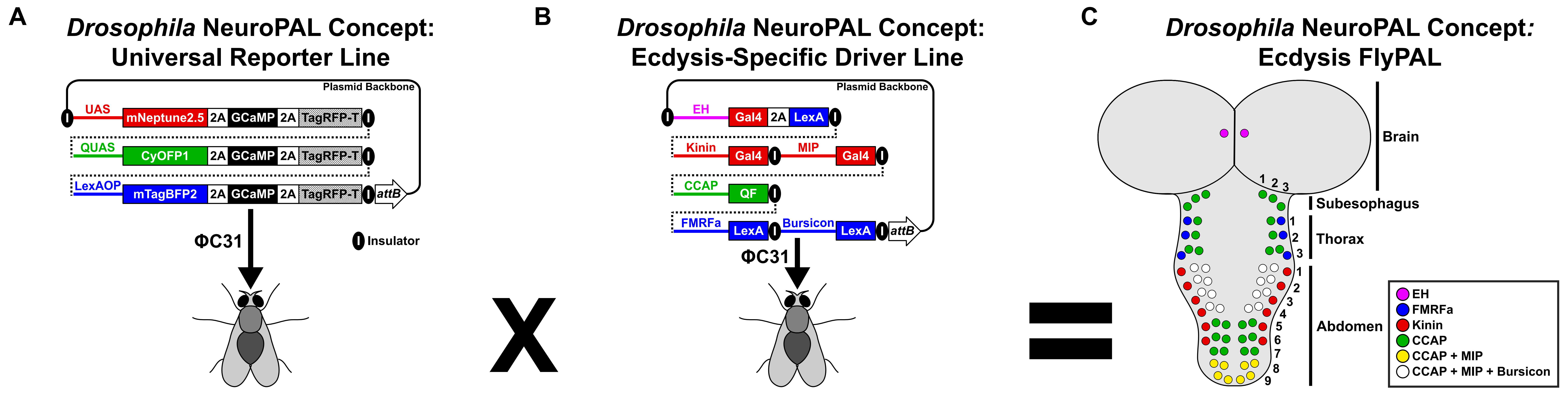
