## Supplemental Text S1 for "NeuroPAL: A Neuronal Polychromatic Atlas of Landmarks for Whole-Brain Imaging in *C. elegans*"

### Supplemental Text S1. Semi-Automated Neuronal Identification Software

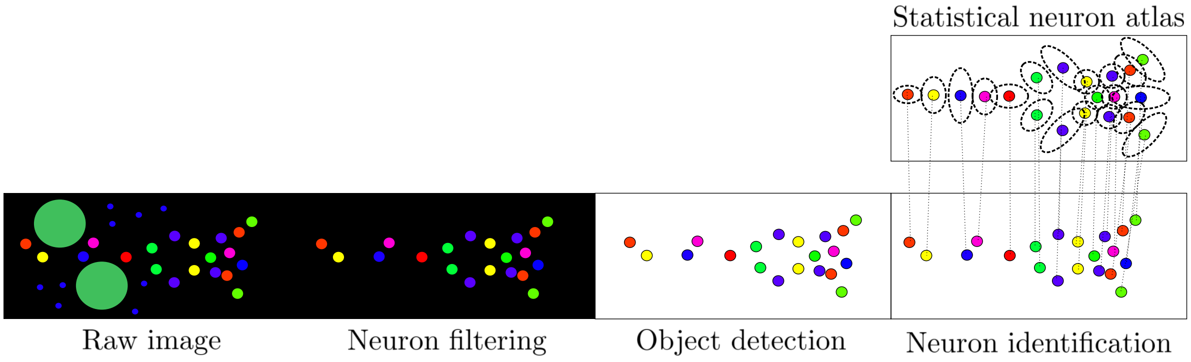

Figure 1.1: The schematic of the proposed *C. elegans* neuron detection and identification pipeline. From left to right: The raw image is supplied by the user and is then filtered to remove objects that are not neurons (i.e lysosomes, gut cells). From the filtered image, neurons are detected. The detected neurons are then aligned with a statistical atlas of neurons to yield their identities.

### 1 Introduction

In this manuscript, we describe a pipeline for automated and semi-automated identification of neurons in *C. elegans* captured in three-dimensional color images of NeuroPAL strains. The proposed pipeline effectively comprises the three steps: (1) filter, (2) detect, and (3) identify, illustrated in the schematic in figure 1.1 (re-drawn from Figure 8 in the main text).

The first step in the pipeline consists of pre-filtering regions in the raw images that correspond to non-neuronal structures such as gut cells and lysosomes. This is followed by the neurons being detected from the filtered image using a greedy sparse reconstruction procedure consisting of a variant of matching pursuit (Mallat and Zhang, 1993; Bergeaud and Mallat, 1995; Elad, 2010). Next, the color and positional features of the set of detected neurons are aligned with the features of a statistical atlas of neurons (Evangelidis and Horaud, 2018) to yield a set of likelihoods for each neuron which are then used to compute probabilistic assignment of identities (Mena et al., 2018). Lastly, and optionally, the proposed model allows user supervision to refine the identifications.

#### 1.1 Organization

This remainder of this note is organized as follows. First, we introduce notation in Table 1. Section 2 describes the filters for removing non-neuronal objects from the image. In section 3, we describe the matching pursuit procedure for detecting neurons. In section 4, we describe the training model for obtaining statistical worm atlases and in section 5 we show how to use the atlas to align and probabilistically identify neurons in out-of-sample worm images. In section 6 we demonstrate the empirical performance of our pipeline in cross-validation settings. Lastly, section 7 showcases some of the layouts and functions of the software package and graphical user interface.

### 2 Step 1: Neuron filter

Due to the use of multiples excitation lasers, auto-fluorescence from endogenous proteins, and even dirt accidentally captured on microscopy slides, worm images often contain bright non-neuronal components such as gut cells, lysosomes, and other fluorescent artifacts. To reduce the number of false positives in detection, we remove these structures in a pre-filtering step (Fig. 2.1).

These structures tend to be spatially segregated from neurons, and can therefore be easily removed manually. We use the *drawpolygon* tool in MATLAB to draw a polygon around these structures and then

| Variable | Definition | Space | Section |
|---|---|---|---|
| $\boldsymbol{I}$ | Microscopy image in $D$ voxels, $C$ colors | $\mathbb{R}^{D\times C}$ | 2,3 |
| $\boldsymbol{m}_k$ | center of $k$th detected neuron | $\mathbb{R}^3$ | 3,4 |
| $\boldsymbol{S}_k$ | covariance of gaussian envelope of $k$th detected neuron | $\mathbb{S}^3_{++}$ | 3 |
| $\boldsymbol{\pi}_k$ | color proportions of $k$th detected neuron | $\mathbb{R}^3$ | 3,4 |
| $\boldsymbol{\mu}_i$ | mean of position/color of neuron i | $\mathbb{R}^d$ | 4,5 |
| $\boldsymbol{\Sigma}_i$ | covariance of position/color of neuron i | $\mathbb{S}^d_{++}$ | 4,5 |
| $\boldsymbol{\beta}_j$ | Random transformation on worm j | $\mathbb{R}^{d\times d}$ | 4,5 |
| $\boldsymbol{\beta}^0_j$ | Random translation on worm j | $\mathbb{R}^d$ | 4,5 |
| $\boldsymbol{P}_j$ | Random permutation on worm j | $\mathcal{P}^{n\times n}$ | 4,5 |
| $\boldsymbol{z}_{i,j}$ | random draw of the stereotypical position/color of neuron i in worm j | $\mathbb{R}^d$ | 4 |
| $\boldsymbol{x}_{i,j}$ | randomly transformed neuron i for worm j | $\mathbb{R}^d$ | 4 |
| $\boldsymbol{y}_{i,j}$ | random permuted neuron i of worm j | $\mathbb{R}^d$ | 4, 5 |
| $\boldsymbol{L}$ | Likelihood matrix of $k$th detection corresponding to the $i$th atlas neuron | $\mathbb{R}^{k\times n}$ | 4,5 |

Table 1: Notation.

create a mask to remove them from the image; this process is an option in the graphical user interface detailed below and requires just a few seconds per worm of user input.

We also developed an alternative automatic removal method that works based on the following observations: gut cells are usually larger-radius balls or hollow rings (larger than regular neuron nuclei), with a green color, whereas lysosomes are usually small balls (or an irregular shape) and blue in color. Therefore we filter these components by size and color: we extract green and blue regions, and filter out those with size greater or less than the typical size of neuronal nuclei, respectively. For each color $c$, we create a binary mask to indicate the on-region for $c$. First, we lightly smooth each color channel individually (with a small 3d Gaussian filter). Then we generate a binary mask $m_c$ to indicate regions with intensity for color $c$ above the threshold (we use the 99th percentile as the threshold). Finally, we eliminate connected components in these masks that contain a total number of pixels above or below a user-defined threshold. The overall algorithm for filtering gut cells and lysosomes is sketched in algorithm 1.

Algorithm 1: Neuron filter - Pseudocode

**Input:** Microscopy image: $\boldsymbol{I} \in \mathbb{R}^{D\times C}$

Apply a Gaussian smoother in each color channel

Mask pixels in each color channel whose intensities are in the 99th percentile

Compute connected components in masks

Eliminate connected components that are green and are larger than $t'$ pixels (Gut elimination)

Eliminate connected components that are blue and are smaller than $t''$ pixels (Lysosome elimination)
**return** Filtered image: $\boldsymbol{I}_f \in \mathbb{R}^{D\times C}$

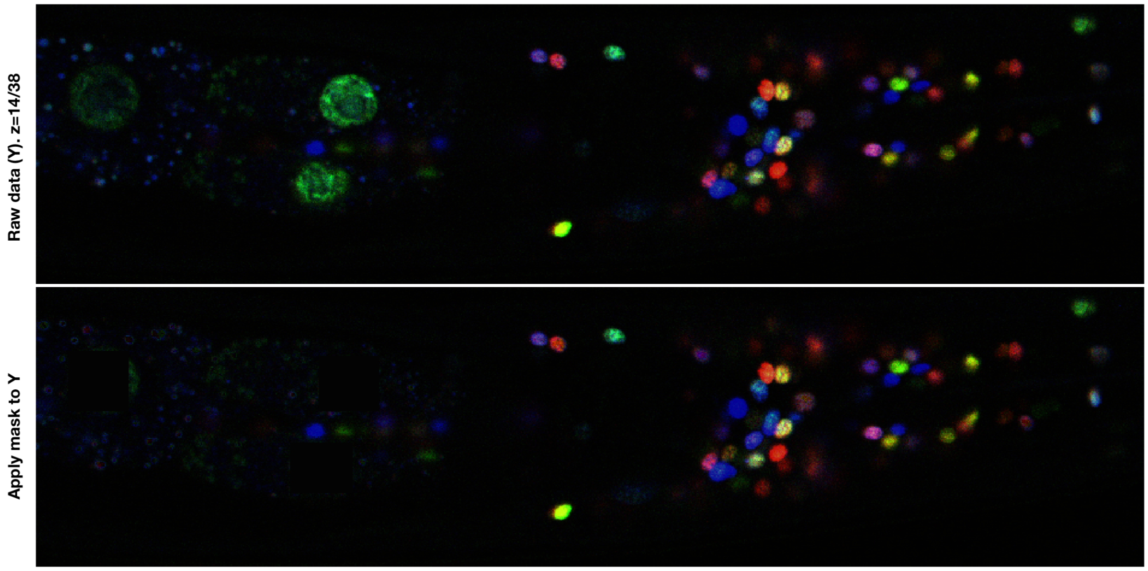

Figure 2.1: Filtered data (worm 38_YAaSP, ventral-dorsal view). Top: raw data (single z-slice). Bottom: filtered data, with with big green gut cells and small blue lysosomes largely removed. For all z slices, see Filter Video.

### 3 Step 2: Detecting neurons

#### 3.1 Nuclear shape model

In NeuroPAL worms, neuronal nuclei are labeled with different colors. To detect these nuclei we greedily minimize the following objective:

$$\left\| \boldsymbol{I}_f - \sum_{k=1}^{K} f_{\theta_k} \right\|_2^2, \tag{3.1}$$

where $\boldsymbol{I}_f \in \mathbb{R}^{D\times C}$ represents the 3-dimensional filtered color image of the worm ($D$ voxels, $C$ colors) and $f_{\theta_k}(\mathbf{v}) \in \mathbb{R}^{D\times C}$ is a function that approximates the multi-color shape of the neuron parametrized by $\theta_k$; $K$ is the number of cells visible in the field of view. We choose $f$ to be a truncated Gaussian function:

$$f_{\theta_k}(\mathbf{v}) = (2\pi)^{-\frac{3}{2}}|\Sigma|^{-\frac{1}{2}} \exp\left[\frac{-(\mathbf{v}-\boldsymbol{m}_k)^T \boldsymbol{S}_k^{-1}(\mathbf{v}-\boldsymbol{m}_k)}{2}\right]; \tag{3.2}$$

$$f_{\theta_k}(\mathbf{v}, c) = b_k^c + \pi_k^c f_{\theta_k}(\mathbf{v}) 1_{[f_{\theta_k}(\mathbf{v})>\boldsymbol{t}_k]} \tag{3.3}$$

here $\boldsymbol{m}_k$ is the spatial mean of the Gaussian bump, $\boldsymbol{S}_k$ controls the size and eccentricity of the Gaussian, $\boldsymbol{t}_k$ controls the amount of truncation of the Gaussian, $\pi_k^c$ controls the brightness of the cell in the $c$-th color channel, and $b_k^c$ is the brightness of the local background in color channel $c$, all for the cell indexed by $k$. We collect these parameters into

$$\theta_k = \{b_k, \pi_k, \boldsymbol{m}_k, \boldsymbol{S}_k, \boldsymbol{t}_k\},$$

and impose constraints on each of these parameters:

$$\mathfrak{C} = \begin{cases} \boldsymbol{\pi}_{\min} \leq \boldsymbol{\pi}_k \leq \boldsymbol{\pi}_{\max} \\ b_{\min} \leq b_k \leq b_{\max} \\ \boldsymbol{m}_{\min} \leq \boldsymbol{m}_k \leq \boldsymbol{m}_{\max} \\ \sigma_{\min} \leq \sigma(\boldsymbol{S}_k) \leq \sigma_{\max} \\ \boldsymbol{t}_{\min} \leq \boldsymbol{t}_k \leq \boldsymbol{t}_{\max}, \end{cases}$$

where the $\sigma(S)$ operator returns the singular values of the matrix $S$.

#### 3.2 Optimization

To optimize this objective, we use a matching pursuit (MP) based strategy. In standard MP (Elad, 2010) we begin with a finite collection of filter functions $f_k$ and then greedily add in the filter element $k$ that leads to the largest reduction in the squared error (3.1). This greedy optimization can be implemented efficiently using convolutions and simple subtractive updates of the objective function, until a maximal number $K$ filters have been added to the model.

In the present context, the family of filters represented by the constraint set $\mathfrak{C}$ above is infinitely large, so we can not apply the standard MP approach directly. Instead, we choose an "average" Gaussian shape (i.e., with covariance $\boldsymbol{S}$ chosen near the "middle" of the constraint set), convolve the image $\boldsymbol{I}_f$ with this shape, and find local optima of the resulting function. (This is analogous to the first step of standard MP.) Then instead of subtracting this "average" shape away from around the peaks located in the first step, we locally optimize the parameter $\theta_k$ to fit the local shape of the image and then subtract this locally-optimized shape away. Then we iterate, adding a new locally-optimized shape to the model at each iteration $k$. (This algorithm can be parallelized by finding multiple local optima $\mu_k$ in the first step and then updating the corresponding parameters $\theta_k$ in parallel for locations $\mu_k$ that are sufficiently far apart.)

If we initialize $\rho_c = G^T \boldsymbol{I}_{f,c}$ (with $\boldsymbol{I}_{f,c}$ the 3d data volume in color channel $c$, and $G^T \boldsymbol{I}_{f,c}$ denoting convolution of $\boldsymbol{I}_{f,c}$ with the average Gaussian shape), then our approach can be summarized as in algorithm 2; see figure 3.1 for an illustration.

In the current optimization procedure there is one main hyperparameter that should be specified before running the optimization: the number of neurons ($K$). We found it convenient to set $K$ to a value larger than the estimated number of neurons in the image to reduce false negatives. This approach results in the detection of some non-neuronal objects that can further be filtered out by the user. To avoid over-segmentation we incorporated an exclusion radius to assure that every pair of the detected neurons have a distance of at least 1.5 $\mu m$. Moreover, to further reduce false positives we subtract the objects that are detected by the algorithm but have a very small size, without accepting them as neurons.

Finally, we emphasize that the above approach is *unsupervised*: we do not require the user to label any neurons to train the model. However, given a sufficient corpus of curated labeled data, we expect supervised or semi-supervised approaches to lead to improved accuracy. Due to the modularity of our overall pipeline, it will be easy to drop in a new detection step here if desired before proceeding to the neural identification step described in the next section.

Algorithm 2: Neuron-like object detection

**Input:** Filtered microscopy image: $\boldsymbol{I}_f \in \mathbb{R}^{D \times C}$

Initialize $\rho_c = G^T \boldsymbol{I}_{f,c}$

**for** $k = 1, \ldots, K$ **do**

Choose $\mu_k = \underset{x}{\operatorname{argmax}}\{\max_c \rho_c(x)\}$ - regional maximum of smoothed image

Set $\boldsymbol{m}_{min}$ and $\boldsymbol{m}_{max}$ in the constraint set $\mathfrak{C}$ to enclose a small box near this initial $\mu_k$, and then optimize

$$\theta_k = \underset{\theta_k \in \mathfrak{C}}{\operatorname{argmin}} \left\| \boldsymbol{I}_f - \sum_{j=1}^{k} f_{\theta_j}(X) \right\|^2$$

Compute residual image: $\rho_c \leftarrow \rho_c - G^T f_{\theta_k}$.

**end for**

**return** $\theta_k = \{b_k, \boldsymbol{\pi}_k, \boldsymbol{m}_k, \boldsymbol{S}_k, \boldsymbol{t}_k\}$ which are the background mean, color proportions, cell centers, and shape described by covariance and truncation value.

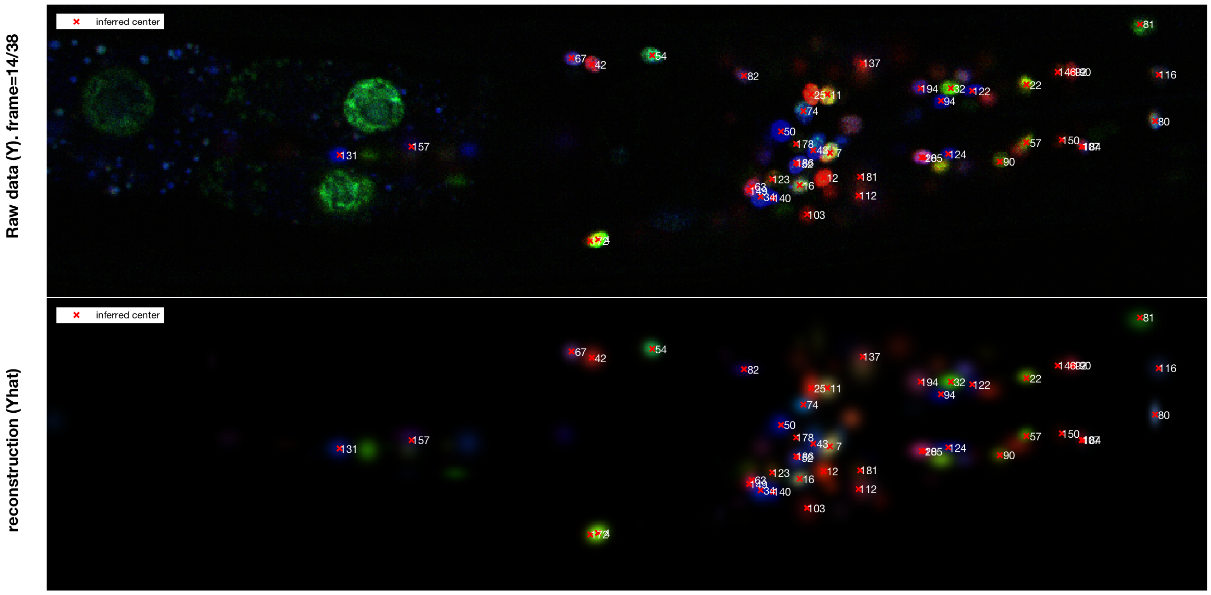

Figure 3.1: Detection step run on filtered data (same worm and z-slice as Fig. 2.1). Top: raw data $\boldsymbol{I}$. Bottom: reconstruction $\sum_k f_{\theta_k}$. Red crosses indicate inferred location of neurons; numbers indicate the order in which these locations were detected (typically brighter cells are detected first; note that some visible cells may be brighter on different $z$ planes and may therefore not be labeled in this plane). For filtered data of all z slices, see full reconstruction video and residue video.

### 4 Step 3a: Statistical neuron atlas construction

Due to variability in illumination and the pose of the worm when imaged, observed neuron positions and their exact color balance may vary across imaged worms. This presents a significant challenge when attempting to obtain correspondences between worms to infer the identities of neurons. Therefore, to normalize the random variability that occurs across different worms, prior to identifying neurons in any given microscopy image, we estimate a statistical atlas of neuron positions and colors.

The approach we take resembles the joint expectation-maximization alignment of point sets technique of Evangelidis and Horaud (2018), with several important differences discussed below. The dataset we are modeling consists of a collection of point sets: each worm corresponds tod one point set, with each point in the set corresponding to the position and color of a single detected neuron. We model each of these positions and colors as samples from a statistical atlas that is common across worms. Each neuron $i$ has a corresponding mean and covariance in this atlas, denoted as $\boldsymbol{\mu}_i$ and $\boldsymbol{\Sigma}_i$, respectively. After drawing all the positions and colors for a given worm $j$ we apply a random affine transformation (parametrized by a matrix $\boldsymbol{\beta}_j$ and translation vector $\boldsymbol{\beta}_j^0$). Finally, since the order of neurons in each point set is arbitrary, we scramble the identities of the neurons with a random permutation, parameterized by a permutation matrix $\boldsymbol{P}_j$. This generative model is summarized in Figure 4.1. See also (Bubnis et al., 2019) for a related model (without the alignment term, and with an inference approach that differs from the methods we describe below).

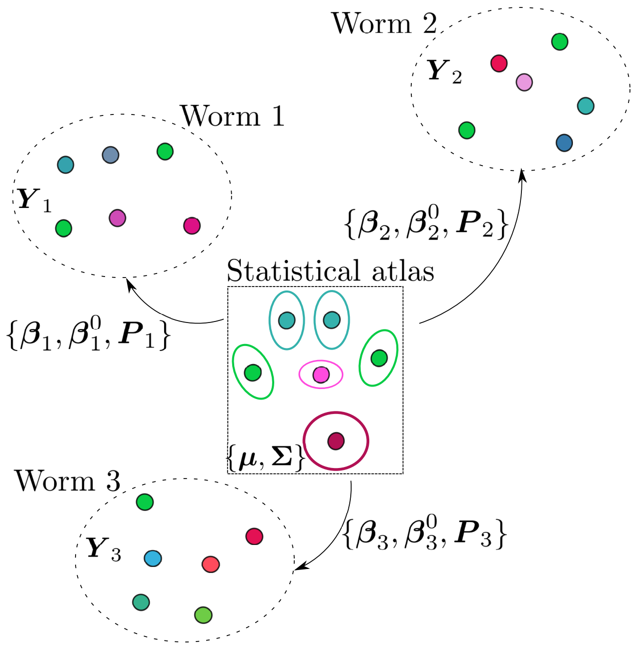

Figure 4.1: Schematic of the generative model of neuron position and color expression. First we draw a position and color for each neuron $i$ from a distribution with mean $\boldsymbol{\mu}_i$ and covariance $\boldsymbol{\Sigma}_i$ (center box); then, to create the observed data $\boldsymbol{y}_{i,j}$ (the color and position of the $i$-th neuron of the $j$-th worm) we apply a random affine transformation $\boldsymbol{\beta}_j, \boldsymbol{\beta}_j^0$ to the positions and colors and a random permutation $\boldsymbol{P}_j$ to the identities (indicated with the arrows to the observed datasets $\mathbf{Y}_j$ for each worm $j$).

Evangelidis and Horaud (2018) infer the parameters of this generative model (i.e., the means and covariances of the statistical atlas, the random transformations, and the random permutations) in a completely unsupervised fashion using a three-way expectation maximization procedure. However, in our dataset, we have access to fully annotated neuron detections. We take advantage of this supervised data to simplify the inference problem.

Now we can describe our model in detail. Neural positions are three-dimensional, and there are three color channels in this dataset; therefore, if we use $\mathbf{y}_{i,j}$ to denote the appended position and color vector of the $i$-th neuron in worm $j$ (as output by the detection step described in the previous section), then $\mathbf{y}_{i,j} \in \mathbb{R}^6$. Each of these observed $\mathbf{y}_{i,j}$ vectors has a corresponding latent vector $z_{i,j}$ in the aligned atlas space. We model this latent vector as Gaussian,

$$z_{i,j} \sim \mathcal{N}(\boldsymbol{\mu}_i, \boldsymbol{\Sigma}_i), \tag{4.1}$$

with means $\boldsymbol{\mu}_i \in \mathbb{R}^6$ and covariances $\boldsymbol{\Sigma}_i \in \mathbb{S}^6_+$ that do not depend on the worm index $j$. We model the covariance $\boldsymbol{\Sigma}_i$ with block structure of the form $\boldsymbol{\Sigma}_i = \begin{bmatrix} \Sigma^i_{\text{position}} & \mathbf{0} \\ \mathbf{0} & \Sigma^i_{\text{color}} \end{bmatrix}$, since position and each color are independently varying.

Now the latent vectors $z_{i,j}$ in the atlas space and observed data $\mathbf{y}_{i,j}$ extracted from the imaged worm $j$ are connected by a worm-specific random affine transformation and permutation. We denote the intermediate affine-transformed variables as $x_{i,j}$:

$$x_{i,j} = z_{i,j}\boldsymbol{\beta}_j + \boldsymbol{\beta}^0_j, \tag{4.2}$$

with $\boldsymbol{\beta}_j$ a $6 \times 6$ matrix (with a similar block structure as $\boldsymbol{\Sigma}_i$) and $\boldsymbol{\beta}^0_j \in \mathbb{R}^6$. then we obtain $\mathbf{y}_{i,j}$ by scrambling the labels via the permutation $p_j$ (corresponding to a permutation matrix $\mathbf{P}_j$):

$$y_{i,j} = x_{p_j(i),j}. \tag{4.3}$$

Given these modelling assumptions, for a dataset of $m$ worms and $n_j$ detected neurons in each worm, we can express the likelihood as:

$$P(\boldsymbol{y}|\boldsymbol{\mu}, \boldsymbol{\Sigma}, \boldsymbol{P}, \boldsymbol{\beta}) = \tag{4.4}$$

$$\prod_{j=1}^{m}\prod_{i=1}^{n_j} \frac{1}{(2\pi)^{d/2}\det((\boldsymbol{\beta}_j\boldsymbol{\Sigma}_{p_{i,j}}\boldsymbol{\beta}_j^T))^{1/2}} e^{-(1/2)(y_{i,j}-\mu_{p_{i,j}}\beta_j-\beta^0_j)(\beta_j\Sigma_{p_{i,j}}\beta_j^T)^{-1}(y_{i,j}-\mu_{p_{i,j}}\beta_j-\beta^0_j)^T} \tag{4.5}$$

which has the negative log likelihood:

$$-L(\boldsymbol{y}|\boldsymbol{\mu}, \boldsymbol{\Sigma}, \boldsymbol{P}, \boldsymbol{\beta}) = \tag{4.6}$$

$$\sum_{j=1}^{m}\sum_{i=1}^{n_j} \frac{1}{2}(y_{i,j} - \mu_{p_{i,j}}\boldsymbol{\beta}_j - \boldsymbol{\beta}^0_j)(\boldsymbol{\beta}_j\boldsymbol{\Sigma}_{p_{i,j}}\boldsymbol{\beta}_j^T)^{-1}(y_{i,j} - \mu_{p_{i,j}}\boldsymbol{\beta}_j - \boldsymbol{\beta}^0_j)^T) +$$

$$\frac{1}{2}\log\det((\boldsymbol{\beta}_j\boldsymbol{\Sigma}_{p_{i,j}}\boldsymbol{\beta}_j^T)) + C \tag{4.7}$$

Since the term $\sum_j \sum_i (1/2)\log\det((\boldsymbol{\beta}_j\boldsymbol{\Sigma}_{p_{i,j}}\boldsymbol{\beta}_j^T)$ is permutation invariant, we can write it as $\sum_j \sum_i (1/2)\log\det((\boldsymbol{\beta}_j\boldsymbol{\Sigma}_i\boldsymbol{\beta}_j^T)$.

Therefore, the maximum likelihood estimate (MLE) for our generative model involves optimizing the following objective:

$$\underset{\boldsymbol{P},\boldsymbol{\beta},\boldsymbol{\mu},\boldsymbol{\Sigma}}{\text{minimize}} \tag{4.8}$$

$$\sum_{j=1}^{m}\sum_{i=1}^{n_j}(y_{i,j} - \mu_{p_{i,j}}\boldsymbol{\beta}_j - \boldsymbol{\beta}^0_j)(\boldsymbol{\beta}_j\boldsymbol{\Sigma}_{p_{i,j}}\boldsymbol{\beta}_j^T)^{-1}(y_{i,j} - \mu_{p_{i,j}}\boldsymbol{\beta}_j - \boldsymbol{\beta}^0_j)^T) \tag{4.9}$$

$$+\sum_{j=1}^{m}\sum_{i=1}^{n_j}\log\det((\boldsymbol{\beta}_j\boldsymbol{\Sigma}_i\boldsymbol{\beta}_j^T). \tag{4.10}$$

#### 4.1 Optimization

To infer the parameters of the generative model, we take an iterative block-coordinate descent approach, similar to Evangelidis and Horaud (2018): we fix $(\boldsymbol{P}, \boldsymbol{\beta}, \boldsymbol{\beta}_0)$ (with $\boldsymbol{P}$ abbreviating the collection of permutations $\boldsymbol{P}_j$ for all worms $j$, and similarly for $\boldsymbol{\beta}, \boldsymbol{\beta}_0$) and solve for $(\boldsymbol{\mu}, \boldsymbol{\Sigma})$, then fix $(\boldsymbol{\mu}, \boldsymbol{\Sigma})$ and solve for $(\boldsymbol{P}, \boldsymbol{\beta}, \boldsymbol{\beta}_0)$.

##### 4.1.1 Inference of the statistical atlas parameters $\boldsymbol{\mu}, \boldsymbol{\Sigma}$

Let $\boldsymbol{P}_j \in \mathcal{P}^{n\times n}$ denote the permutation matrix, $\boldsymbol{Y}_j = [\mathbf{y}_{1,j}^T \dots \mathbf{y}_{n,j}^T]^T \in \mathbb{R}^{n\times d}$ denote the row stacked features of the neurons of the jth worm, and let $\boldsymbol{\mu} = [\boldsymbol{\mu}_1^T \dots \boldsymbol{\mu}_n^T] \in \mathbb{R}^{n\times d}$ denote the row stacked neuron means.

The generative model can be written in matrix form as:

$$\boldsymbol{Y}_j = \boldsymbol{P}_j \boldsymbol{\mu} \boldsymbol{\beta}_j + \mathbf{1}\boldsymbol{\beta}_j^0 + \boldsymbol{E} \tag{4.11}$$

where $\boldsymbol{E}_i \sim \mathcal{N}(\mathbf{0}, \boldsymbol{\beta}_j \boldsymbol{\Sigma}_{\boldsymbol{P}_{i,j}} \boldsymbol{\beta}_j^T)$ denotes the row stacked uncertainty terms.

Since $\boldsymbol{P}_j^T \boldsymbol{P}_j = \mathbf{I}$ because $\boldsymbol{P}$ is a permutation matrix and assuming that $\boldsymbol{\beta}_j$ is a non-degenerate transformation, its inverse exists and can be used to write the system as:

$$\boldsymbol{P}_j^T \boldsymbol{Y}_j \boldsymbol{\beta}_j^{-1} - \mathbf{1}\boldsymbol{\beta}_j^0 \boldsymbol{\beta}_j^{-1} = \boldsymbol{\mu} + \boldsymbol{V} \tag{4.12}$$

where $\boldsymbol{V}_i \sim \mathcal{N}(\mathbf{0}, \boldsymbol{\Sigma}_i)$ is a term to quantify uncertainty.

This equation can be used to infer $\boldsymbol{\mu}$ and $\boldsymbol{\Sigma}$ by computing the first and second moments of $\boldsymbol{V}$:

$$\boldsymbol{\mu}^* = \frac{1}{m}\sum_{j=1}^{m} \boldsymbol{P}_j^T \boldsymbol{Y}_j \boldsymbol{\beta}_j^{-1} - \mathbf{1}\boldsymbol{\beta}_j^0 \boldsymbol{\beta}_j^{-1} \tag{4.13}$$

$$\boldsymbol{\Sigma}_i^* = \frac{1}{m}\sum_{j=1}^{m} (\boldsymbol{P}_{j,i}^T \boldsymbol{Y}_j \boldsymbol{\beta}_j^{-1} - \boldsymbol{\beta}_j^0 \boldsymbol{\beta}_j^{-1} - \boldsymbol{\mu}_i)^T (\boldsymbol{P}_{j,i}^T \boldsymbol{Y}_j \boldsymbol{\beta}_j^{-1} - \boldsymbol{\beta}_j^0 \boldsymbol{\beta}_j^{-1} - \boldsymbol{\mu}_i) \tag{4.14}$$

##### 4.1.2 Inference of the transformation terms $\boldsymbol{\beta}, \boldsymbol{\beta}_0$

We can infer the transformation and translation terms $\boldsymbol{\beta}, \boldsymbol{\beta}_0$ if we view equation 4.12 as a weighted linear regression with a different goodness of fit criteria for each neuron quantified by a mahalanobis norm using the statistical atlas covariance $\boldsymbol{\Sigma}_i$.

The weighted regression can be posed as the following unconstrained minimization problem:

$$\underset{\boldsymbol{\beta}_j^{-1}, \boldsymbol{\beta}_j^0 \boldsymbol{\beta}_j^{-1}}{\text{minimize}} \sum_{i=1}^{n_j} (\boldsymbol{P}_{j,i}^T \boldsymbol{Y}_j \boldsymbol{\beta}_j^{-1} - \boldsymbol{\beta}_j^0 \boldsymbol{\beta}_j^{-1} - \boldsymbol{\mu}_i) \boldsymbol{\Sigma}_i^{-1} (\boldsymbol{P}_{j,i}^T \boldsymbol{Y}_j \boldsymbol{\beta}_j^{-1} - \boldsymbol{\beta}_j^0 \boldsymbol{\beta}_j^{-1} - \boldsymbol{\mu}_i)^T$$

which has the derivatives wrt to $\boldsymbol{\beta}_j^{-1}, \boldsymbol{\beta}_j^0 \boldsymbol{\beta}_j^{-1}$ as

$$\frac{\partial}{\partial \boldsymbol{\beta}_j^{-1}} = \sum_{i=1}^{n_j} (\boldsymbol{P}_{j,i}^T \boldsymbol{Y}_j)^T (\boldsymbol{P}_{j,i}^T \boldsymbol{Y}_j \boldsymbol{\beta}_j^{-1} - \boldsymbol{\beta}_j^0 \boldsymbol{\beta}_j^{-1} - \boldsymbol{\mu}_i) \boldsymbol{\Sigma}_i^{-1} = 0$$

$$\frac{\partial}{\partial \boldsymbol{\beta}_j^0 \boldsymbol{\beta}_j^{-1}} = \sum_{i=1}^{n_j} (\boldsymbol{P}_{j,i}^T \boldsymbol{X}_j \boldsymbol{\beta}_j^{-1} - \boldsymbol{\beta}_j^0 \boldsymbol{\beta}_j^{-1} - \boldsymbol{\mu}_i) \boldsymbol{\Sigma}_i^{-1} = 0$$

Algorithm 3: Affine transformation optimization

**Input:** Statistical atlas parameters $\boldsymbol{\mu}, \boldsymbol{\Sigma}$, neuron correspondences for $j$th worm $\boldsymbol{P}_j$, $\epsilon$ convergence tolerance
**Initalize:** $[\boldsymbol{\beta}_j^{-1}]^0$ and $[\boldsymbol{\beta}_j^0\boldsymbol{\beta}_j^{-1}]^0$ randomly
**while** Not converged **do**

$[\boldsymbol{\beta}_j^0\boldsymbol{\beta}_j^{-1}]^t \leftarrow \Big(\sum_{i=1}^n (\boldsymbol{P}_{j,i}^T\boldsymbol{Y}_j[\boldsymbol{\beta}_j^{-1}]^{t-1} - \boldsymbol{\mu}_i)\boldsymbol{\Sigma}_i^{-1}\Big)\Big(\sum_{i=1}^n \boldsymbol{\Sigma}_i^{-1}\Big)^{-1}$

$[\boldsymbol{\beta}_j^{-1*}]^t \leftarrow \text{reshape}\Big(\Big(\sum_{i=1}^n (\boldsymbol{\Sigma}_i^{-1} \otimes (\boldsymbol{P}_{j,i}^T\boldsymbol{Y}_j)^T(\boldsymbol{P}_{j,i}^T\boldsymbol{Y}_j))\Big)^{-1}\Big(\sum_{i=1}^n \text{vec}((\boldsymbol{P}_{j,i}^T\boldsymbol{Y}_j)^T([\boldsymbol{\beta}_j^0\boldsymbol{\beta}_j^{-1}]^t + \boldsymbol{\mu}_i)\boldsymbol{\Sigma}_i^{-1})\Big)\Big)$

Check convergence $\|[\boldsymbol{\beta}_j^{-1}]^t - [\boldsymbol{\beta}_j^{-1}]^{t-1}\|_F \leq \epsilon$

**end while**
**return** $\boldsymbol{\beta}_j^* = ([\boldsymbol{\beta}_j^{-1}]^t)^{-1}, \boldsymbol{\beta}_j^{0*} = [\boldsymbol{\beta}_j^0\boldsymbol{\beta}_j^{-1}]^t\boldsymbol{\beta}_j^*$

First we solve for $\boldsymbol{\beta}_j^0\boldsymbol{\beta}_j^{-1}$:

$$\boldsymbol{\beta}_j^0\boldsymbol{\beta}_j^{-1*} = \Big(\sum_{i=1}^n (\boldsymbol{P}_{j,i}^T\boldsymbol{Y}_j\boldsymbol{\beta}_j^{-1} - \boldsymbol{\mu}_i)\boldsymbol{\Sigma}_i^{-1}\Big)\Big(\sum_{i=1}^n \boldsymbol{\Sigma}_i^{-1}\Big)^{-1} \tag{4.15}$$

To analytically solve for $\boldsymbol{\beta}_j^{-1}$, we utilize the fact that $\text{vec}(ABC) = (C^T \otimes A)\text{vec}(B)$ where $\text{vec}(\cdot)$ denotes the vectorization operation and $\otimes$ denotes Kronecker product. This yields the following relation:

$$\sum_{i=1}^n (\boldsymbol{\Sigma}_i^{-1} \otimes (\boldsymbol{P}_{j,i}^T\boldsymbol{Y}_j)^T(\boldsymbol{P}_{j,i}^T\boldsymbol{Y}_j))\text{vec}(\boldsymbol{\beta}_j^{-1}) = \sum_{i=1}^n \text{vec}((\boldsymbol{P}_{j,i}^T\boldsymbol{Y}_j)^T(\boldsymbol{\beta}_j^0\boldsymbol{\beta}_j^{-1} + \boldsymbol{\mu}_i)\boldsymbol{\Sigma}_i^{-1})$$

which gives a vectorized solution for $\boldsymbol{\beta}_j^{-1}$:

$$\text{vec}(\boldsymbol{\beta}_j^{-1*}) = \Big(\sum_{i=1}^n (\boldsymbol{\Sigma}_i^{-1} \otimes (\boldsymbol{P}_{j,i}^T\boldsymbol{Y}_j)^T(\boldsymbol{P}_{j,i}^T\boldsymbol{Y}_j))\Big)^{-1}\Big(\sum_{i=1}^n \text{vec}((\boldsymbol{P}_{j,i}^T\boldsymbol{Y}_j)^T(\boldsymbol{\beta}_j^0\boldsymbol{\beta}_j^{-1} + \boldsymbol{\mu}_i)\boldsymbol{\Sigma}_i^{-1})\Big) \tag{4.16}$$

Note that the expressions for the zero gradient solutions of $\boldsymbol{\beta}_j^{-1}$ involves $\boldsymbol{\beta}_j^0\boldsymbol{\beta}_j^{-1}$ and likewise, the zero gradient solution of $\boldsymbol{\beta}_j^0\boldsymbol{\beta}_j^{-1}$ involves $\boldsymbol{\beta}_j^{-1}$. Therefore we employ an iterative coordinate descent method to yield an optimum solution for $\boldsymbol{\beta}_j^{-1}$ and $\boldsymbol{\beta}_j^0\boldsymbol{\beta}_j^{-1}$. These optima can then be used to infer $\boldsymbol{\beta}_j^*$ and $\boldsymbol{\beta}_j^{0*}$. The affine transformation optimization procedure is outlined in algorithm 3.

##### 4.1.3 Permutation inference

Lastly, we can solve for the permutation $\boldsymbol{P}_j$ by setting up a $n \times n$ transport matrix $\boldsymbol{D}$ where

$$\boldsymbol{D}_{u,v} = (\boldsymbol{\mu}_u\boldsymbol{Y}\boldsymbol{\beta}_j + \boldsymbol{\beta}_j^0 - \boldsymbol{Y}_{j,v})\boldsymbol{\Sigma}_u^{-1}(\boldsymbol{\mu}_u\boldsymbol{Y}\boldsymbol{\beta}_j + \boldsymbol{\beta}_j^0 - \boldsymbol{Y}_{j,v})^T \tag{4.17}$$

and obtaining $\boldsymbol{P}_j$ through the linear assignment optimization solved through the Hungarian algorithm of Kuhn (1955) which minimizes the following objective:

$$\boldsymbol{P}_j^* = \arg\min_{p\in\mathcal{P}} \sum_{u,v} p_{u,v}\boldsymbol{D}_{u,v}. \tag{4.18}$$

#### 4.2 Statistical atlas of neuron positions and colors

If we have access to several annotated worms, meaning that we have access to both the detections and their correspondences to the identities of neurons, $\boldsymbol{P}$, we can infer the transformation terms $\{\boldsymbol{\beta}, \boldsymbol{\beta}_0\}$ as well as the parameters of the statistical atlas $\{\boldsymbol{\mu}, \boldsymbol{\Sigma}\}$ using the procedure outlined in algorithm 4.

In words, algorithm 4 operates in the following way. First, the targeted inference parameters are initialized using the neuron centers and colors for a random worm. Then, the remaining worms are affinely aligned to the hypothetical atlas by solving the linear system for $\{\boldsymbol{\beta}_k, \boldsymbol{\beta}_j^0\}$ in equation 4.16. The means and covariances of the aligned neurons are then used to update the atlas parameters $\boldsymbol{\mu}$ and $\boldsymbol{\Sigma}$. This procedure is iteratively repeated until convergence. The resulting statistical atlas consisting of the mean stereotypical neuron positions and colors and their covariances can be seen in Figure 3 in the main text and Fig. 6.2 below.

---

Algorithm 4: Train statistical neuron atlas

**Input:** $\{\mathbf{y}_{i,j}\}$ (colors and positions) and $\{\boldsymbol{P}_{i,j}\}$ (neuron correspondences) for $j = 1, \ldots, m$ worms in a training set and $i = 1, \ldots, n$ neurons, $\epsilon$ (convergence tolerance)

**Initalization**

Select random worm $j \sim \text{Unif}[m]$

Set means as neuron centers of worm $j$: $\boldsymbol{\mu}_i^0 \leftarrow \mathbf{y}_{i,j}$ for $i = 1, \ldots, n$

Set covariances as identity: $\boldsymbol{\Sigma}_i^0 \leftarrow \boldsymbol{I}_6$ for $i = 1, \ldots, n$

**while** Not converged **do**

$t \leftarrow t + 1$

**for** j=1,. . . ,n **do**

Solve alignment of $j$th worm to atlas $\{\boldsymbol{\mu}, \boldsymbol{\Sigma}\}$ using equations 4.15 and 4.16

**end for**

Update $\boldsymbol{\mu}^t, \boldsymbol{\Sigma}^t$ using equations 4.13

Check convergence $||\boldsymbol{\mu}^t - \boldsymbol{\mu}^{t-1}||_F \leq \epsilon$ and $||\boldsymbol{\Sigma}^t - \boldsymbol{\Sigma}^{t-1}||_F \leq \epsilon$

**end while**

**return** Statistical atlas of neuron colors and positions $\{\boldsymbol{\mu}^t, \boldsymbol{\Sigma}^t\}$.

---

Algorithm 5: Neuron likelihood computation

**Input:** Neuron detections $\mathbf{y}_k \in \mathbb{R}^6$ for $k = 1, \dots, K$ for a test worm, statistical atlas $\boldsymbol{\mu}, \boldsymbol{\Sigma}$, $\epsilon$ (convergence tolerance).

Initialize $\boldsymbol{\beta}^0 = \boldsymbol{I}_6, \boldsymbol{\beta}_0^0 = \mathbf{0}_6$

**while** Not converged **do**

$t \leftarrow t + 1$

Compute assignment $\boldsymbol{P}^t$ using equation 4.18.

Compute alignment $\boldsymbol{\beta}^t, \boldsymbol{\beta}_0^t$ using equations 4.15, 4.16

Check convergence $\|\boldsymbol{\beta}^t - \boldsymbol{\beta}^{t-1}\|_F \leq \epsilon$

**end while**

**for** i=1,. . . ,n **do**

**for** k=1,. . . ,K **do**

Compute likelihood $\boldsymbol{L}_{i,k}$ of the $k$th neuron from the test worm corresponding to the $i$th atlas neuron using equation 5.4.

**end for**

**end for**

**return** Likelihood matrix $\boldsymbol{L}$

### 5 Step 3b: Probabilistic neuron identification

#### 5.1 Deterministic vs probabilistic estimates of neuronal identity

An ideal automatic neural identification algorithm would always output the true identity. For the current NeuroPAL worms this ideal remains somewhat unrealistic: in some cases even expert users may find it challenging to identify every neuron with full confidence from the 3d color images analyzed here. Therefore it is important to develop methods that quantify the uncertainty in our predictions.

We take the probabilistic generative model from section 4 as our starting point. Suppose a statistical atlas has been trained using the above procedures, i.e., the parameters $(\boldsymbol{\mu}, \boldsymbol{\Sigma})$ have been estimated. Now, given images from a new test worm, we can compute the maximum-likelihood alignment $\boldsymbol{\beta}^*$ and matching $\boldsymbol{P}^*$ by iterating equations 4.16 (alignment) and equation 4.18 (matching), given colors and locations $\boldsymbol{y}_j$ extracted from this test worm. This iterative procedure is outlined in algorithm 5.

The resulting locally best matching $\boldsymbol{P}^*$ will be useful to make deterministic predictions, but is not useful for quantifying uncertainty. Conceptually, what we really want is the posterior distribution over $\boldsymbol{P}$ given the observed data — but the space of permutation matrices $\boldsymbol{P}$ is very large and it is not tractable to work with this posterior distribution directly. Instead, we will approximate *marginals* of this posterior distribution. In particular, we would like to compute the matrix $\boldsymbol{\rho}$ such that $\boldsymbol{\rho}_{k,i}$ is the posterior probability that the $k$-th observed neuron in the test worm corresponds to the $i$-th neuron in the atlas. Unlike deterministic assignments, for a single $k$, $\boldsymbol{\rho}_{k,\cdot}$ may assign mass to *more than one candidate*, and we will use that information to supplement the deterministic estimate. Additionally, $\boldsymbol{\rho}$ admits an interpretation in terms of confidence: rows $\boldsymbol{\rho}_{k,\cdot}$ that are closer to the uniform distribution are the ones for which the model gives low confidence about predictions. Below we will use this information to develop a semi-automated strategy in which the user is prompted to manually annotate these low-confidence neurons; we will show that prediction accuracy increases significantly faster by following this adaptive strategy compared to simple random labeling baselines. However, first we need a method for approximating $\boldsymbol{\rho}$.

#### 5.2 Variational marginal inference with Sinkhorn approximation

The posterior over the matching $P$ for the held out worm expresses as

$$P(\boldsymbol{P}|\boldsymbol{y}, \boldsymbol{\Sigma}, \boldsymbol{\mu}) = \int P(\boldsymbol{P}, \boldsymbol{\beta}|\boldsymbol{y}, \boldsymbol{\Sigma}, \boldsymbol{\mu}) d\boldsymbol{\beta}. \tag{5.1}$$

The observed data provides strong constraints on the affine transformation $\boldsymbol{\beta}$; therefore in the following we replace integration with respect to $\beta$ in the equation 5.1 with a point mass at the MLE, $\beta^*$. Then, by Bayes' rule, and using a uniform prior over $P$, we have that

$$P(\boldsymbol{P}|\boldsymbol{y}, \boldsymbol{\Sigma}, \boldsymbol{\mu}) \propto P(\boldsymbol{y}|\boldsymbol{\mu}, \boldsymbol{\Sigma}, \boldsymbol{\beta}^*, \boldsymbol{P}). \tag{5.2}$$

Now, as in equation 4.6, we can write the right side above as (we exclude the normalizing constant, as it doesn't depend on $\boldsymbol{P}$)

$$\begin{aligned} -\log p(\boldsymbol{y}|\boldsymbol{\mu}, \boldsymbol{\Sigma}, \boldsymbol{P}, \boldsymbol{\beta}^*) &\propto \sum_{k=1}^{K} \frac{1}{2}(\boldsymbol{y}_k - \boldsymbol{\mu}_{\pi_k}\boldsymbol{\beta}^* - \boldsymbol{\beta}^{*0})(\boldsymbol{\beta}^*\boldsymbol{\Sigma}_{\pi_k}\boldsymbol{\beta}^{*T})^{-1}(\boldsymbol{y}_k - \boldsymbol{\mu}_{\pi_k}\boldsymbol{\beta}^* - \boldsymbol{\beta}^{0*})^T) \qquad (5.3) \\ &= \sum_{i=1}^{K}\sum_{k=1}^{K} P_{k,i}\left(\frac{1}{2}(\boldsymbol{y}_k - \boldsymbol{\mu}_i\boldsymbol{\beta}^* - \boldsymbol{\beta}^{*0})(\boldsymbol{\beta}^*\boldsymbol{\Sigma}_i\boldsymbol{\beta}^{*T})^{-1}(\boldsymbol{y}_k - \boldsymbol{\mu}_i\boldsymbol{\beta}^* - \boldsymbol{\beta}^{0*})^T)\right). \end{aligned}$$

The above expresses the fact that each $\boldsymbol{P}$ corresponds to a matching between canonical $\boldsymbol{x}_i = \boldsymbol{z}_i\beta_i^* + \beta^{*0}, \boldsymbol{z}_i \sim \mathcal{N}(\boldsymbol{\mu}_i, \boldsymbol{\Sigma}_i)$ (equations 4.1, 4.2) and observed $\boldsymbol{y} = P\boldsymbol{x}$ neural positions and colors. Therefore, if we define the matrix $\boldsymbol{L}$ of likelihoods of that each $\boldsymbol{y}_k$ comes from the atlas model for neuron $i$

$$\log \boldsymbol{L}_{k,i} = -\frac{1}{2}(\boldsymbol{y}_k - \boldsymbol{\mu}_i\boldsymbol{\beta}^* - \boldsymbol{\beta}^{*0})(\boldsymbol{\beta}^*\boldsymbol{\Sigma}_i\boldsymbol{\beta}^{*T})^{-1}(\boldsymbol{y}_k - \boldsymbol{\mu}_i\boldsymbol{\beta}^* - \boldsymbol{\beta}^{0*})^T), \tag{5.4}$$

we see that all the dependencies of the posterior $P(\boldsymbol{P}|\boldsymbol{y}, \boldsymbol{\Sigma}, \boldsymbol{\mu})$ on data $\boldsymbol{y}$ and inferred parameters $\boldsymbol{\beta}^*, \boldsymbol{\mu}, \boldsymbol{\Sigma}$ are encapsulated through $\boldsymbol{L}$, and moreover that the dependency between $\boldsymbol{P}$ and $\boldsymbol{L}$ is linear. We then write

$$P(\boldsymbol{P}|\boldsymbol{y}, \boldsymbol{\Sigma}, \boldsymbol{\mu}) = \frac{1}{Z_{\boldsymbol{L}}}\exp(\langle \log \boldsymbol{L}, \boldsymbol{P}\rangle_F), \tag{5.5}$$

where $\langle A, B\rangle_F$ is the (Frobenius) matrix inner product and $Z_{\boldsymbol{L}}$ is the normalizing constant

$$Z_{\boldsymbol{L}} = \sum_{\pi \in \mathcal{S}_K} \prod_{i=1}^{K} \boldsymbol{L}_{i,\pi_i}. \tag{5.6}$$

Unfortunately, in general the problem of computing $\boldsymbol{\rho}$ from the posterior 5.5 is intractable because it requires access to $Z_{\boldsymbol{L}}$, also known as the permanent of $\boldsymbol{L}$, and whose computation is known to be a #P-hard problem Valiant (1979) ($Z_{\boldsymbol{L}}$ is defined a summation over the set $\mathcal{S}_K$ of all permutations of $K$, of size $K!$).

To surmount this difficulty we will appeal to the so-called Sinkhorn approximation, which can be computed efficiently and is easy to implement. Specifically, we approximate $\boldsymbol{\rho}$ as $S(\boldsymbol{L})$, where $S(\boldsymbol{L})$ is the Sinkhorn algorithm applied to $\log \boldsymbol{L}$. Briefly, $S(\boldsymbol{L})$ iteratively computes a row and column normalization of $L = exp(\log \boldsymbol{L})$ for $n_s$ iterations. It is known in the limit $n_s \to \infty$, $S(\boldsymbol{L})$ is indeed a doubly stochastic matrix.

We defer to 5.3 for a detailed explanation of this approximation as a type of variational inference method. We also refer the reader to Linderman et al. (2018); Mena et al. (2018) for related uses of the Sinkhorn algorithm in neural identification problems.

Algorithm 6: Sinkhorn algorithm for approximate matching inference.

Input: A square log likelihood matrix $\log \boldsymbol{L}$
Output: A doubly stochastic matrix $S(\boldsymbol{L})$
**for** $l = 0, \ldots, n_s$ **do**
    $\log \boldsymbol{L} \leftarrow \log \boldsymbol{L} - \text{LogSumExp}_r(\log \boldsymbol{L})$ ▷ Row normalization in log space
    $\log \boldsymbol{L} \leftarrow \log \boldsymbol{L} - \text{LogSumExp}_c(\log \boldsymbol{L})$ ▷ Column normalization in log space
**end for**
$S(\boldsymbol{L}) \leftarrow \exp(\log \boldsymbol{L})$

###### 5.2.1 Unequal sizes

In principle, our method is designed only for square matrices, but in real applications we may want probabilistic assignments with unequal sizes, as typically the output of matching pursuit may contain false positives (non-neural identified objects) or false negatives (e.g., dim neurons that may be challenging to detect). We found it is possible to account for such situations by simply expanding the original matrix to enforce it to be square, and filling the new entries $\log L_{k,i}$ with a very negative number. In this way we avoid violating the equal size constraint on permutation matrices. To see this, call $\tilde{L}$ the expanded matrix and denote $\tilde{k}, \tilde{i}$ the novel row and column indexes. If $\log L_{i,j} = -\infty$, from equation 5.5 we see all assignments $P_{k,\tilde{i}}$, $P_{\tilde{k},i}$, $P_{\tilde{k},\tilde{i}}$ have zero probability, and therefore all the mass in the assignments is distributed as if these new indices didn't exist.

##### 5.3 Details on the Sinkhorn approximation

Here we explain why $S(\boldsymbol{L})$ is a sensible approximation to $\boldsymbol{\rho} = \boldsymbol{\rho}^{\boldsymbol{L}}$. First, by elementary convex duality in exponential families, we see that marginal inference of $\boldsymbol{\rho}^{\boldsymbol{L}}$ is intimately related to the computation of $Z_{\boldsymbol{L}}$.

Indeed, notice that equation (5.5) defines an exponential family over permutations with sufficient statistic $\boldsymbol{P}$ and parameter $\log \boldsymbol{L}$, and recall Theorem 3.4 of Wainwright et al. (2008):

$$\log Z_{\boldsymbol{L}} = \sup_{\mu \in \mathcal{M}} \langle \log \boldsymbol{L}, \mu \rangle - A^*(\mu), \tag{5.7}$$

where $\mathcal{M}$ is the marginal polytope (here, the Birkhoff polytope, the set of doubly stochastic matrices) and $A^*(\mu)$ is the dual function $\log Z_{\boldsymbol{L}}$, i.e.

$$A^*(\mu) = \sup_{\boldsymbol{L}} \langle \log \boldsymbol{L}, \mu \rangle - \log Z_{\boldsymbol{L}}. \tag{5.8}$$

Moreover, for a given $\boldsymbol{L}$, $\mu(\boldsymbol{L})$ achieving the supremum in (5.7) is exactly the matrix of marginals, $\mu(\boldsymbol{L}) = \rho^{\boldsymbol{L}}$ and the dual function $A^*(\mu(\boldsymbol{L}))$ coincides with the negative entropy of the distribution in equation (5.5). From this we see marginal inference of $\rho^{\boldsymbol{L}}$ and computation of the permanent $Z_{\boldsymbol{L}} = perm(\boldsymbol{L})$ are linked by the optimization problem in (5.8).

Now, we appeal to the variational inference framework (Wainwright et al., 2008) to justify the approximation $S(\boldsymbol{L}) \approx \boldsymbol{\rho}^{\boldsymbol{L}}$. In variational inference, the intractability of $Z_{\boldsymbol{L}}$ is handled through approximations to either the entropy function, or the optimization set in equations 5.7, 5.8. These approximations lead to approximate normalizing constants, and the quality of the variational routine will typically relate to the quality of the approximation of $Z_{\mathbf{L}}$.

The Sinkhorn approximation is a type of variational inference. This comes from the following well-known variational representation of $S(\boldsymbol{L})$ (Mena et al., 2018; Helmbold and Warmuth, 2009):

$$S(\boldsymbol{L}) = \arg\sup_{\mu\in\mathcal{M}} \langle \log \boldsymbol{L}, \mu \rangle - \sum_{i,j} \mu_{i,j} \log \mu_{i,j}. \tag{5.9}$$

Thus, the approximation is justified by considering the entry-wise entropy $-\sum_{i,j} \mu_{i,j} \log \mu_{i,j}$ as a sensible approximation of the entropy of (5.5). Although this is ultimately a heuristic, it is known there is some control in the corresponding approximations to the permanent (Linial et al., 2000). Explicit bounds are stated in the following proposition:

**Proposition 1.** *Define the Sinkhorn approximation of the (log) permanent as the value that* $S(L)$ *achieves in equation* (5.9)*. Then,*

$$perm(\boldsymbol{L}) \le perm_S(\boldsymbol{L}) \le e^n perm(\boldsymbol{L}). \tag{5.10}$$

*Proof.* We use the fact that the permanent of a doubly stochastic matrix $B$ of size $n$ satisfies (Linial et al., 2000)

$$e^{-n} \le perm(B) \le 1.$$

Also, it can be verified that $S(\boldsymbol{L}) = diag(x)\boldsymbol{L}diag(y)$, where $diag(x), diag(y)$ are positive vectors turned into diagonal matrices (Peyré et al., 2019). Then,

$$perm(S(\boldsymbol{L})) = \left(\prod_{i=1}^{n} x_i\right)\left(\prod_{i=1}^{n} y_i\right) perm(\boldsymbol{L}).$$

Additionally, we obtain the (log) Sinkhorn approximation of the permanent of $\boldsymbol{L}$, $perm_S(L)$, by evaluating $S(\boldsymbol{L})$ in the problem it solves, (5.9). By simple algebra and using the fact that $S(\boldsymbol{L})$ is a doubly stochastic matrix we see that

$$\log perm_S(\boldsymbol{L}) = -\sum_{i=1}^{n} \log(x_i) - \sum_{j=1}^{n} \log(y_i).$$

By combining the last three displays we obtain

$$e^{-n} \le perm(L)/perm_S(\boldsymbol{L}) \le 1, \tag{5.11}$$

from which it follows that

$$perm(\boldsymbol{L}) \le perm_S(\boldsymbol{L}) \le e^n perm(\boldsymbol{L}).$$

□

We conclude this section by discussing the relation with the better known Bethe approximation (Huang and Jebara, 2007; Chertkov et al., 2010; Vontobel, 2014; Tang et al., 2015). In this method the dual function $A^*(\mu)$ is approximated by the value it would take if the underlying Markov Random Field had a tree structure (Yedidia et al., 2001). The corresponding approximation $B(L)$ of the marginals can be computed through belief propagation (Huang and Jebara, 2007; Vontobel, 2013), and it enjoys better theoretical guarantees than the Sinkhorn approximation. Indeed, the Bethe approximation of the permanent, $perm_B(\cdot)$ is a tighter approximation to the permanent, with the following known bounds (Gurvits and Samorodnitsky, 2014; Anari and Rezaei, 2018)

$$perm(\boldsymbol{L}) \le perm_B(\boldsymbol{L}) \le \sqrt{2}^n perm(\boldsymbol{L}). \tag{5.12}$$

However, we observed a much better empirical performance of the Sinkhorn approximation. This is explained by two reasons, summarized in figure 5.1. First, the Bethe approximation arises as the solution of a non-convex problem (Vontobel, 2014), but in practice local minima from random initial guesses lead to worse solutions than $S(\boldsymbol{L})$, defined through a convex program. Second, typically there are huge entry-wise fluctuations in $\boldsymbol{L}$, and in such situations the Sinkhorn approximation is much tighter than its the theoretical upper bound.

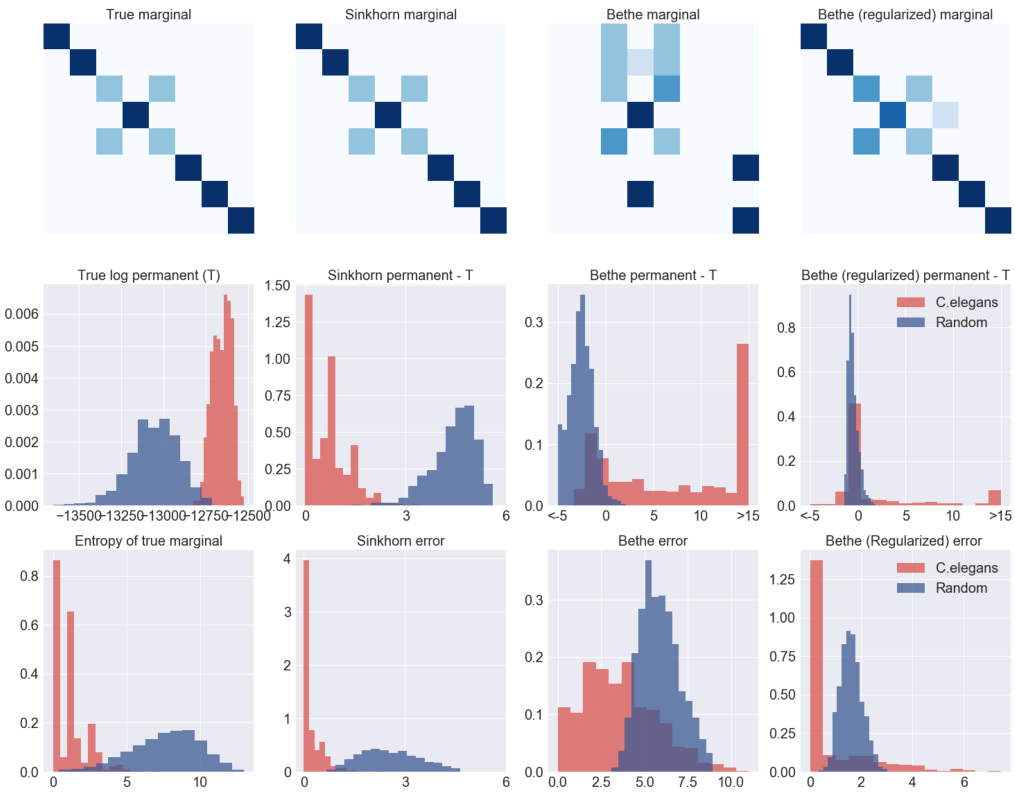

Figure 5.1: Empirical assessment of Sinkhorn and Bethe approximations on real and simulated data. Top row: example of true marginal from the model Sinkhorn, Bethe, and regularized Bethe approximations. The regularized Bethe (Huang and Jebara, 2007) uses a penalty term $\epsilon = 0.1$ to encourage smoother optimization in the log space. Second row: first plot shows histogram of the true permanents in real and simulated data. The remaining plots show differences between approximations and true values. Notice the values for the Bethe approximation are inconsistent with theoretical bounds, a consequence of convergence to local but not global optima. Third row: first plot shows the histogram of entropy $-\sum_{i,j} P_{i,j} \log P_{i,j}$ of true marginal, and the remaining plots are histograms of the the $l_1$ approximation error of different methods. Real data were obtained by sampling 8x8 random sub-matrices from 14 log likelihood matrices, corresponding to 14 worms. Simulated data were obtained with entries simulated independently from $\mathcal{N}(\mu, \sigma^2)$ with $\mu$ equal to the mean log likelihood on real matrices and $\sigma^2 = 5$.

### 6 Results / numerical validations

#### 6.1 Further details of statistical atlases

In this section we provide further details on the statistical atlas presented in Figure 3 in the main text. In particular, Fig. 6.1 illustrates the variability of the positions of neurons in individual worms, while Fig. 6.2 illustrates the variability when extracting neuron colors (real-world microscopy images suffer from anisotropy, camera sensor saturation, dense neuron groups whose colors can commingle, and other such confounds).

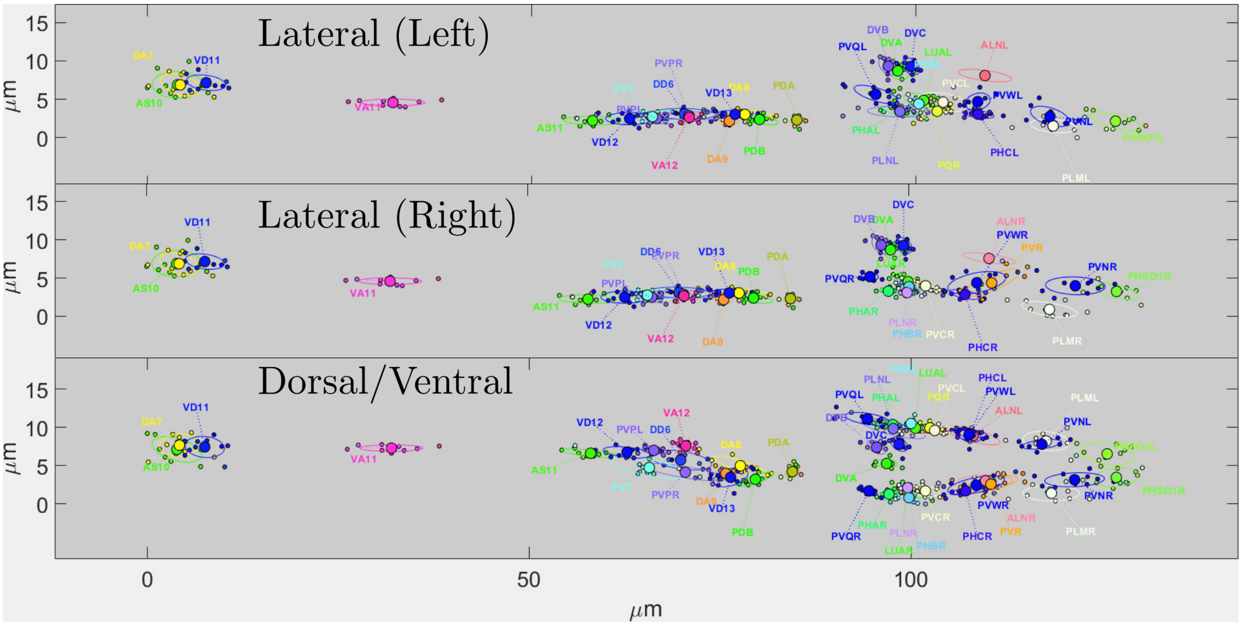

Figure 6.1: The statistical atlas of positions of *C. elegans* neurons of the tail. The centers denote the mean positions while the ellipses denote the contours that delineate the half quantile. The smaller points denote the aligned neurons of individual worms that were used to train the statistical atlas. Note that this is 2D projection of a 3D atlas, therefore the x and y axes denote the major and minor axes of position in pixels. Top: Lateral projection with only left sided neurons. Middle: Lateral projection with only right sided neurons. Bottom: Dorsal/ventral projection with all neurons. For worm-wise alignment to the statistical atlases, see the videos for head and tail. Note: Lateral ganglia are removed for dorsal/ventral view in the head for clarity purposes. Likewise, ventral ganglion is removed from the lateral view.

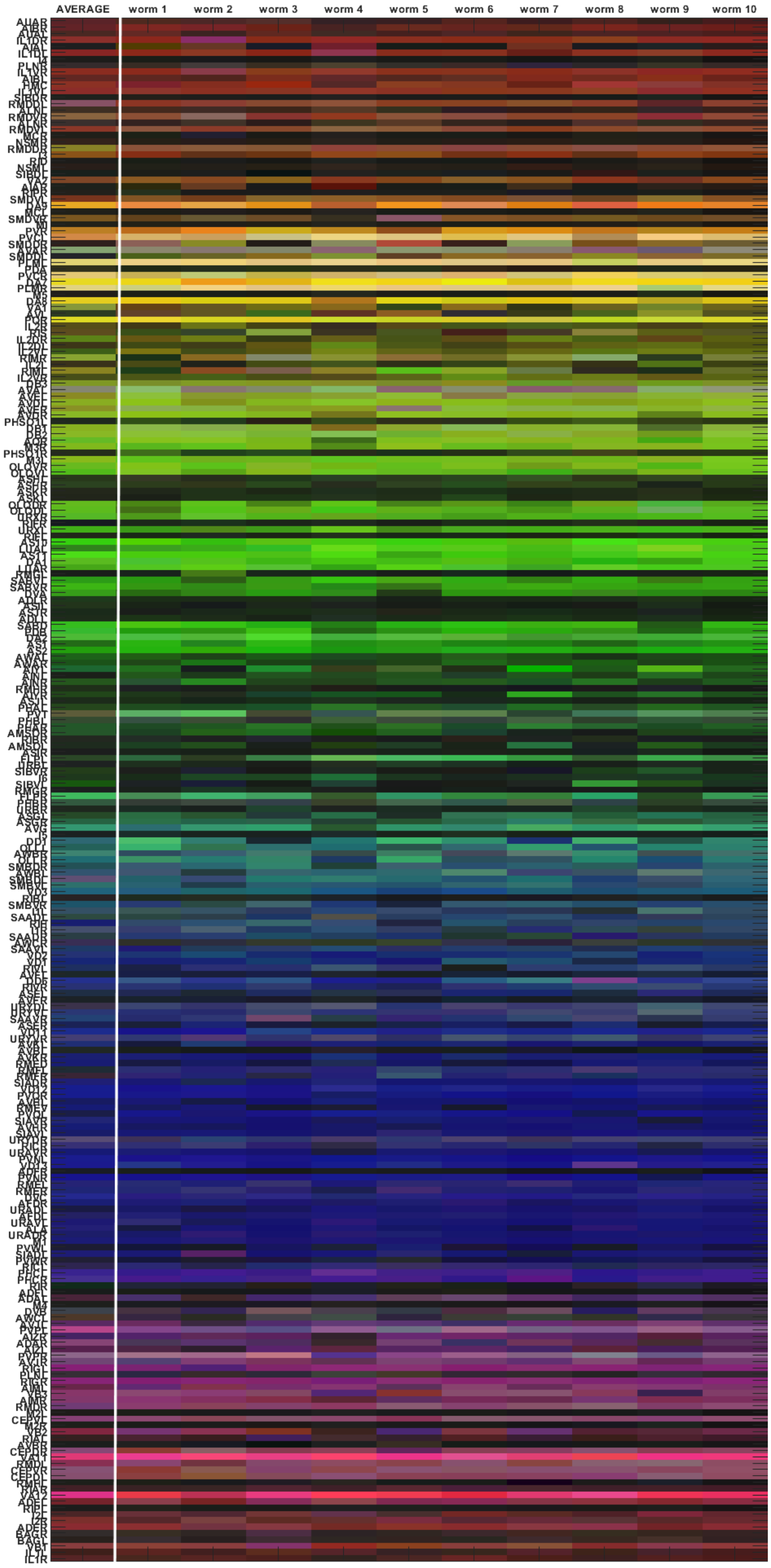

Figure 6.2: The statistical atlas of colors of NeuroPAL neurons. The rows indicate neurons. The first column indicates the mean color estimated in the statistical atlas. The subsequent columns denote the colors of the neurons of training worms aligned to the statistical atlas to demonstrate the extent of variability present upon color extraction (we extract the colors by computing the median of the pixel intensities in a $1.5 \times 1.5 \times 0.75$ $\mu m$ patch centered at the location of each neuron in the z-scored image). Extracted color variability is due to image anisotropy, camera sensor saturation, dense neuron groupings, and a variety of other confounds. Note that black colored elements may indicate neurons that are not found or are pan-neuronal colored.

#### 6.2 Quantifying the performance of the detection step

We used the common evaluation metrics to quantify the performance of our detection algorithm.

$$\text{Accuracy} = \frac{\text{TP} + \text{TN}}{\text{TP} + \text{TN} + \text{FP} + \text{FN}}; \quad \text{Precision} = \frac{\text{TP}}{\text{TP} + \text{FP}}; \quad \text{Recall} = \frac{\text{TP}}{\text{TP} + \text{FN}};$$

In this context TP corresponds to correctly identified neurons up to a small pre-defined error (3 $\mu m$, approximating adult hermaphrodite neuronal nuclear diameter). FP is the number of incorrectly identified neurons that consists of multiple counting of the same cell or the wrong detection of non-neuronal objects. FN counts the number of incorrectly rejected neurons, i.e. the neurons that are identified by a human expert but the algorithm is unable to recover them. TN is always zero because there are no negative samples that the algorithm needs to reject.

To avoid double counting in calculating TP and FP we need to uniquely assign each location that is recovered by the algorithm to a neuron that is hand annotated by human expert, and lies within a distance of 3 $\mu m$ from that location. For that we compute the pairwise distance matrix between hand annotated neurons and recovered locations and set the values that are larger than 3 $\mu m$ to infinity. We then run the Hungarian algorithm on the resulting distance matrix to find the unique assignments.

As shown in figures 6.3 our algorithm reaches the accuracy of 80% in head and 88% in tail. This occurs when we find $K = 191$ and $K = 44$ neurons in head and tail in which case we get similar precision and recall values that are both equal to 90% and 93% in head and tail respectively.

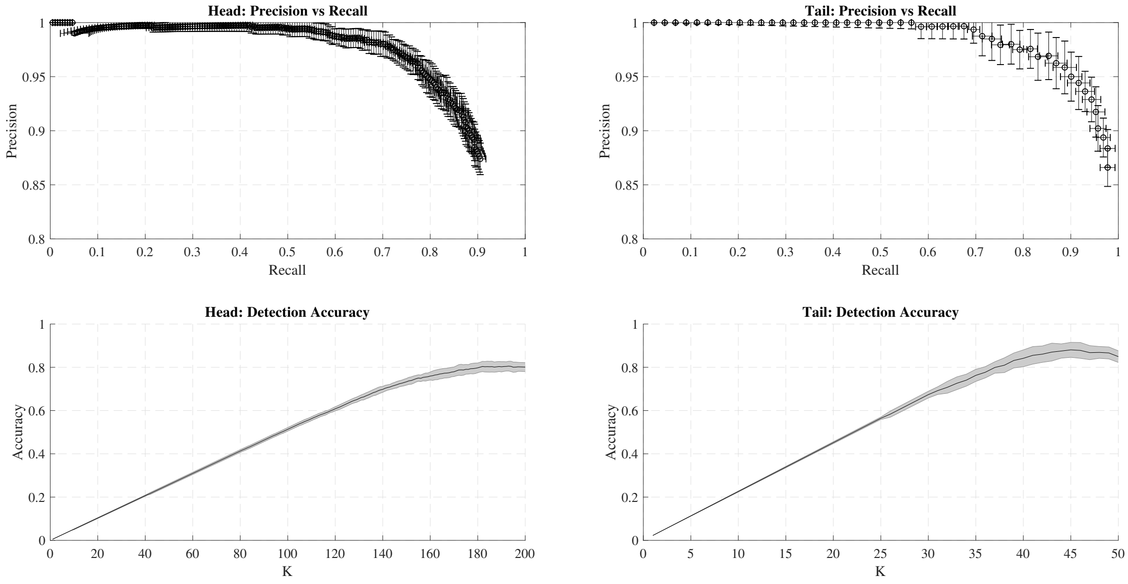

Figure 6.3: Evaluation of the detection performance for 10 worms. Error bars and shades correspond to one standard deviation away from the mean.

#### 6.3 Identification error / uncertainty analysis

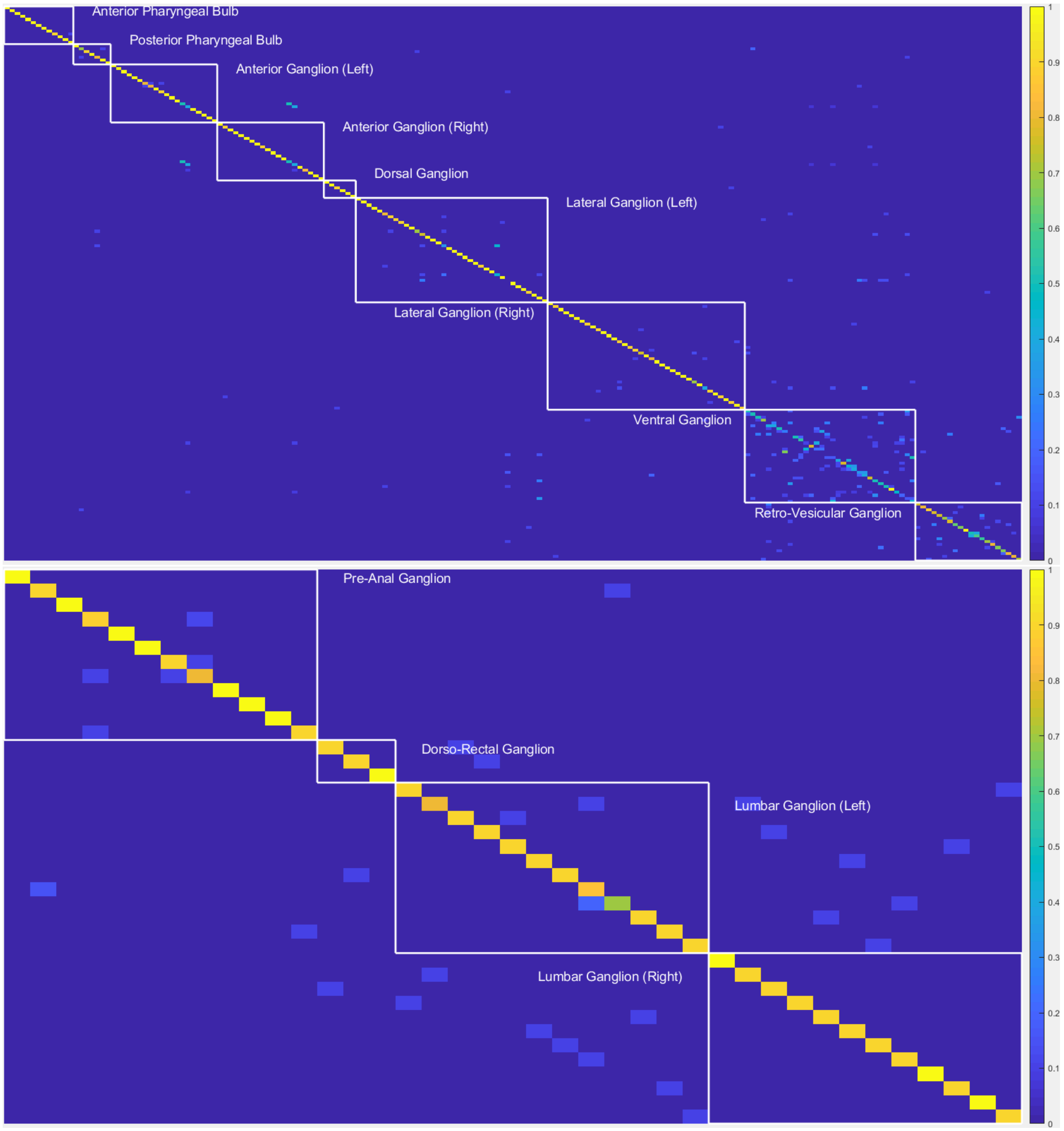

Figure 6.4: Confusion matrices of head (top) and tail (bottom) neuron identification using the full proposed model. The matrix is organized by ganglia. The rows indicate out of sample neurons and the columns indicate training neurons in the same order as the testing neurons. The cells of this matrix record the rate of assignment of the test neurons to training neurons where the colors indicate assignment rate scaled by the color-bar; ideally a fully diagonal matrix indicates perfect identification accuracy. The confusion matrix was computed using leave-one-out cross validation procedure on 10 worms.

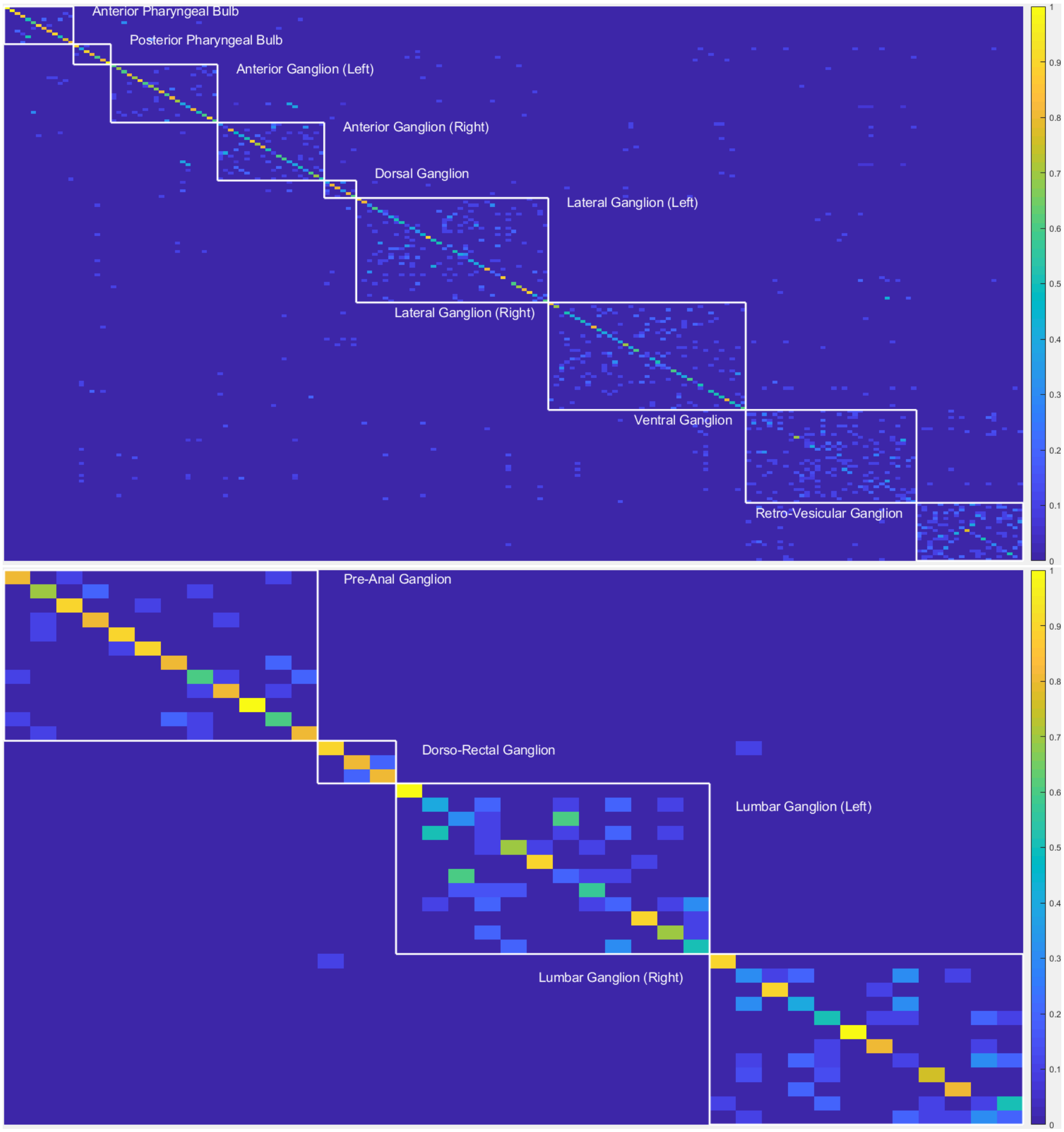

Figure 6.5: Confusion matrices of head (top) and tail (bottom) neuron identification using **only position** information. Conventions as in previous figure.

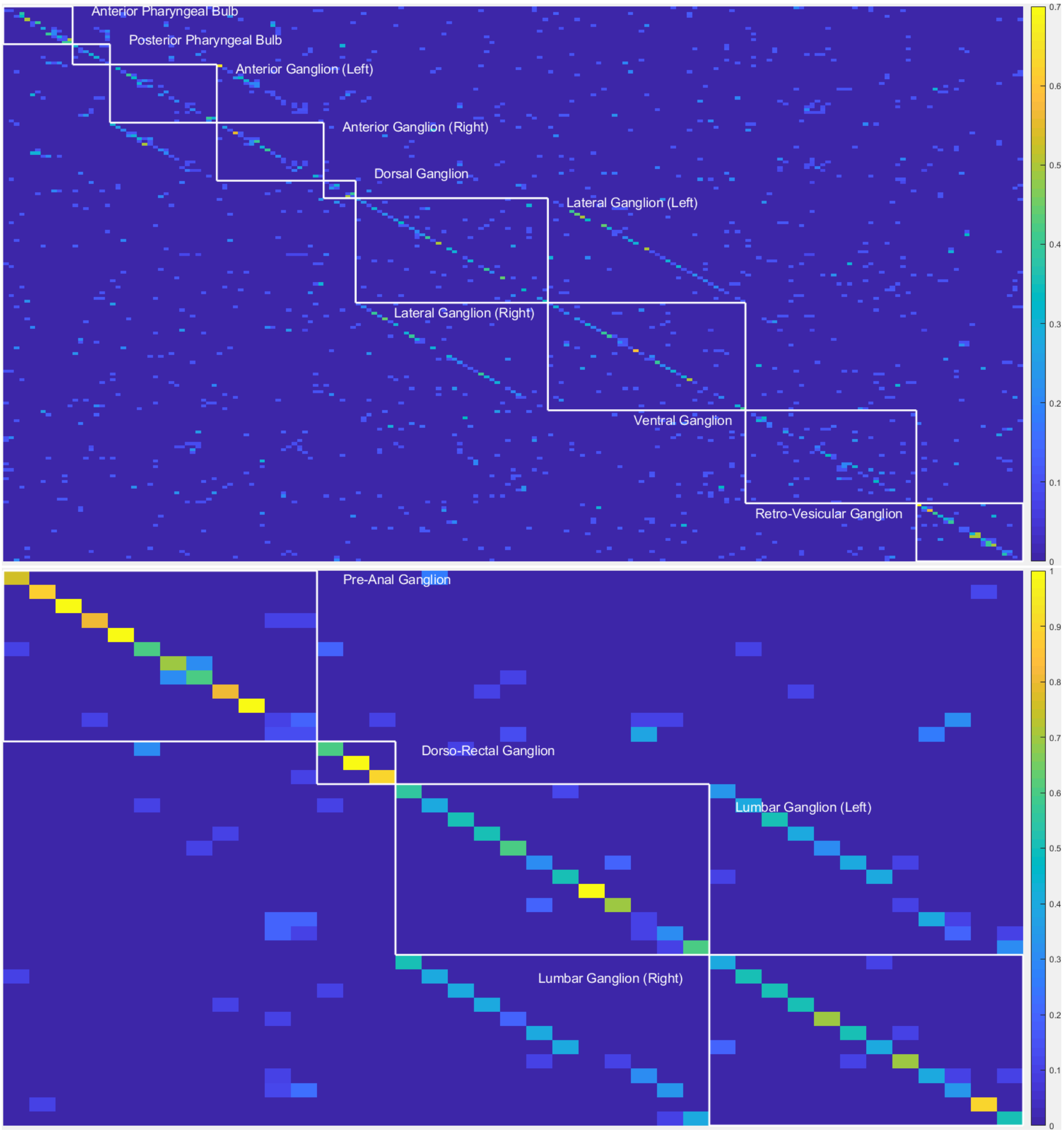

Figure 6.6: Confusion matrices of head (top) and tail (bottom) neuron identification using **only color** information. Conventions as in previous figure. Note the left/right symmetry visible in many of the errors.

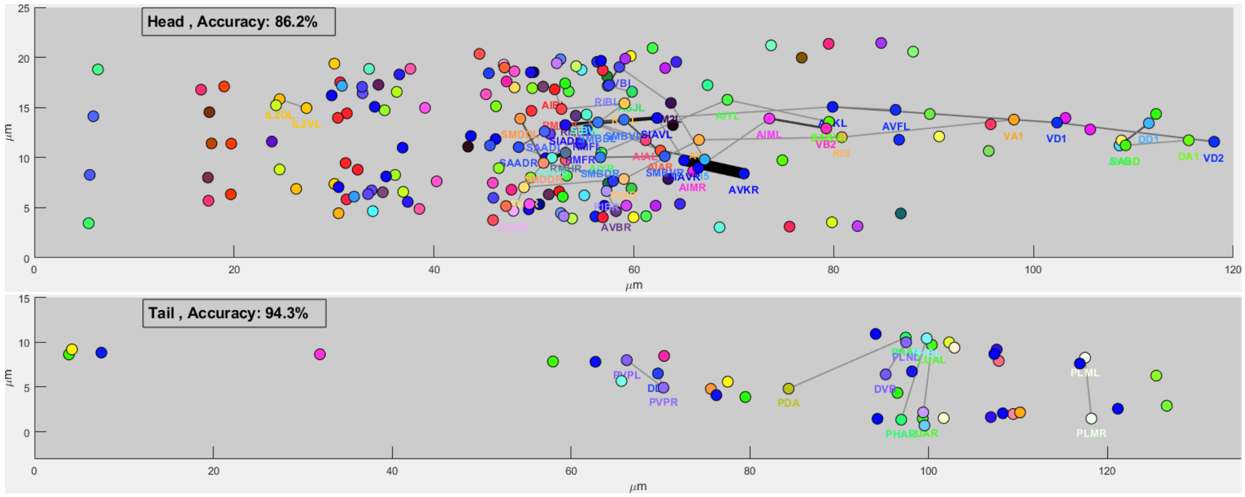

Figure 6.7: The neuron misidentification pairs are illustrated as edges in a graph. The thickness of the edges relative to marker sizes denotes the confusion rates i.e. lines as thick as markers indicate 100% confusion while lines 20% as thick as markers indicate 20% confusion. Only errors exceeding 10% are shown. The reduction of confusion as a function of user neuron labeling can be seen in the movies for head and tail.

#### 6.4 Calibration

Figure 6.8 addresses the issue of *calibration* of the probabilistic output of our algorithm (Guo et al., 2017): does the error rate scale directly with the algorithm's confidence in each identification? Figure 6.8 shows that confidence and accuracy [1] are largely in good agreement, as desired.

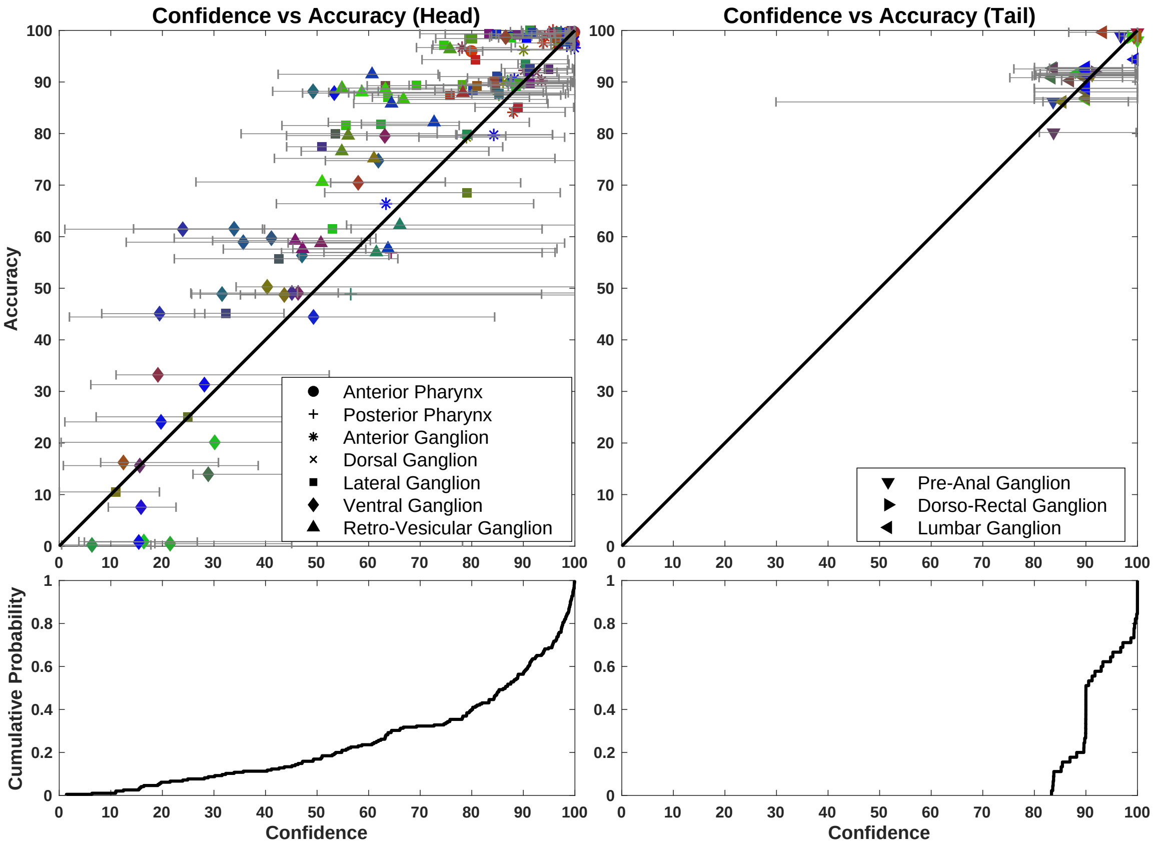

Figure 6.8: Calibration. Top row: reliability scatterplots, i.e. accuracy vs confidence for each individual neuron on tails (left) and heads (right), computed as averages over the worms with ground truth ($n$ = 10). Horizontal lines display 25% and 75% percentiles for confidences. Marker color indicates mean neural color, marker symbol indicates ganglia. Both accuracy and confidence . Bottom row: corresponding confidence CDFs. Bottom row: confidence cdf from the same averages in the scatterplot.

#### 6.5 Guiding annotation by most uncertain neurons

The graphical user interface successively recommends which neurons to annotate manually. Once a neuron is annotated, the reduction in uncertainty and error can be propagated to the remaining neurons by a suitable update on the entries of $\boldsymbol{L}$ and $\boldsymbol{P}$. We found an efficient strategy is to recommend the neurons for which the model expresses the highest uncertainty, i.e. the ones for which the distribution $\rho_{i,\cdot}$ is closest to the uniform distribution. Figure 6.9 shows that following this strategy, the accuracy increases much faster

[1] We used neuron-class to determine accuracy in those cases where left/right neuron pairs are present in the same ganglion, near the midline, such that their left/right sidedness cannot be distinguished (e.g., we used the 'RMF' class in place of 'RMFL' and 'RMFR').

than by a random recommendation. We illustrate the evolution of $\rho$ as a function of number of human labels (in a single example worm) in two videos: head and tail.

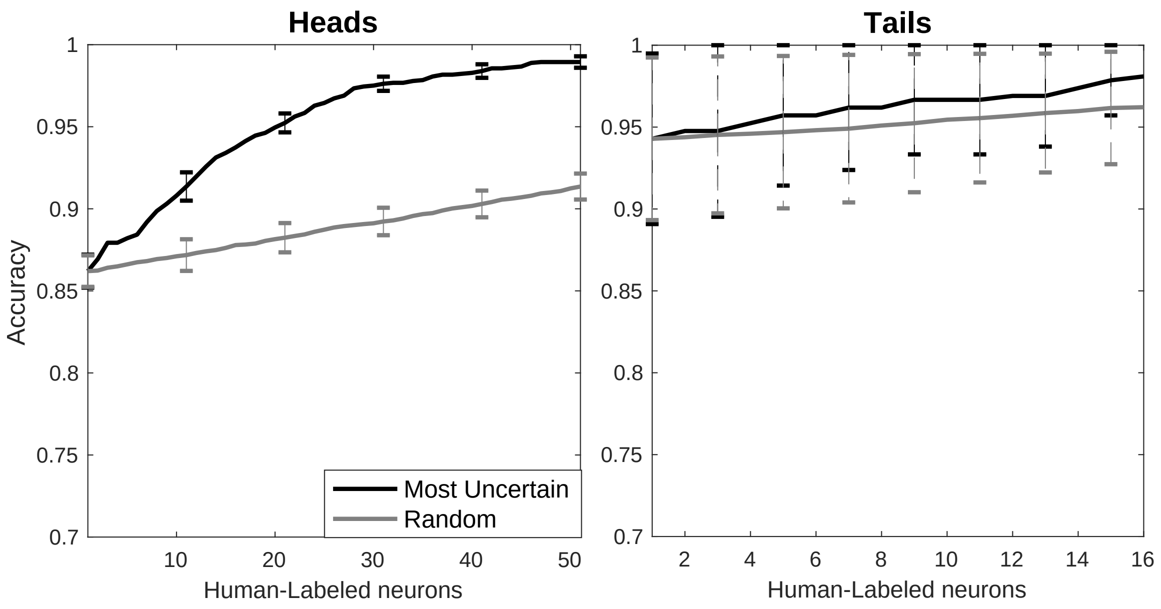

Figure 6.9: Accuracy as a function of number of human labels, for both random and most uncertain recommendations. Note that labeling the most uncertain neurons first leads to faster accuracy improvements.

#### 6.6 Semi-supervised approach for incorporating information from unlabeled worms

It may be easier to image many worms than to fully manually label all of the worms. Can we take advantage of information in the unlabeled worms to improve the accuracy of our estimated statistical atlas? It is natural to extend our expectation maximization algorithm to handle both labeled (supervised) and unlabeled (unsupervised) worms; this is an instance of "semi-supervised learning." In the E step we probabilistically impute the identities of the unlabeled neurons, as in algorithms 5 and 6 for assigning identity on test worms; the M step is the standard M step for updating parameters in the mixture of Gaussians model. Preliminary results are encouraging; full details will be provided in a future draft.

### 7 Graphical user interface

We have incorporated the above procedures into a GUI[2] that displays the raw images overlaid by the locations of the detected nuclei along with the corresponding $P_{ij}$ values of the most likely matches. The user can choose to examine the nuclei for which the corresponding row of $P_{ij}$ has maximal uncertainty (i.e., the nuclei for which we are least sure of the neuronal label). After the user updates the label for a neuron, the GUI quickly updates the label probabilities $P$ (and uncertainty order) for the remaining cells. This closed-loop approach can significantly reduce the overall labor required by the user to label the full population with a high degree of confidence.

#### 7.1 Object oriented design

To build a fast and scalable system we followed object oriented design practices. We identified internally coherent and externally independent classes in our environment and established their properties and behavior, as well as their interactions with other modules. We have separate packages for data handling, methods, logging, biological objects, and user interface. Each of these packages contains relevant classes and the functionalities of the system is built upon the interactions between the classes. For reusability purposes we tried to develop a modular system with independent modules. This allows the users of the system to reuse different compartments of the system for other purposes. An illustration of the packaging and class relations is presented in figs. 7.1a and 7.1b.

#### 7.2 Functionalities

We designed this system to allow researchers to identify cells quickly and accurately. More specifically the system meets the following requirements:

- Automatically detect cell locations and colors in NeuroPAL images.
- Add, remove, or modify the detected cell locations manually.
- Automatically identify and label cells in NeuroPAL images.
- Annotate the cells manually and modify the cell identities found by automatic labeling.
- Load and save the result of labeling and detection.
- Apply basic image processing filters to improve visualization. The implemented methods include gamma correction, histogram matching, decimation (downsampling), and fixed and adaptive thresholding.

A schematic of the use cases of the system and how each use case is performed through the graphical user interface is shown in figs. 7.1c and 7.2.

[2]https://github.com/amin-nejat/CELL_ID

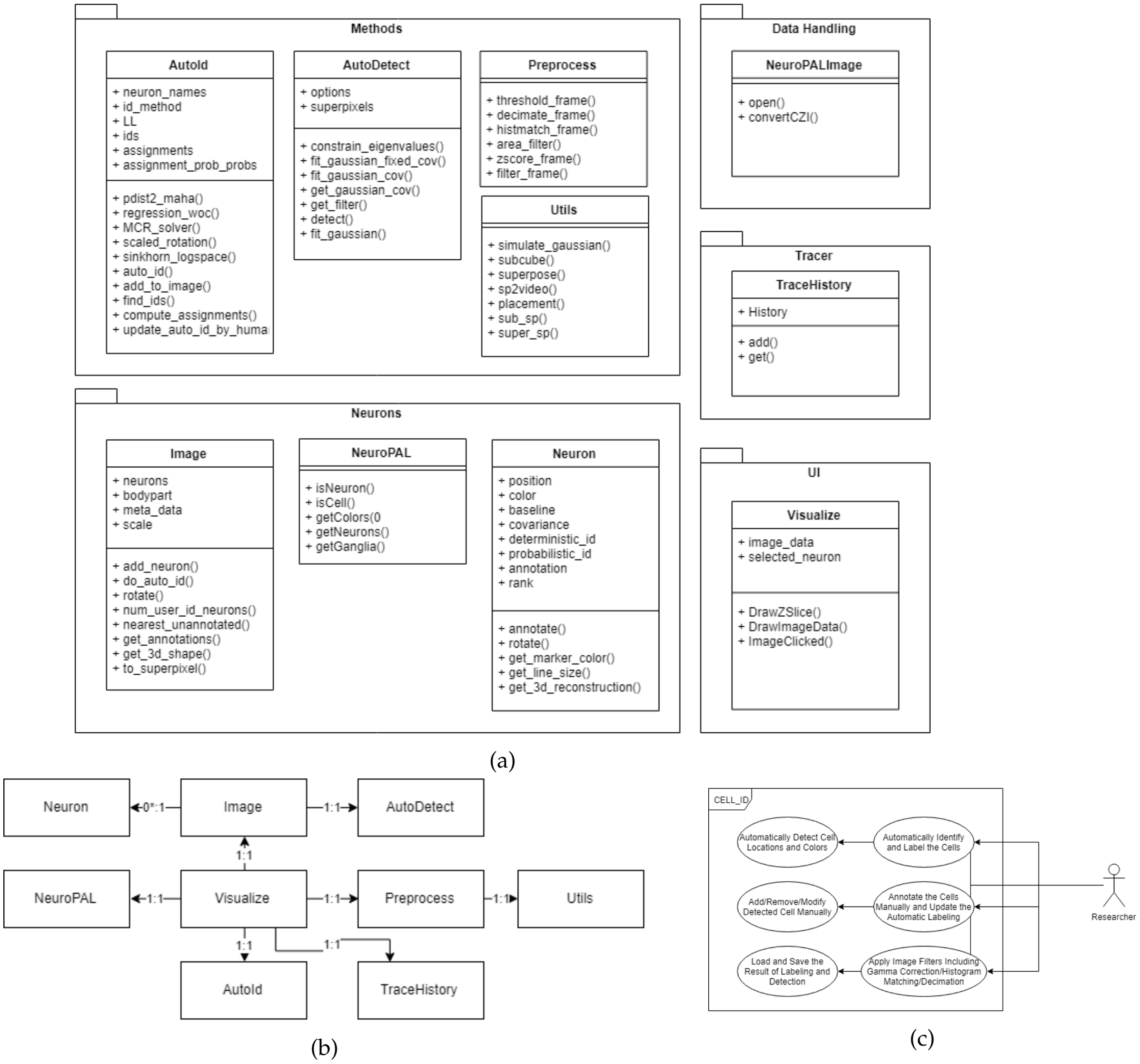

Figure 7.1: Object oriented design plots showing the packaging, class relations, and the use cases of the system. (a) packaging diagram, (b) class relations diagram, (c) use case diagram.

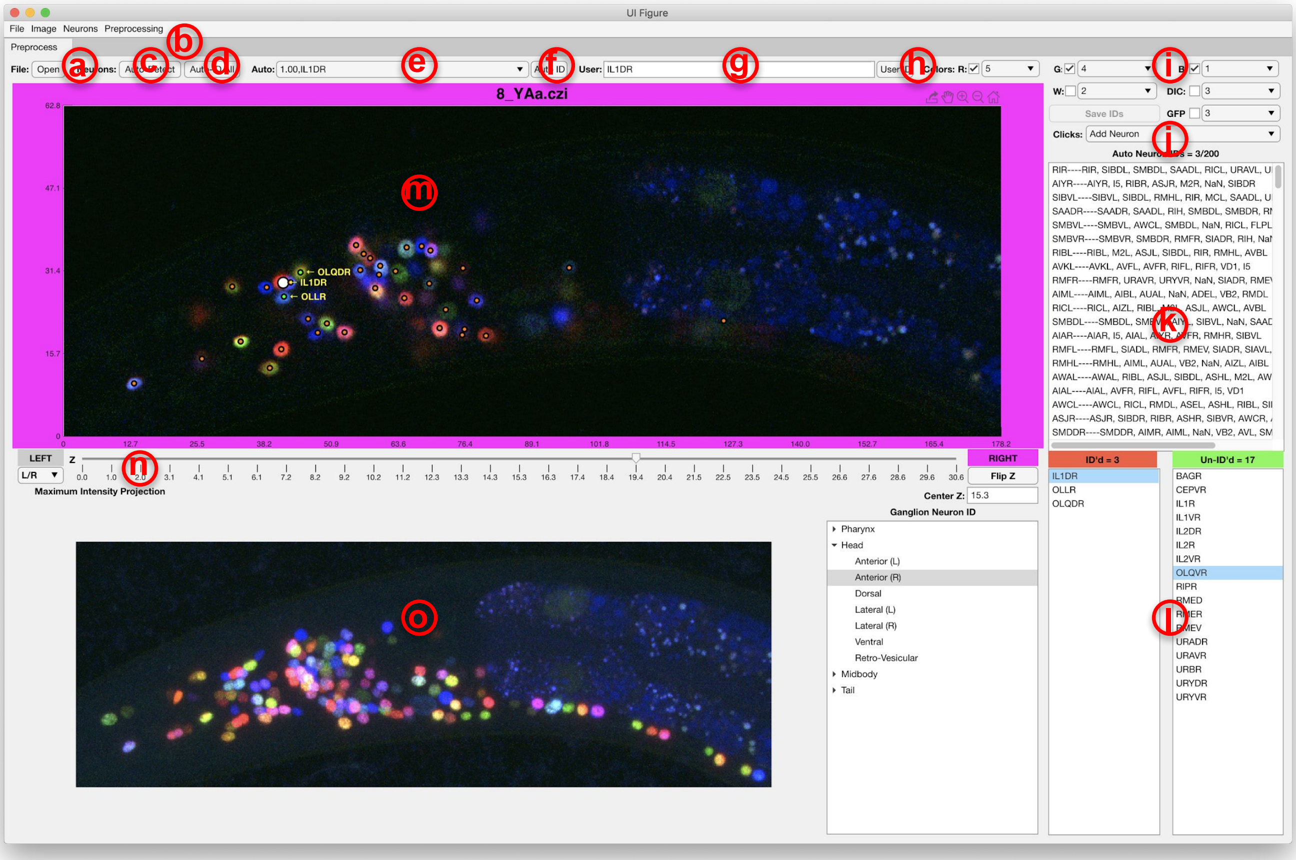

Figure 7.2: GUI environment. (a) open and load a NeuroPAL image, (b) menu for basic preprocessing and improving the visualization, (c) automatically detect the neurons in NeuroPAL images, (d) automatically identify and label the detected neurons (e) clicking on each neurons shows the potential names that our method has found for that neuron and their associated uncertainties, (f) confirm the neuron identity, (g,h) manually annotate neurons, (i) selecting color channels used for visualizing NeuroPAL image, (j) select double click action: manually add a neuron/automatically find a neuron near the clicked location, (k) list of neuron names sorted by their uncertainty according to the result of our method, (l) true neuron names and colors categorized by their corresponding ganglia, (m) z-stacks of NeuroPAL image and text labels for annotated or confirmed neurons, (n) controller for moving through different z-stacks, (o) maximum intensity projection of the image.

#### Acknowledgements

Thanks to S. Linderman and A. Leifer for many helpful discussions. We gratefully acknowledge funding from NIBIB R01 EB22913, NSF NeuroNex Award DBI-1707398, the Simons Collaboration on the Global Brain, and the Gatsby Charitable Foundation.
