## Supplemental Text S2 for "NeuroPAL: A Neuronal Polychromatic Atlas of Landmarks for Whole-Brain Imaging in *C. elegans*"

### Supplemental Text S2. Optimal-Coloring Software

### Optimal cell coloring

#### 1 Introduction

The NeuroPAL transgene contains a combination of 41 selectively overlapping neuron-specific reporters, each of which expresses a subset of four distinguishably-colored fluorophores. This combination of reporters and colors generates a comprehensive color-coded atlas for the entire hermaphrodite nervous system. In NeuroPAL, each neuron expresses a stereotyped combination of fluorophores to yield an invariant color map across individuals, where every neuron is uniquely identified by its color and position.

We built NeuroPAL empirically, laboriously testing a large variety of reporter-fluorophore combinations. To facilitate the generation of further analogous multicolor landmarking solutions, for any collection of cells in any organism, we developed an optimal-coloring algorithm. Our algorithm computes approximately optimal solutions in order to offer multiple reporter-fluorophore combinations to test in vivo (`Optimal-Coloring Software: https://www.hobertlab.org/neuropal/`). With this software, the amount of empirical testing required to construct future multicolor landmarking reagents is expected to be significantly reduced.

As an overview of our algorithm, in order to be individually identifiable, neighboring cells of different types must be distinguishable from each other. Cells that cannot be distinguished by morphology must be distinguished by color and/or intensity differences larger than a *discrimination margin*. In our software, the user chooses this margin, decides which cells must be distinguishable from each other, specifies the number of landmark fluorophores to use in combination with a list of available reporters that have known expression, and restricts the total number of reporter-fluorophore combinations permitted by their transgenesis techniques. Given these inputs, the algorithm generates solutions (reporter-fluorophore combinations) that approximately minimize the sum of color margin violations; this sum serves as a surrogate for the absolute number of color margin violations.

We show that our algorithm generates multiple, promising approximately-optimal-coloring solutions to test *in vivo*. We find that, while NeuroPAL significantly outperforms random reporter-fluorophore combinations, a more optimal multicolor solution may still exist to drive even greater visual distinguishability among neighboring neurons. Furthermore, our results indicate that even when real-world limitations severely restrict the number of distinguishable colors, discriminability between color intensities, and/or the number of reporters that can be used, our algorithm still generates solutions that benefit cell identification. These results show the power of our approximately-optimal-coloring technique in aiding the design of real-world multicolor landmarking solutions to identify collections of cells in any model organism.

#### 2 Maximum margin graph coloring

Here we describe a general-purpose computational technique for combining cell-specific reporters which drive a small set of fluorescent-colored proteins towards obtaining an optimal color map to distinguish cell types (e.g., similar to the reporter-fluorophore combinations used to color NeuroPAL). The idea is that neighboring cells exhibit distinguishable, identifiable coloring (i.e., by co-opting a small subset of the cell's transcriptional profile in order to also express a distinguishable profile of colored proteins). The overall concept of selecting an optimal set of colors to distinguish neighboring cells (by their type) falls under a subfield of graph theory termed fractional graph coloring [Scheinerman and Ullman, 2011]. Here the objective is to color the vertices of a given graph using color mixtures such that no two adjacent vertices share a common mixture component. We relax this objective further by allowing neighboring cells to share a color component (i.e., a cell can be red and green, while its neighbor is red and blue) as long as a minimal color margin between all neighboring cells is maintained. We use the terminology approximately-optimal to allude to the fact that the optimization problem of obtaining optimal colors is NP-hard to solve exactly, and our relaxed formulation gives us an approximate solution.

Let $\boldsymbol{R} \in \{0,1\}^{n \times r}$ denote the expression of $r$ cell-specific reporters in each of the $n$ cells. As an example of $\boldsymbol{R}$, *C. elegans* has a published "Brain Atlas" of nearly 1,200 gene reporters ($r$) accompanied by their precise neuronal expression ($n$) [Hobert et al., 2016], curated from WormBase, a community-wide database of worm reporters. Other model organisms, e.g., fly, fish, and mouse, have analogous databases: FlyBase, ZFIN, and MGI [Eppig, 2017, Thurmond et al., 2019, Ruzicka et al., 2019] respectively. Let $\boldsymbol{X} \in \{0,1\}^{r \times c}$ denote the optimization variable for fluorescent-color assignment of the $r$ reporters, using $c$ different colors (e.g., mTagBFP2, CyOFP1, and mNeptune2.5 as were used in NeuroPAL). As an example, if $c = 3$ for $red$, $green$, and $blue$, then $\boldsymbol{X}_j = [1\ \ 0\ \ 0]$ denotes a red coloring of the $j$th reporter. Furthermore, let $\boldsymbol{A} \in \{0,1\}^{n \times n}$ denote the adjacency matrix of the $n$ cells such that $\boldsymbol{A}_{i,j} = 1$ if cells $i$ and $j$ are within each other's spatial covariance ellipses that capture $95\%$ of probability mass (e.g., the statistical atlas of worm neuron positions and their variability, measured using NeuroPAL – see **Text S1**). If a map of cell positions is unavailable, this adjacency matrix $\boldsymbol{A}$ can further be relaxed to accommodate any arbitrary user-defined metric, as long as it encodes cells that are sufficiently close to each other so as to require color discrimination.

We then optimize the following margin maximization problem cast as a mixed integer quadratic program:

$$
\begin{aligned}
\underset{\boldsymbol{X},m}{\text{minimize}} \; & \sum_{i,j} m_{i,j} \\
\text{s.t. } & \|\boldsymbol{R}_i\boldsymbol{X} - \boldsymbol{R}_j\boldsymbol{X}\|_F^2 \geq \gamma \boldsymbol{A}_{i,j} - m_{i,j} \\
& \boldsymbol{X} \in \{0,1\}^{r\times c} \\
& \boldsymbol{R}\boldsymbol{X}\boldsymbol{1} = \boldsymbol{1} \\
& \boldsymbol{X}^T\boldsymbol{1} \geq \boldsymbol{1} \\
& m_{i,j} \geq 0
\end{aligned}
\tag{1}
$$

The goal of this optimization program is to ensure that all nearby cells have significant color differences. Here $\gamma$ is the minimum color margin desired between the colors of adjacent neurons and $m_{i,j}$ denotes the "margin violations" that we aim to minimize. The interpretation of the first constraint is that, if cell $i$ and cell $j$ are neighbors (i.e., $\boldsymbol{A}_{i,j} = 1$), then the color difference between them, encoded by the term $\|\boldsymbol{R}_i\boldsymbol{X} - \boldsymbol{R}_j\boldsymbol{X}\|_F^2$, should be at least $\gamma$. However, to soften this constraint, we enable some color differences to be less than the desired margin provided that we pay a penalty for violating this margin, encoded by $m_{i,j}$, This constraint drives the optimization towards ensuring that these neighboring cells are distinguishable by color. Otherwise, if they are not neighbors (i.e., $\boldsymbol{A}_{i,j} = 0$), the objective does not pay a price for having similar colors. Of note, $\gamma$ permits discrimination via color intensity as is the case in NeuroPAL where, for example, neighboring cells express reporters with stereotypical intensity levels that distinguish a faint-green neuron from its bright-green neighbor. Concretely, if $\gamma = 0.33$, then neurons with color profiles [0 1 0] and [0 0.66 0], which correspond to bright-green and faint-green, are considered to be sufficiently discriminated from each other. The constraint $\boldsymbol{X} \in \{0,1\}^{r\times c}$ also permits solutions of color assignments to reporters whose expression patterns are simply noted, per cell, in the binary format of "expressed" or "absent" (as is typically the case for the worm Brain Atlas and other community-procured, model-organism databases). The constraint $\boldsymbol{R}\boldsymbol{X}\boldsymbol{1} = \boldsymbol{1}$ ensures that the color we obtain for each cell, $\boldsymbol{R}_i\boldsymbol{X}$, is a bonafide color, such that its color channel values sum to one. The main reason for this constraint is to ensure that all neurons have color intensity in at least some channel so that they can be detected in imaging[1]. The constraint $\boldsymbol{X}^T\boldsymbol{1} \geq \boldsymbol{1}$ ensures that all available colors are utilized. Lastly, $m_{i,j} \geq 0$ is a constraint such that we only penalize positive margin violations and color assignments that exceed the margin requirement are not penalized. For example, if two adjacent neurons have the colorings [1 0 0] (red) and [0 1 0] (blue), such that the color difference exceeds the necessary margin of 0.33, this surplus color difference is not penalized by the model.

The overall program cannot be solved in computationally tractable time if the adjacency matrix $\boldsymbol{A}$ is dense and if $n$ or $c$ is large because in general integer programming is NP-hard. However, it is possible to relax the problem such that we admit a continuous expression of colors. Adding a sparsity regularizer encourages continuous optimization to yield discrete solutions [Donoho, 1995, Donoho, 2006].

$$
\begin{aligned}
\underset{\boldsymbol{X},m}{\text{minimize}} \; & \sum_{i,j} m_{i,j} + \lambda\|\boldsymbol{X}\|_1 \\
\text{s.t. } & \|\boldsymbol{R}_i\boldsymbol{X} - \boldsymbol{R}_j\boldsymbol{X}\|_F^2 \geq \gamma \boldsymbol{A}_{i,j} - m_{i,j} \\
& \boldsymbol{X} \geq 0 \\
& \boldsymbol{R}\boldsymbol{X}\boldsymbol{1} = \boldsymbol{1} \\
& \boldsymbol{X}^T\boldsymbol{1} \geq \boldsymbol{1} \\
& m_{i,j} \geq 0
\end{aligned}
\tag{2}
$$

The optimization problem in (2) is a non-convex problem, since it can be written as a quadratically constrained quadratic program (QCQP) with constraint matrices that are not generally positive semidefinite [Boyd and Vandenberghe, 2004]. An intuitive demonstration of the non-convexity and the existence of multiple optimal solutions of (1) and (2) is that given a valid color assignment to all reporters, we can permute the color order of all reporters in the optimal solution without changing the objective function, yielding another optimal solution with a completely different color swatch. However, the main advantage of (2) over (1) is that (1) must be solved combinatorially while (2) can be optimized to approximate (1) using coordinate descent with multiplicative updates [Arora et al., 2012].

In (2) $\lambda$ controls the amount of sparsity that we desire in the reporter usage matrix $\boldsymbol{X}$. As $\lambda$ is increased, fewer reporters are used to drive color separation. The solution of the above objective in (2) generates a locally-optimal coloring solution of the relaxed problem. However, this solution may serve as only one possible surrogate among many possible solutions of the integer programming objective expressed in (1), since this program is inherently non-convex with the potential for multiple local minima. Beneficially, these local minima can be interpreted as multiple solutions that attain a high-quality color map. Thus, a simple pseudo-combinatorial optimization of (1) with random initializations can be used to generate multiple good solutions that provide alternatives in the event of any unforeseen real-world biological confounds to the obtained solution.

[1] We have also considered a relaxed constraint, $\boldsymbol{R}\boldsymbol{X}\boldsymbol{1} \leq \boldsymbol{1}$, such that the color channels sum to at most one. This option can be enabled in our code if there is a pancellular marker for the set of cells in consideration (in the NeuroPAL case, a panneuronal reporter). In this case, the algorithm might choose to not label a given cell with any colors (i.e., the cell would be dark in the landmark color channels) but would still be detectable in the pancellular marker channel; such "dark" cells would still be distinguishable from their color-labeled neighbors. The benefit of the $\boldsymbol{R}\boldsymbol{X}\boldsymbol{1} = \boldsymbol{1}$ constraint is that it obviates the need for a pancellular marker, using only the landmark colors to distinguish cells.

#### 3 Optimization

The optimization of the objective (2) can be undertaken by multiplicative matrix updates [Arora et al., 2012] that exploit the fact that all the variables involved are non-negative. First, we create an auxiliary variable, $\boldsymbol{Z}$, that denotes the colors of the neurons such that these colors are driven by reporter expression ($\boldsymbol{R}$) multiplied by reporter color tags ($\boldsymbol{X}$), $\boldsymbol{Z} = \boldsymbol{RX}$. We then iteratively update $\boldsymbol{X}$ and $\boldsymbol{Z}$ using multiplicative updates outlined in algorithm 1. Here $\oslash$ and $\odot$ denote elementwise division and multiplication, respectively.

**Algorithm 1** Maximum margin approximate graph coloring

**Input:** $\boldsymbol{A} \in \{0,1\}^{n\times n}$ (Adjacency matrix), $\boldsymbol{R} \in \mathbb{R}_+^{n\times r}$ (reporter rank constraints), $c \in \mathbb{Z}_+$ (number of colors), $\gamma > 0$ (allowable color margin), $\lambda \geq 0$ (sparsity level)
**Initialize:** $\boldsymbol{X} \sim \text{Unif}[0,1]^{r\times c}$, $\boldsymbol{Z} \sim \text{Unif}[0,1]^{n\times c}$
**while** not converged **do**

$$\boldsymbol{Z} \leftarrow \boldsymbol{Z} \odot \frac{\left(\sum_{i,j} A_{i,j}\mathbf{1}\left[(\boldsymbol{z}_i-\boldsymbol{z}_j)(\boldsymbol{z}_i-\boldsymbol{z}_j)^T \leq \gamma\right] A_{i,j}\boldsymbol{z}_i + \boldsymbol{Z}\right)}{\left(\sum_{i,j} A_{i,j}\mathbf{1}\left[(\boldsymbol{z}_i-\boldsymbol{z}_j)(\boldsymbol{z}_i-\boldsymbol{z}_j)^T \leq \gamma\right] A_{i,j}\boldsymbol{z}_j + \boldsymbol{RX}\right)} \text{ (Multiplicative update for auxiliary variable } \boldsymbol{Z})$$

$\boldsymbol{Z} \leftarrow \boldsymbol{Z} \oslash \boldsymbol{Z}\mathbf{1}$ (Column normalization)
$\boldsymbol{X} \leftarrow \boldsymbol{X} \odot \frac{2\boldsymbol{R}^T\boldsymbol{Z}}{2\boldsymbol{R}^T\boldsymbol{RX}+\gamma}$ (Multiplicative update for $\boldsymbol{X}$)
$\boldsymbol{X}^T \leftarrow \boldsymbol{X}^T \oslash \min\{\boldsymbol{X}^T\mathbf{1}, \mathbf{1}\}$ (Column-sum constraint)
$\boldsymbol{Z} \leftarrow \boldsymbol{RX}$ (Update auxiliary variable $\boldsymbol{Z}$)
**end while**
**return** $\boldsymbol{X} \in \mathbb{R}_+^{r\times c}$ (Color assignments)

The multiplicative updates for the main variable $\boldsymbol{X}$ and the auxiliary variable $\boldsymbol{Z}$ are derived by considering the Karesh-Kuhn-Tucker (KKT) conditions of (2) and taking the quotients of the negative component of the derivative update step with the positive component, ensuring a monotonic decrease in the objective function while respecting the non-negativity constraints [Arora et al., 2012]. Further column normalization steps are undertaken to meet the additional constraints that at least one color channel is utilized for each reporter chosen and that the overall color of each neuron is bonafide (color values sum to one).

#### 4 Results

##### 4.1 Approximately optimal coloring examples

Using algorithm 1, we generated simulated NeuroPAL alternatives to test the effects of various real-world limitations on our approximately-optimal-coloring solutions. As such, we sampled a variety of restrictions limiting the number of distinguishable colors (e.g., a restriction present when using 2-photon microscopy), distinguishable color intensities (e.g., a restriction present when using reporters with variable expression levels), and the number of reporters used (e.g., a restriction imposed by organism-specific transgenesis techniques). As mentioned earlier, we used the statistical atlas of neuronal position and variability (**Text S1**), measured using NeuroPAL, for our adjacency matrix $\boldsymbol{A}$. For the list of cell-specific reporters and their expression, $\boldsymbol{R}$, we used one of two sets: a) the "Brain Atlas" list of 1192 reporters (unique = 654)[2] with binary information for each cell's expression (present or absent) and b) a refined "NeuroPAL-tested" list of 97 reporters (unique = 91)[2], tested as candidates for NeuroPAL, wherein we noted stereotypical intensity information for each cell's expression (high, medium-high, medium-low, low, and undetectable). Our approximately optimal-coloring solutions are merely *in silico* simulations and do not capture real-world biological confounds (e.g., potential reporter crosstalk) nor do they capture all real-world NeuroPAL cellular-discrimination techniques (e.g., distinguishing neuronal nuclei via size differences). Nonetheless, many of these real-world settings can be encoded into the algorithmic variables and our method provides a variety of solutions to test in vivo, should any issues arise. For example, the graph adjacency matrix that encodes proximity between neurons can be modified to rule out proximity between neurons that may be spatially close but morphologically very distinct. Thus, these neurons would not need to be distinguished by color.

We plotted a selection of our approximately-optimal NeuroPAL alternatives, under a variety of real-world restrictions (**Figure 1**), to provide a visual approximation of what these simulations might look like *in vivo* (see **Table S7** for the reporter-fluorophore combinations representing each simulation). The approximately-optimal NeuroPAL alternatives using both the NeuroPAL-tested and Brain Atlas reporter sets, simulated under constraints similar to NeuroPAL, show strong visual distinguishability between neighboring neurons (**Figure 1B-C**). These simulations suggest that both theoretical alternatives may prove to be strong competitors to NeuroPAL *in vivo*. Surprisingly, restricting the reporter set to only 10 reporters still generates an approximately-optimal solution that, both visually and in its measure of color violations, suggests it too might prove a close competitor to NeuroPAL *in vivo* (**Figure 1E**). Further restrictions yield solutions that compare less favorably to the aforementioned ones, but that may nonetheless be useful for some constrained applications (**Figure 1D and F-I**).

[2]The uniqueness value of the reporter set indicates its level of redundancy, or lack thereof. Brain Atlas has uniqueness of 654 out of 1192 reporters, indicating that it contains 654 reporters with unique expression patterns. Similarly, 91 out of the 97 NeuroPAL-tested reporters are non-redundant. We used the reduced sets of uniquely expressed reporters, for our analysis, since redundant reporters can simply be interchanged with their duplicates.

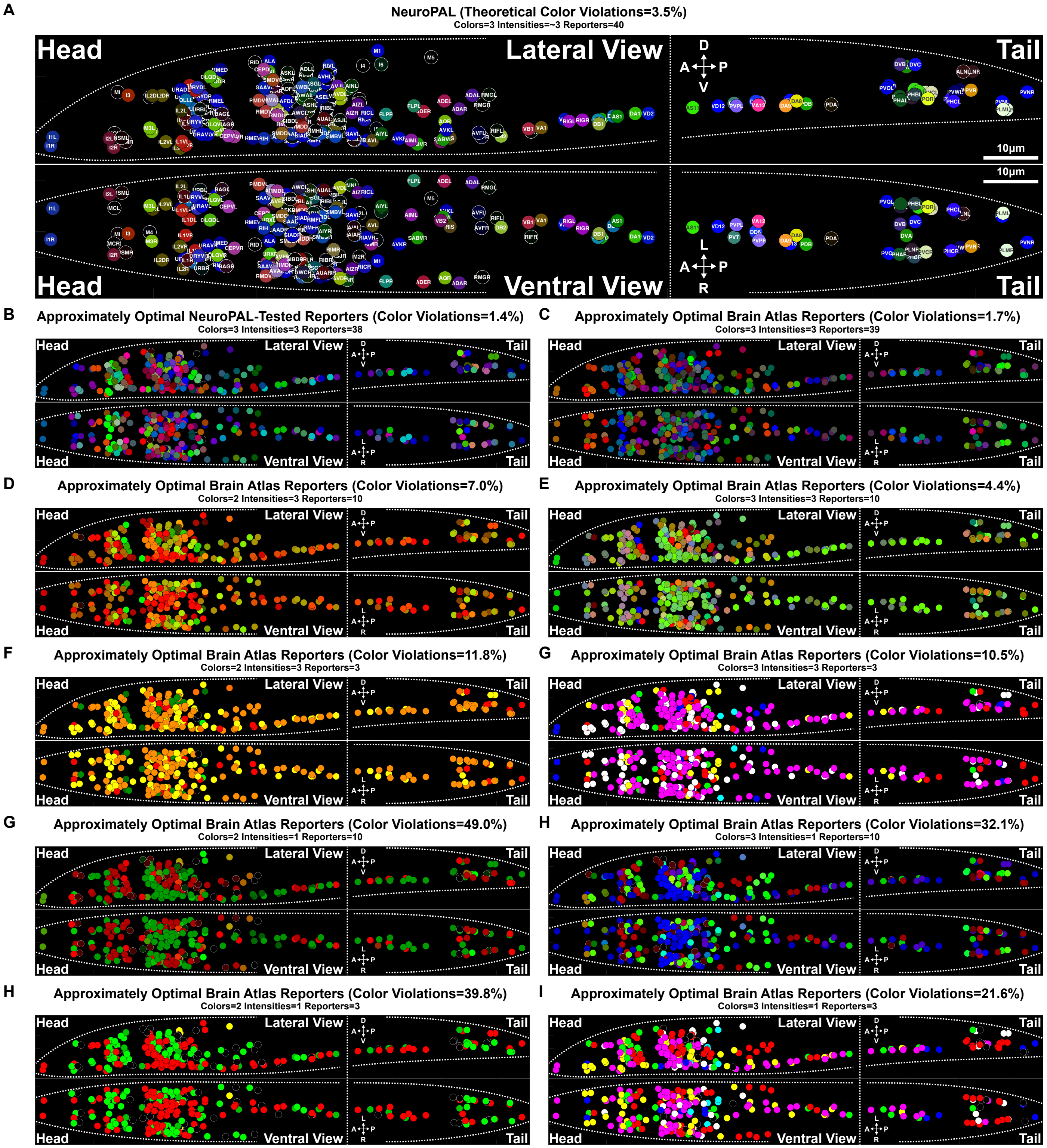


**Figure 1: Approximately-optimal simulated NeuroPAL alternatives.**
See **Table S7** for the reporter-fluorophore combinations representing each simulation in this figure.
(A) A generalization of a NeuroPAL worm, plotted using an atlas of cell positions and colors that were generated from 10 young-adult NeuroPAL worms (**Text S1**), for comparison with the simulated alternatives plotted below. NeuroPAL color violations serve as a rough baseline to evaluate the performance of the approximately-optimal solutions.
(B-C) Two approximately-optimal NeuroPAL alternatives (chosen as those with the minimal color violations, among 10,000 simulations), with algorithmic values approximating those used in constructing NeuroPAL (3 distinguishable colors, with 3 distinguishable intensities, using approximately 40 reporters). Both alternatives exhibit a low percentage of color violations and could prove useful *in vivo* as alternatives to NeuroPAL. (B) A NeuroPAL alternative generated using NeuroPAL-tested reporters, a list of 97 reporters that were tested as candidates for NeuroPAL and whose cell-specific expression levels were recorded (approximated as high, medium-high, medium-low, low, and undetectable expression). (C) A NeuroPAL alternative generated using Brain Atlas, a list of 1192 neuron-specific reporters (curated from WormBase) and whose cell-specific expression patterns are approximated as binary (present or absent). (D-I) Simulated NeuroPAL alternatives (chosen as those with the minimal color violations, among 7,500 simulations), using the Brain Atlas reporters, evaluating real-world limitations that restrict the optimal-coloring solutions to use only 2 or 3 colors, 1 or 3 distinguishable intensities, and 3 or 10 reporters.

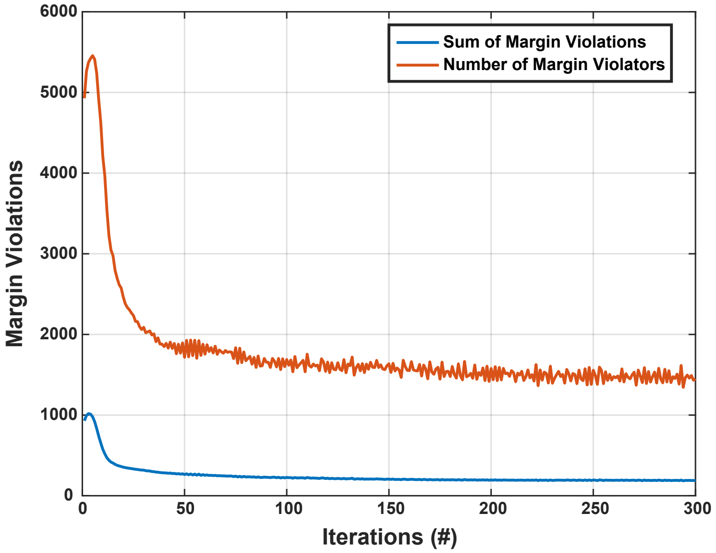


**Figure 2: The optimal-coloring objective function exhibits convergence in margin violations**.
The convergence behavior of the optimal-coloring algorithm 1 exhibits monotonically decreasing margin violations following an initial non-monotonic transient. The initial non-monotonicity is due to random initialization that may not be in the feasible set; in this case, the updates steer the solution towards feasibility, momentarily increasing the margin violations prior to monotonically decreasing until convergence. Here we used an adjacency matrix, $\boldsymbol{A}$, derived from the positional atlas of *C. elegans* neurons by linking any two neurons whose covariance ellipses overlap with 95% probability. $\boldsymbol{R}$ is set to use Brain Atlas as the list of available reporters, $c$=3 colors, $\gamma$=1/3 as our minimum color margin (corresponding to discriminating 3 levels of color intensity), and $\lambda$=100 as our sparsity parameter for reporter usage (roughly corresponding to using 40 reporters). The algorithm reaches a minimal solution, within 100 iterations, which takes about 11 seconds in total.

##### 4.2 Algorithmic convergence

Algorithm 1, that optimizes the objective in (2), is initialized with a random seed and converges to a particular local minimum in a non-convex optimization landscape that may have many minima. The convergence rate of the algorithm can be seen in **Figure 2**. This figure shows that, after an initial transient that serves to locate a feasible point satisfying the constraints, the margin violations are monotonically reduced until convergence. The convergence speed for 235 neurons (the head and tail of the worm, excluding its ventral nerve cord and midbody neurons), 654 Brain Atlas reporters, 3 colors (as used in NeuroPAL), 1/3 as our minimum color margin (corresponding to discriminating 3 levels of color intensity), and 100 as our sparsity parameter for reporter usage (roughly corresponding to using 40 reporters) was about 25 seconds for 300 iterations in total. The experiments were run using MATLAB version 9.7.0.1319299 (R2019b) Update 5. The computer was a MacBook pro laptop (15-inch, 2018), running macOS Mojave version 10.14.6, with 32 GB RAM and 6 Intel Core i9 2.9GHz processors. In general, the algorithm scales quadratically with number of cells and linearly with number of reporters.

##### 4.3 Assessment of optimality

To evaluate approximately optimal coloring solutions, we measure the percentage of their color violations, defined as neighboring neuron pairs which fall below the discrimination margin. It should be noted that the reported margin violation percentages are computed only as a proportion of neighboring neuron pairs and not *all* neuron pairs. For example, if all neurons are colored the same way, all pairs of neurons will exhibit color margin violations at a 100% rate. Conversely, if there are only 10 pairs of neurons that are deemed to be neighbors and of these, only one pair of neurons have colors that are indistinguishable, then the margin violation rate will be 10%. Minimizing the percentage of color violations is equivalent to maximizing the number of neighboring neuron pairs that are distinguishable from each other. We evaluated NeuroPAL, using a statistical atlas of neuron positions and colors generated from 10 young-adult *otIs669* (OH15262) animals (see **Text S1**). NeuroPAL, which comprehensively identifies all neuron types, exhibited a theoretical color violation of 3.5%. Our color violation metric uses several simplifying assumptions and, thus, we examined these assumptions to understand which neuron pairs contributed to the NeuroPAL margin violations. We discovered several categories of margin violations that, on the whole, are not problematic for real-world neuronal distinguishability. We enumerate these categories, providing examples of NeuroPAL color violations for each one, and discuss their impact on modeling optimal-coloring solutions.

Foremost, a subset of *C. elegans* neurons are left-right neighboring pairs of the same type (AIA, AVF, PVP, RIF, RIG, RMF, RMH, and SABV) and, to date, no reporters are known that distinguish the left and right neurons of each pair. The inability to distinguish these left-right neuron pairs does not impact comprehensive neuronal identification since each pair represents a single cell type. As an example, while NeuroPAL distinguishes RIG neurons from other cell types, it cannot distinguish between RIGL and RIGR. Thus our

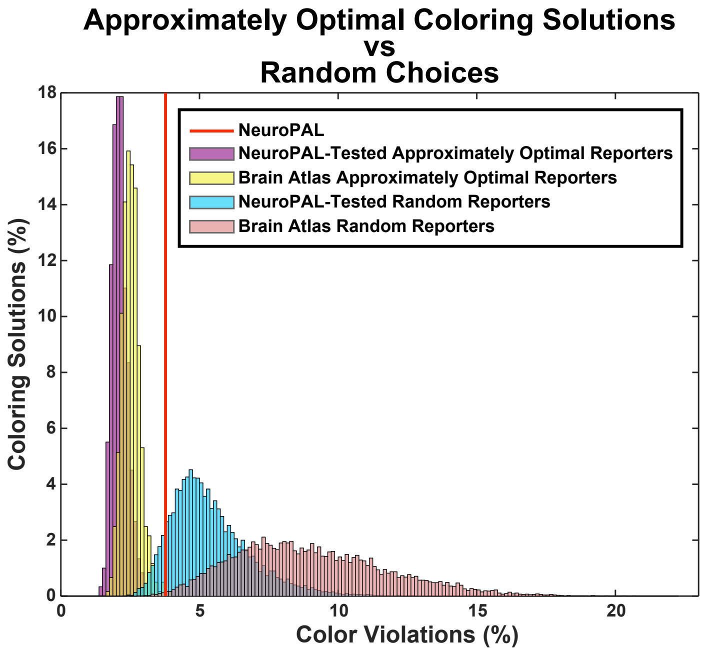


**Figure 3: Approximately optimal coloring versus random reporter-fluorophore combinations.**
Histograms comparing the approximate optimal-coloring algorithm to random reporter-fluorophore combinations. All coloring solutions use NeuroPAL-like parameters (3 colors, 3 distinguishable color intensities, and exactly 40 reporters). The approximately optimal-coloring solutions, using the NeuroPAL-tested and Brain Atlas reporter lists, considerably outperform their randomly-generated counterparts. Even though NeuroPAL usually outperforms randomly-generated reporter-fluorophore combinations, our results suggest a more optimal alternative may exist that provides even greater color distinguishability between neighboring neurons.

metric correctly notes that this pair of neighbors violate the margin of discrimination. This subset of indistinguishable left-right neuron pairs represents a lower bound (0.3%) for optimal coloring and, thus, a minimal percentage of color violations inherent in every solution.

Next, our atlas of neuronal positional variability does not represent pairwise positional relationships. For example, the I3 neuron in the pharynx neighbors the adjacent, but biologically-separate, IL2DL/R neurons in the anterior ganglion. Neurons in different ganglia are restricted by biological structures and are thus distinguishable. The pharynx represents a rigid structure (basal lamina) and neurons therein have even more pairwise positional stereotypy than those in other ganglia. Therefore, although our atlas of positional variability does not encode their positional relationships, neuron types such as I3 and IL2D need not be distinguishably colored in order to be discriminated. Furthermore, even the nearly-adjacent, pharyngeal I1L/R neuronal pair are distinguishable because they have stereotyped positions, embedded in the rigid structure of the pharynx, and are further divided onto opposing sides of a long, thin tube for food intake (the buccal cavity). If desired, the adjacency matrix $A$ can be manually updated to reflect these invariant relationships by eliminating their corresponding entries.

Lastly, our choice of color margin (1/3) is a conservative estimate meant to ensure that neuronal colors are easily visually distinguished. Our conservative choice of color margin not only reflects a measure of visual distinguishability but it also relates to the cell-specific expression levels of the reporters themselves. The NeuroPAL-tested reporters were annotated, for each cell in which they are expressed, as an approximation of five intensity levels (high, medium-high, medium-low, low, and undetectable). As such, we expect our metric to work with at least these levels of discrimination – in practice, we find that an even more refined set of intensities can sometimes be discriminated. However, Brain Atlas reporter expression is merely represented in a binary format (present or absent). *In vivo*, these Brain Atlas reporters will still be distinguishable within some range of intensities but the outcome is far more unpredictable. Thus, we chose a relatively conservative estimate for the color margin to help enforce larger discrimination between neuron colors. As an example, in NeuroPAL, URA is distinguishable from its neighbor URY because, while both neurons express blue (mTagBFP2), only URY expresses red (mNeptune2.5). Nonetheless, the color differences between URA and URY fall below our conservative discrimination margin and thus these pairs of 4-fold symmetric neurons are counted as violating the margin of discrimination. This example points to a further opportunity of interest. Since all microscopy measures of color are encoded as RGB measurements, software may be better poised to discriminate color differences than even human experts. Therefore, the choice of discrimination margin can either be set to reflect the goal of human discrimination or further refined for machine-vision approaches (see **Text S1**).

Importantly, since our color violation metric only approximates real-world neuronal distinguishability, we modeled our optimal-coloring parameters as choices that reflect more conservative ones than those needed for comprehensive neuronal identification. As such, even non-zero color violations can be expected to represent solutions that approach, and potentially achieve, comprehensive neuronal identification. In particular, color violations below those of NeuroPAL (3.5%) may offer alternatives with even greater visual distinguishability and thus improve on this manually-designed transgene. Overall, low values of color violations are indicative of promising approximately-

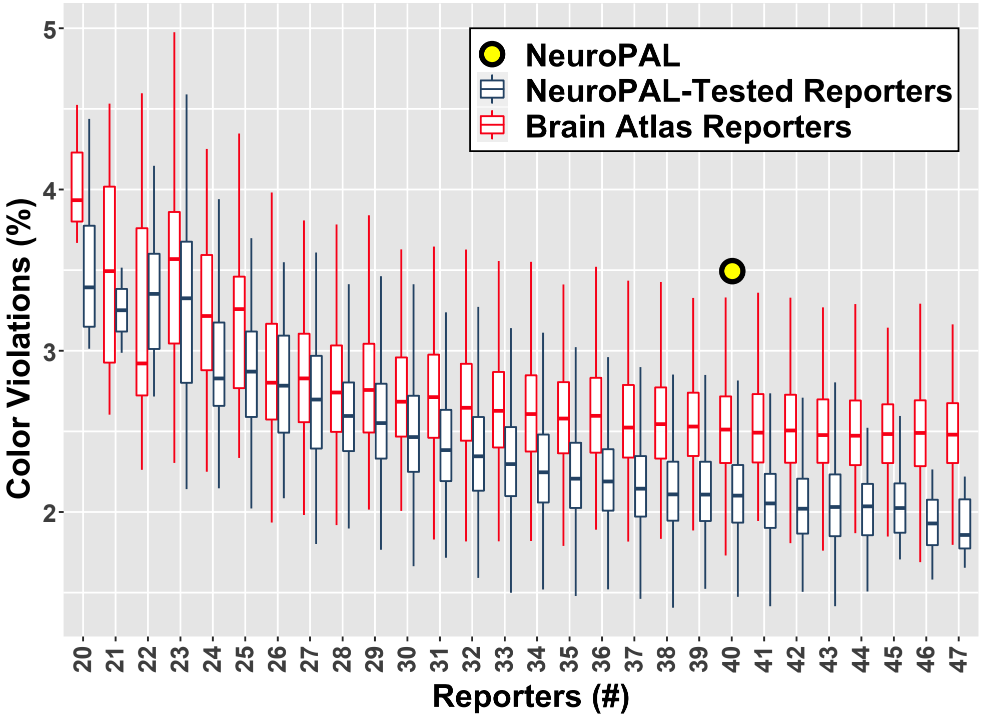


**Figure 4: Approximately optimal coloring results with 40 reporters.**
10,000 approximately-optimal-coloring solutions, generated using the NeuroPAL-tested and Brain Atlas reporter lists, with parameters similar to NeuroPAL (3 colors, distinguishable at 3 intensities, using roughly 40 reporters). NeuroPAL performance is marked by a yellow dot. Our results suggest that many of these simulated NeuroPAL alternatives might perform favorably *in vivo*. Furthermore, we can observe a general downward trend in the mean color-margin violations as more NeuroPAL-tested and Brain Atlas reporters are used. The color margin violation reduction reaches a diminishing rate at about 40 reporters.

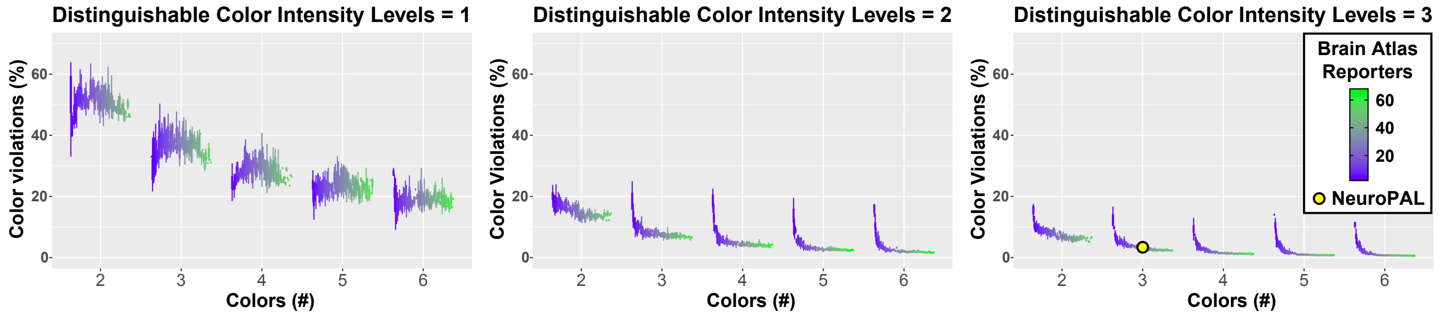


**Figure 5: Approximately optimal coloring results under real-world constraints.**
7,500 approximately-optimal-coloring solutions, generated using the Brain Atlas reporter list, that sample a wider range of parameters representing other possible real-world scenarios: 2-6 colors, distinguishable at 1-3 intensities, using 2-71 reporters. The distribution of approximate solutions is displayed by grouping them into several hierarchical subcategories. We grouped solutions by their different intensity levels, then by the number of colors that they used, and lastly by the number of reporters that they used. Our results indicate that at least two distinguishable color intensities and roughly five or more reporters strongly benefit minimal color violations, but low color violations can still be achieved below these limits. In general, we can confirm an algorithmically-expected trend that more colors, more intensity levels, and most importantly, more reporters, enable us to get a lower color-margin violation. However, the trend displays diminishing returns due to geometrical constraints on neuron positions and reporter redundancy.

optimal-coloring solutions. We find that, if one removes NeuroPAL margin violations contributed by the categories discussed above, the NeuroPAL color violations drop to zero.

Running the optimal-coloring algorithm 1 (with random seeds) leads to a variety of different solutions one can test *in vivo*. We generated 10,000 approximately-optimal-coloring solutions, using both our Brain-Atlas and NeuroPAL-tested reporters, enforcing algorithmic values that best reflect NeuroPAL (3 colors, distinguishable at 3 intensities, using roughly 40 reporters – the 41st reporter is panneuronal and is implicit in every solution), then compared these with NeuroPAL color violations. From these solutions we selected all samples that use exactly 40 reporters (as in NeuroPAL), leaving us with approximately 600 samples each, generated from the NeuroPAL-tested and Brain Atlas reporters (the two leftmost distributions in **Figure 3**). We compared these to randomly-generated reporter-fluorophore combinations

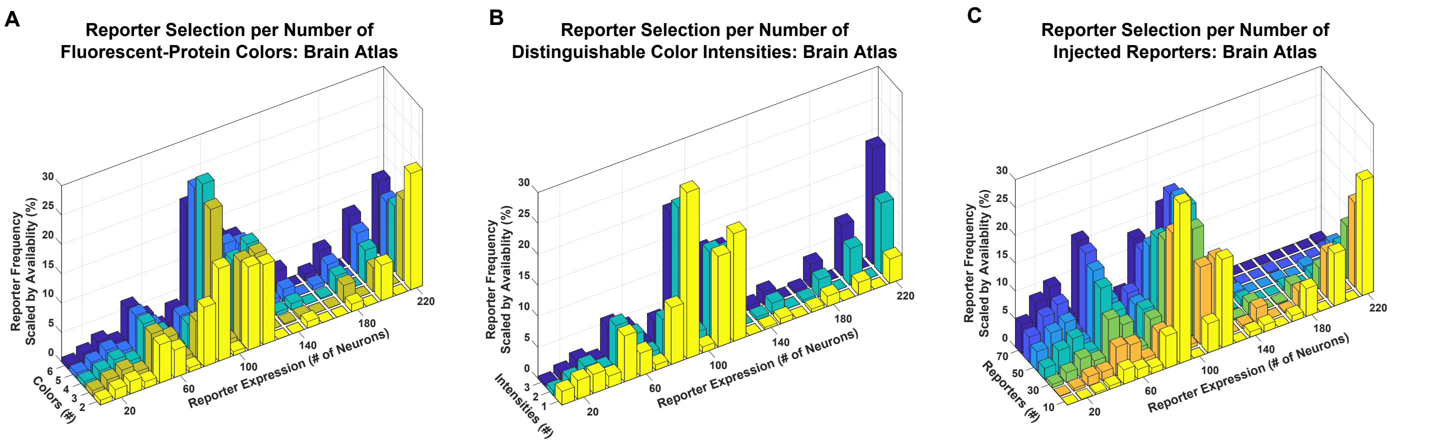


**Figure 6: Approximately optimal reporter selection under real-world constraints.**
The reporters chosen by the algorithm are scaled by their representation in the Brain Atlas reporter list. For each range of cellular expression (e.g., targeting from 1-10, 11-20, and 21-30 neurons), the total number of reporters used among all algorithmic solutions that are within this range are divided by the total number of reporters available in Brain Atlas that match this same range of cellular expression. These are displayed as their frequency in algorithmic solutions scaled by their availability. Narrowly expressed reporters (e.g., those targeted to 10 or fewer cells) are abundantly represented and thus, the algorithm choosing them is less meaningful than it choosing broadly expressed reporters (e.g., those expressed in nearly 220 cells) which have far less representation.
(A) Limiting the number of colors available did not lead to any salient trend in the expression of the reporters one should use, equally favoring roughly narrow to broad cellular expression patterns.
(B) Limiting the number of distinguishable color intensities, from three to one, led to a clear trend favoring the use of reporters with narrow (targeted) expression patterns in place of those with broad expression patterns. This suggests that under this restriction, the algorithm favors a strategy that targets coloring to specific neurons in order to collectively distinguish neighboring cell pairs.
(C) Limiting the number of reporters used favors those with broad expression patterns. Under this constraint, the algorithm may use broad coloring to achieve at least coarse discriminability, over many cells, rather than targeting just a subset of cells.

that were formulated by choosing 40 random reporters from either the NeuroPAL-tested or Brain Atlas lists and combining each of these reporters with anywhere from 1-3 randomly chosen colors (the two rightmost distributions in **Figure 3**). When restricted to exactly 40 reporters, our approximately-optimal-coloring solutions using the Brain-Atlas and NeuroPAL-tested reporter lists exhibited color violations ranging from 1.7-3.9% and 1.5-3.0%, respectively. In comparison, our randomly-generated fluorophore combinations using the Brain-Atlas and NeuroPAL-tested reporter lists exhibited color violations ranging from 2.4-22.4% and 2.3-15%, respectively. These results indicate that our algorithm consistently generates solutions that outperform these random combinations. NeuroPAL outperforms these random combinations but our results suggest that a more optimal NeuroPAL may exist that provides even greater color distinguishability between neighboring neurons.

Our algorithmically-generated NeuroPAL alternatives, which use exactly 40 reporters, exhibited competitive color violation scores, suggesting that many of these solutions might perform favorably *in vivo*. Thus, to assess algorithmic performance over a broader range in the number of reporters permitted we further evaluated all 10,000 samples, generated for both the NeuroPAL-tested and Brain Atlas reporter lists (**Figure 4**). Brain-Atlas and NeuroPAL-tested reporter results still exhibited low color violations, ranging from 1.7-5.3% and 1.4-4.6%, respectively, suggesting that even with far fewer reporters, our algorithm generates competitive NeuroPAL alternatives.

Given these results, we expanded our testing to accommodate other possible real-world scenarios ranging from 2-6 distinguishable colors, with 1-3 distinguishable intensities, using 2-71 reporters chosen from the Brain Atlas and generated 7,500 approximately optimal-coloring solutions (**Figure 5**). As expected, these restrictions generated solutions with color violations ranging from 0.5%, suggesting a near-optimal coloring solution (with a theoretical lower bound of 0.3%), to 79%, suggesting a non-optimal coloring solution. Our results indicate that, for worm neurons, at least two distinguishable color intensities (high and low) and roughly five or more reporters strongly benefit minimal color violations (**Figure 1B-E**). Nonetheless, low color violations could still be achieved below these limits.

##### 4.4 Approximately optimal reporter selection

We investigated the tradeoffs, present in the optimal-coloring algorithm solutions, when constrained by the aforementioned real-world scenarios (**Figure 6**). We found that limiting the number of colors available did not lead to any salient trend in the expression of the reporters one should use, equally favoring roughly narrow to broad cellular expression patterns (**Figure 6A**). However, limiting the number of distinguishable color intensities, from three to one, led to a clear trend favoring the use of reporters with narrow (targeted) expression patterns in place of those with broad expression patterns (**Figure 6B**). This suggests that under this restriction, the algorithm favors a strategy that targets coloring to specific neurons in order to collectively distinguish neighboring cell pairs. When limiting the number of reporters used, we found the opposite trend (**Figure 6C**). Here, when fewer reporters were available the algorithm favored those with broad expression patterns, to achieve at least coarse discriminability, over many cells, rather than targeting just a subset of cells.

##### 4.5 Software

We developed a software script in MATLAB to generate approximately optimal-coloring solutions for any collection of cells in any organism, available at `https://github.com/Eviatar/Optimal_Coloring`.

#### 5 Conclusion

The algorithmic examples above illustrate how the NeuroPAL technique could be extended to offer multicolor landmarking solutions for any collection of cells, in any model organism, using similar databases of reporter expression (e.g., using Flybase for fly, ZFIN for zebrafish, and MGI for mouse) [Eppig, 2017, Thurmond et al., 2019, Ruzicka et al., 2019, Harris et al., 2020]. Our adjacency matrix ($\boldsymbol{A}$) for *C. elegans* neurons used a precise measure of neuron positions built using 10 NeuroPAL animals. Such precision is rarely available for other cell collections but a considerable number of fluorescent images exist for the reporters listed in model organism databases [Chiang et al., 2011, Jenett et al., 2012, Manning et al., 2012, Li et al., 2014, Oh et al., 2014, Randlett et al., 2015, Costa et al., 2016, Erö et al., 2018, Kunst et al., 2019]. Furthermore, recent work has augmented such images with electron micrograph reconstructions that cover nearly the full cellular contents of these model organism brains [Kasthuri et al., 2015, Ryan et al., 2016, Hildebrand et al., 2017, Zheng et al., 2018]. These datasets can inform either a simple, manually-annotated adjacency matrix that delineates which cell types must be distinguishable or they can inform more complex models that use more refined estimates to represent cell adjacency. The worm nervous system is small enough to permit a single strain, such as NeuroPAL, that identifies every neuron therein. Most other model organism brains are much larger and, as such, a single strain solution cannot identifiably color every neuron. In these cases, a more reasonable strategy is to develop a set of application-specific strains that together identify a sufficient collection of neurons so as to analyze the neural circuits of interest. As we show here our algorithm generates approximately-optimal-coloring solutions that can be beneficial despite a variety of real-world restrictions that limit the number of distinguishable colors available, the distinguishable intensity levels of these colors, and/or the number of reporters permitted by transgenesis techniques. Beyond this, the NeuroPAL technique of multicolor cell identification need not be restricted to neurons and the proposed algorithmic approach here can be extended to any type of cell in any model organism. We envision that our algorithm will have broad utility in designing color maps for cell identification.
